# Cohesin forms fountains at active enhancers in *C. elegans*

**DOI:** 10.1101/2023.07.14.549011

**Authors:** Bolaji N. Lüthi, Jennifer I. Semple, Anja Haemmerli, Saurabh Thapliyal, Kalyan Ghadage, Klement Stojanovski, Dario D’Asaro, Moushumi Das, Nick Gilbert, Dominique A. Glauser, Benjamin Towbin, Daniel Jost, Peter Meister

## Abstract

Transcriptional enhancers must locate their target genes with both precision and efficiency. In mammals, this specificity is facilitated by topologically associated domains (TADs), which restrict the enhancer search space through three-dimensional genome organization. In contrast, the nematode genome lacks such TAD-based segmentation despite harboring over 30’000 sequences with chromatin signature characteristic of enhancers, thereby raising the question of how enhancer-promoter specificity is achieved. Using high-resolution Hi-C in *C. elegans*, we identify distinct 3D chromatin structures surrounding active enhancers, which we term fountains. These structures span 38 kb in average, are unique to active enhancers, and are enriched for the major somatic cohesin complex. Fountains collapse upon *in vivo* cohesin cleavage, indicating their cohesin dependency. Notably, fountains accumulate topological stress, as evidenced by the enrichment of topoisomerases and the psoralen-binding signature of negatively-supercoiled DNA. Functionally, fountain disassembly correlates with transcriptional upregulation of active enhancer-proximal genes, suggesting that fountains act as spatial repressors of enhancer activity. This repression is particularly pronounced for neuronal genes, including the *skn-1/Nrf* gene, which becomes upregulated, switches isoform and transcription start site upon cohesin loss in a pair of head neurons. Behaviorally, cohesin cleavage alters nematode movement and foraging behavior, linking enhancer-driven transcriptional changes to neural circuit function and organismal phenotypes, reminiscent of pathologies caused by cohesin mutations in humans. Together, our findings uncover fountains as a novel 3D chromatin feature that modulates enhancer activity in a TAD-less genome, establishing a mechanistic link between genome architecture, gene regulation and behavior.

## Introduction

Gene transcription is tightly regulated by a combination of promoter-proximal elements and more distant enhancer sequences. Enhancers increase the transcription of target genes by recruiting sequence-specific transcription factors, which in turn attract additional proteins to facilitate transcription. Notably, enhancers are capable of activating transcription independently of their location relative to their target gene, and can act at large distances from the promoter, up to several tens of kilobases in mammals^1^.

Recent chromosome conformation capture studies in mammals have shown that enhancer and target promoter are often located in the same megabase-sized three-dimensional domain known as a topologically associated domain (TAD) while TAD segmentation regulates promoter/enhancer contacts and the downstream transcriptional regulation^2–4^. The formation of TADs is the result of chromatin looping, whereby cohesin extrudes chromatin until it reaches sequence elements bound by the DNA-binding protein CTCF (^for^ ^review^ ^5^).

In *C. elegans,* two different studies identified between 19’000 and 30’000 sequences with enhancer-type chromatin features (Fig. 1ab for L3 stage enhancers^6,7^). These sequences exhibit an open chromatin structure (ATAC-seq peaks), characteristic histone marks (high H3K4 monomethylation coupled to low H3K4 trimethylation), short-stretch bidirectional transcription, and enrichment for initiator sequence element (Inr)^6,7^. A limited number of these sequences were individually tested for their capacity to activate a minimal promoter driving GFP independently of their location relative to the promoter. This confirmed that these sequences are *bona fide* enhancers leading to cell- and developmental stage-specific expression of the transgene. It is currently unclear whether and how the activity of these enhancers is limited to their target genes, as unlike mammals, nematodes do not harbor megabase-sized TADs on their autosomes, although smaller, kilobase-sized compartments similar to A and B compartments can be identified in Hi-C data^8^. Additionally, no CTCF homolog or functional homolog has been identified in the *C. elegans* genome, and no nematode boundary elements have been described to date. Moreover, the primary long range loop extruder in nematodes is not cohesin as in mammals, but condensin I^9^. Taken together, these observations suggest that cohesin function is largely divergent in nematodes compared to mammals and that an alternative mechanism to TADs might regulate enhancer-promoter contacts and the ensuing gene expression. Hereafter, we show that active nematode enhancers are located at the tip of small-scale 3D structures, which we call fountains and similar to described jets^10^, plumes or flares^11–13^, created by cohesin activity. Cohesin^COH-1^ cleavage leads to upregulation of active enhancer- and fountain-proximal genes, in particular genes expressed in neurons. Strikingly, cohesin^COH-1^ cleavage leads to a breadth of behavioral changes, linking 3D genome organization by cohesin, neuronal gene expression and nervous system function in animal behavior.

**Figure 1.**
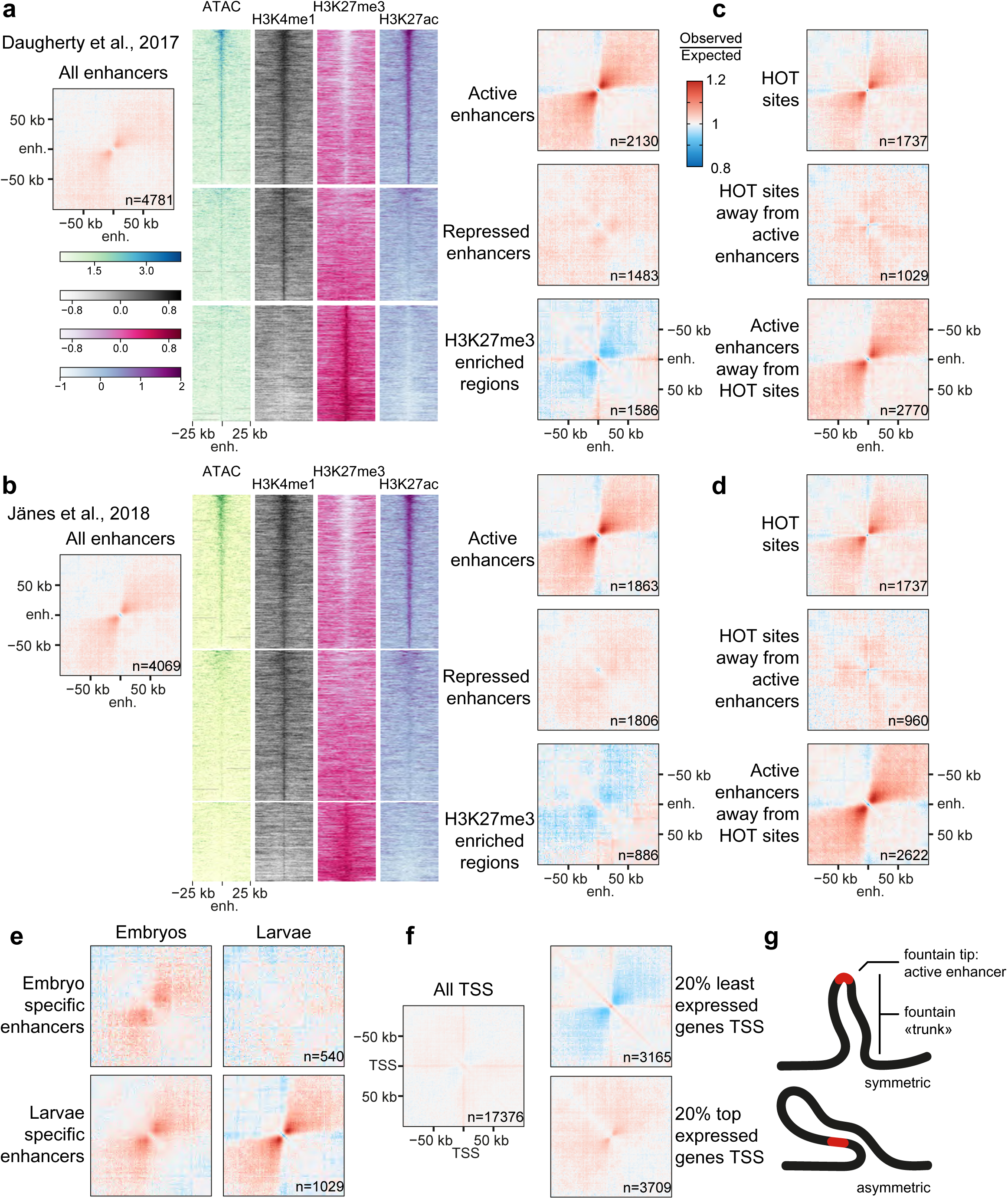
Active enhancers colocalize with loose 3D structures or fountains. **a.** Average contact frequency maps centered on annotated enhancers from Daugherty et al.^7^, highlighting the formation of 3D structures. Further segmentation into active or repressed enhancers and H3K27me3-covered regions highlight the specificity of fountains for active enhancers. **b.** Average contact frequency maps centered on annotated putative enhancers (all stages) from Jänes et al.^6^, further segmented using histone marks as in Daugherty et al. **c,d.** Highly Occupied Target (HOT) sites are not forming fountains. Average contact frequency maps for all HOT sites, for HOT sites located more than 6 kb away from active enhancers and for active enhancers located more than 6 kb away from HOT sites, using active enhancers from Daugherty et al. (c) or Jänes et al. (d). **e.** Fountains for active enhancers are specific for their developmental stage. Average contact maps for embryo-specific and L3 larvae-specific ATAC-seq peaks (ATAC-seq peaks in the top 3 deciles at the considered stage and in the bottom 3 deciles at the other stage). **f.** Transcription start sites (TSS) do not colocalize with fountains. Average contact frequency maps centered on all TSS, and TSS segmented by gene expression. Small fountains present in the most expressed 20 % genes are most likely due to active enhancers located in the vicinity of the TSS. **g.** Cartoon of a typical fountain, with the active enhancer located at the tip of the fountain structure, both a symmetric and asymmetric fountain are shown.

## Results

### Active enhancer loci correlate with 3D fountains

To assess whether enhancers would colocalize with specific three-dimensional genomic features, we generated average chromatin conformation capture contact maps for third larval stage animals centered on enhancer sequences^6,7^. These contact maps revealed a clear and distinct enlarged second diagonal perpendicular to the main Hi-C diagonal, extending several kilobases from the enhancers (Fig. 1ab). These increased contact probabilities between the enhancers and neighboring sequences are indicative of the formation of a loose loop–like structure centered on the enhancer sequences, resulting in the partial alignment of two branches of the loop either side of the enhancers (Fig. 1g). Hereafter, we call these structures fountains, in line with other manuscripts describing these structures^14,15^.

Enhancers were further characterized in one of the studies above using ChromHMM genome segmentation^7^. This division classifies enhancers into active enhancers covered with H3K4me1 and H3K27ac, repressed enhancers with H3K4me1, weak H3K27me3, and no H3K27ac, and H3K27me3-enriched regions covered with H3K27me3, deposited by the PRC2 Polycomb complex (Fig. 1a). When we averaged the contact maps of the different enhancer types and the H3K27me3-enriched regions, we observed that active enhancers were associated with fountains, whereas repressed enhancers did not show any fountains, and H3K27me3-covered regions showed a decrease of expected contacts in the second diagonal, as well as a cross-like high contact probability feature, suggesting that H3K27me3 regions cluster together *in vivo* (Fig. 1a). Similarly, when we segmented the second enhancer dataset^6^ based on the same ChromHMM chromatin segmentation^7^, we made almost identical observations: active enhancers were associated with fountains, while either repressed enhancers or trimethylated H3K27-covered regions were not (Fig. 1b).

In previous ChIP studies, a set of genomic regions bound by multiple transcription factors were identified, named Highly Occupancy Target (HOT) sites^16,17^. As enhancers are binding sites for transcription factors, HOT sites and enhancers often co-occur on nearby loci. To investigate whether HOT sites, active enhancers, or both types of sequence elements colocalize with fountains, we created average contact maps centered on HOT sites that either overlapped with active enhancers or were distinct from them. Our results indicate that HOT sites only correlated with fountains (Fig. 1cd) when these overlapped with active enhancers. In contrast, HOT sites not overlapping with active enhancers did not create fountains, while active enhancers not overlapping with HOT sites did (Fig. 1cd).

We next wondered whether fountains would be stage-specific, by selecting enhancers active in embryos and inactive in the third larval stage, or conversely. We calculated average chromatin conformation contact maps for these different enhancer sets at different stages using our Hi-C data of third larval stage and publicly available Hi-C data performed with the same protocol in embryos^18^. For both enhancer sets, we observed clear fountains in their respective developmental stages (Fig. 1e). In contrast, no fountains could be observed for enhancers active only in embryos using larvae Hi-C maps, and larvae-specific enhancers created smaller and weaker fountains in embryonic Hi-C data (Fig. 1e). The latter might be due to the fact that mixed stage embryos were used to perform Hi-C, in which a variable proportion of animals are already in late developmental stages during which some larval enhancers are already active. We conclude that fountain formation is stage-specific, in agreement with the activity of the enhancers.

Finally, we explored whether fountains were unique to enhancers or if they were associated with other open chromatin regions such as active transcription start sites (TSS). The nematode genome is highly compact and enhancers are located only a couple kilobases away from their putative target promoters^6,7^. To examine TSS-specific 3D structures, we calculated the average contact maps of the bottom or top 20% of all expressed genes ranked by expression level. We found no evidence of fountain formation in the bottom 20%, while the 20% most expressed genes showed very limited fountain formation (Fig. 1f), vastly smaller than the fountains observed at active enhancers (Fig. 1ab). In summary, our data shows that fountain formation is a feature specific to active enhancers and not selectively associated with transcribed genes or HOT sites. We envision fountains as loose loop-like structures with the active enhancer sitting at the tip of the loop (Fig. 1g).

### Enhancers are binding sites for cohesin^COH-1^

Recent studies have described similar structures orthogonal to the Hi-C diagonal in bacteria^19^, *C. elegans*^14^, fungi^13^, T cells^10^ and zebrafish sperm^11^ as well as during zygotic genome activation in zebrafish^15^. These structures are thought to be formed through bidirectional loop extrusion by SMC complexes repeatedly loaded at defined loci (in our case, the active enhancers). Nematodes express five different SMC complexes in the soma: canonical condensin I and II, an X-chromosome specific condensin I variant, and two variants of cohesin, differing by their kleisin subunit, SCC-1 or COH-1. We previously demonstrated that condensin I performs long-range loop extrusion during interphase (>100 kb), while condensin II has no interphasic function^9^. Cohesin^SCC-1^ is involved in sister chromatid cohesion during mitosis and exclusively expressed in dividing cells^20^. Early immunofluorescence studies showed that COH-1 is expressed in all cells, suggesting cohesin^COH-1^ is the major interphasic cohesin. In our previous study, we showed that the common SMC-1 subunit of cohesin is expressed ubiquitously at high levels^9^ (Fig. 2a), suggesting cohesin^COH-1^ is present in most, if not all cells. Direct quantification of COH-1 and SCC-1 abundance using identical tags on the two kleisin subunits showed that in entire animals, COH-1 is 6 times more abundant than SCC-1^9^. To determine whether enhancer sequences were enriched for cohesin^COH-1^, we used modENCODE ChIP-seq data^17^. We observed that cohesin^COH-1^ is specifically enriched in a broad region centered on active enhancers, extending several kilobases away from the enhancer itself (Fig. 2b, upper part, red line and heatmap). In contrast, repressed enhancers had only a small enrichment limited to the enhancer sequences (green line). Cohesin^COH-1^ was slightly depleted on H3K27me3-covered regions compared to neighboring sequences (blue line). To further investigate whether cohesin^COH-1^ enrichment on active enhancers correlated with the size of the fountains, we classified active enhancers into 5 classes based on their cohesin^COH-1^ enrichment and created average contact frequency maps for each class. Our analysis revealed that active enhancers with high cohesin^COH-1^ enrichment generated large fountains extending several tens of kilobases away from the active enhancer locus (Fig. 2b, right side), while active enhancers of the second-to-lowest quintile formed very small fountains and active enhancers of the lowest cohesin^COH-1^ ChIP-seq enrichment quintile did not correlate with fountains.

**Figure 2.**
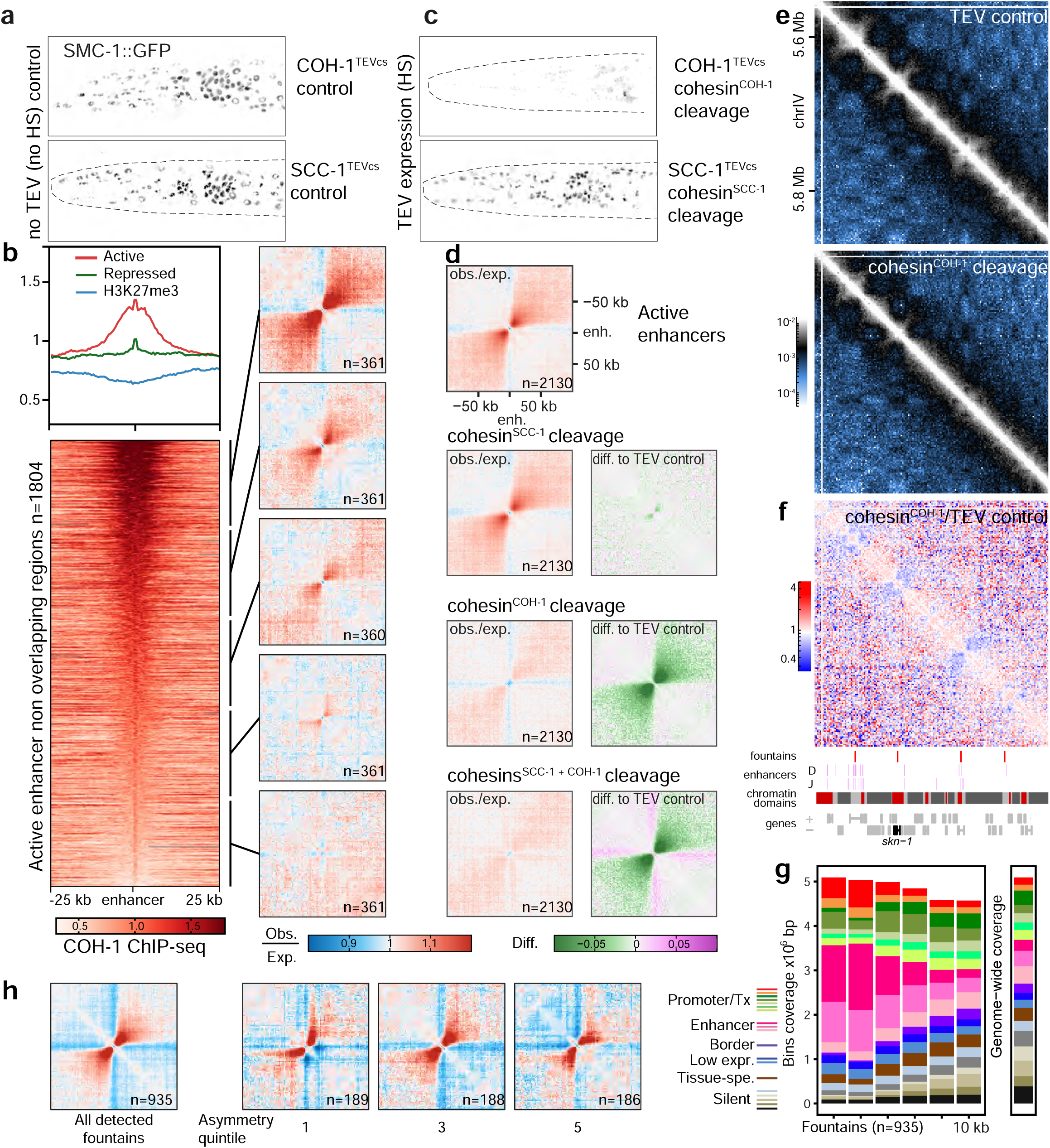
Cohesin^COH-1^ creates fountains at active enhancers. **a.** Fluorescence signal in the head of third larval stage control animals expressing SMC-1::GFP without TEV expression. The dark patch in each nucleus corresponds to the nucleolus. Strains in which TEV cut sites have been inserted in either of the two alternative cohesin kleisins COH-1 and SCC-1 are shown. **b.** (top left) Average COH-1 ChIP-seq profiles in young adult animals at active (red) or repressed (green) enhancers and H3K27me3-enriched regions (blue). (bottom left) Heatmap of COH-1 enrichment on active enhancer regions, sorted by COH-1 ChIP-seq enrichment. (left) Average contact frequency maps centered on enhancers segmented according to COH-1 ChIP-seq abundance. **c.** SMC-1::GFP signal as in a upon cleavage of COH-1 and SCC-1. **d.** Cleavage of cohesin^COH-1^ but not cohesin^SCC-1^ leads to disappearance of active enhancer fountains. Average contact frequency maps, centered on active enhancers upon cleavage of COH-1 and/or SCC-1. The difference between the kleisin cleavages and TEV control is shown on the right with a green/magenta color scale. **e.** Contact frequency map in TEV control animals (top) and upon cohesin^COH-1^ cleavage (bottom) in a section of chromosome IV. **f.** (top) Ratio contact frequency map between cohesin^COH-1^ cleavage and TEV control, highlighting the disappearance of the fountains (blue, negative values). (bottom) Tracks showing detected fountains, active enhancers (Daugherty et al.^7^: D; or Jaenes et al.^6^: J), chromatin domains^23^ and location of genes (bottom). **g.** Autosomal chromatin state coverage of detected fountain bins (left) and their adjacent bins, sorted by distance to the detected fountains. Autosome-wide coverage is shown on the right side. Chromatin states from ^23^. **h.** Average contact frequency maps centered on detected fountains for all fountains and for top, middle and bottom asymmetry quintiles of all fountains, based on the asymmetry score described in the methods section.

To directly investigate the role of cohesin^COH-1^, we made use of our previously described inducible cleavage system for the cohesin kleisins COH-1 and/or SCC-1^21^. When TEV protease expression was induced during the first larval stage (three hours after diapause exit), cleavage of COH-1 (89% cleaved^9^), but not of SCC-1 caused the disappearance of most GFP-tagged SMC-1^9^ (Fig. 2c, quantified in ^9^), indicating that COH-1 cleavage leads to the degradation of the entire cohesin^COH-1^ complex and confirming that cohesin^COH-1^ is the major cohesin isoform. We calculated average contact maps centered on active enhancers in control animals and upon cleavage of either or both cohesin kleisins. Cleavage of SCC-1 only marginally modified contact maps, in contrast to the cleavage of COH-1, in which fountains were almost completely absent (Fig. 2de). Similar results were obtained upon simultaneous cleavage of COH-1 and SCC-1 (Fig. 2d). Cohesin^COH-1^ is thereby necessary for the maintenance of fountains at active enhancers. Given the loop extrusion activity of cohesin^22^, the most likely model for the formation of the fountains is the specific loading of the cohesin complex at active enhancers followed by bidirectional loop extrusion randomly stopping on either side, as suggested by polymer modeling^15^.

### Cohesin-dependent fountains colocalize with active enhancers

The effects of cohesin^COH-1^ cleavage can be visualized by creating contact-ratio maps between cleaved and control contact maps (Fig. 2e, ratio in f). We used these ratio maps to locate fountains, as changes in contact frequency are most likely direct effects of cohesin^COH-1^ cleavage. We filtered the diagonal of the ratio maps with a model-based kernel similar to the average fountain shape observed around enhancers, leading to a fountain similarity score for each locus along the genome (Fig. S1, Methods section and Supplementary File 1). Using this approach, we identified 935 fountains in the entire genome, whose tips are the local optima of the similarity score with significant prominence (“fountain score”; Supplementary Files 2 and 3 for the location and a gallery of all detected fountains). For each fountain, we measured its length, corresponding to the typical extension of the fountain around the tip (*i.e.* the typical loop size extruded by cohesins loaded at the tip), and its (a)symmetry, corresponding to possible deformations in fountain shapes along 5’ or 3’ sides of the tip (*i.e.* asymmetries in cohesin progression around the tip; see Methods and Fig. S1a-f for details and validation of the approach, as well as a comparison with fontanka^15^). Using this method, we determined that the size of 80% of the fountains lies between 22 and 78 kb, with a median fountain length of 38 kb, but some larger fountains could still be detected at 150 kb (Fig. S1j). Notably, while most fountains were symmetric (Fig. S1g), a significant number of fountains were asymmetric, with more contacts on one side than on the other, implying that loop extrusion by cohesin^COH-1^ might be directionally biased in those cases (Fig. 2h). Asymmetry correlated weakly with the abundance of enhancer type chromatin states, enriched on the side where the fountains were longest (Fig. S1i).

Regarding their chromosomal location, fountains did not show any preference, neither for the center nor the arms of chromosomes (Fig. S2a-d), yet they were slightly depleted on chromosome IV and V compared to chromosomes I to III. We did not observe any particular preference for fountain location relative to the boundaries of the X chromosome TADs in dosage-compensated hermaphrodite animals (Fig. S2e). We next compared the location of fountains with ChIP-seq data. It is important to notice here that the resolution of Hi-C data (2 kb) is orders of magnitudes lower than ChIP-seq data. We found that COH-1 was enriched at fountains and positively correlated with fountain score (R=0.52, Fig. S1k). Chromatin states at fountain tips as well as those of the neighboring bins were enriched for enhancer states compared to genome-wide coverage (Fig. 2g), further supporting the notion that fountains are related to enhancers. When we compared the location of detected fountains relative to the different types of enhancers, we found that fountain tips are significantly closer to active enhancers than to repressed ones or H3K27me3-covered regions (Fig. S3ab). Similarly, fountain tips and their +/- 2 kb flanking regions more often contained multiple active enhancers when compared to equally sized control regions located at equal distance between fountain tips, with up to 6 or 11 active enhancers located at the fountain tips and the two neighboring bins, depending on the enhancer mapping study^6,7^. In contrast, repressed enhancers or H3K27me3-covered regions were not enriched in fountain tips and their flanking regions (Fig. S3cd). To further test if fountains preferentially form at clustered active enhancers, we divided the whole genome into 6 kb bins and counted the number of active enhancers overlapping with each bin. While fountains represent only 5.6% of all genomic 6 kb bins, fountain tip bins were more likely to overlap with 6 kb genomic bins encompassing larger numbers of active enhancers. Accordingly, fountain tip bins are vastly overrepresented in genomic bins clustering three or more active enhancers (Fig. S3ef). Since we did see many fountains at bins with more than one enhancer, it seems likely that fountain strength, and therefore detection, improves with the number of enhancers clustered together. Indeed we saw a clear correlation between the number of active enhancers per fountain bin and the fountain prominence score produced by our detection algorithm (Fig. S3gh). Together, we demonstrated that fountains identified on differential Hi-C maps and dependent on cohesin^COH-1^ integrity are primarily associated with active enhancers, while their prominence correlates with the number of clustered enhancers.

A previous study segmented the nematode genome based on histone marks and ATAC-seq data into active, regulated and border domains^23^. We therefore asked whether fountain tips would preferentially colocalize with one type of chromatin domains. Indeed, active domains were enriched at fountain tips, as well as on either side of them (Fig. S4a). Conversely, fountains located in active domains were significantly more prominent than fountains located in border or regulated domains (Fig. S4b). These features, although identified genome-wide, were also apparent when considering individual loci (Fig. 2f, S4c, S11a).

In summary, the genome-wide identification of the locations of cohesin^COH-1^-dependent fountains on Hi-C maps revealed a robust correlation between fountains and active enhancers, as well as COH-1 enrichment. Furthermore, there is a slight correlation between fountain asymmetry and unequal transcriptional activity and/or enhancer chromatin state between the two branches of the loop that emanate from the fountain tips.

### Active enhancers and fountains are bound by topoisomerases

To further understand fountain formation, we examined published ChIP-seq data at enhancer sequences acquired at the same developmental stage^24^. Bidirectional transcription at enhancers results in the production of short transcripts, and as expected, RNA polymerase II is enriched at enhancer sequences (Fig. 3b). In contrast to cohesin^COH-1^, which extends several kilobases away from the enhancer itself along the entire length of the fountains, RNA polymerase II is only enriched at the tip of the active enhancers (Fig. 3ab; refs ^6,24^). Interestingly, we found that the enrichment of RNA polymerase II at active enhancers correlated with the abundance of COH-1 (Spearman correlation coefficient R=0.63), which in turn determined fountain sizes (Fig. 2b). These findings prompted us to explore the interplay between enhancer transcription and fountain formation.

**Figure 3.**
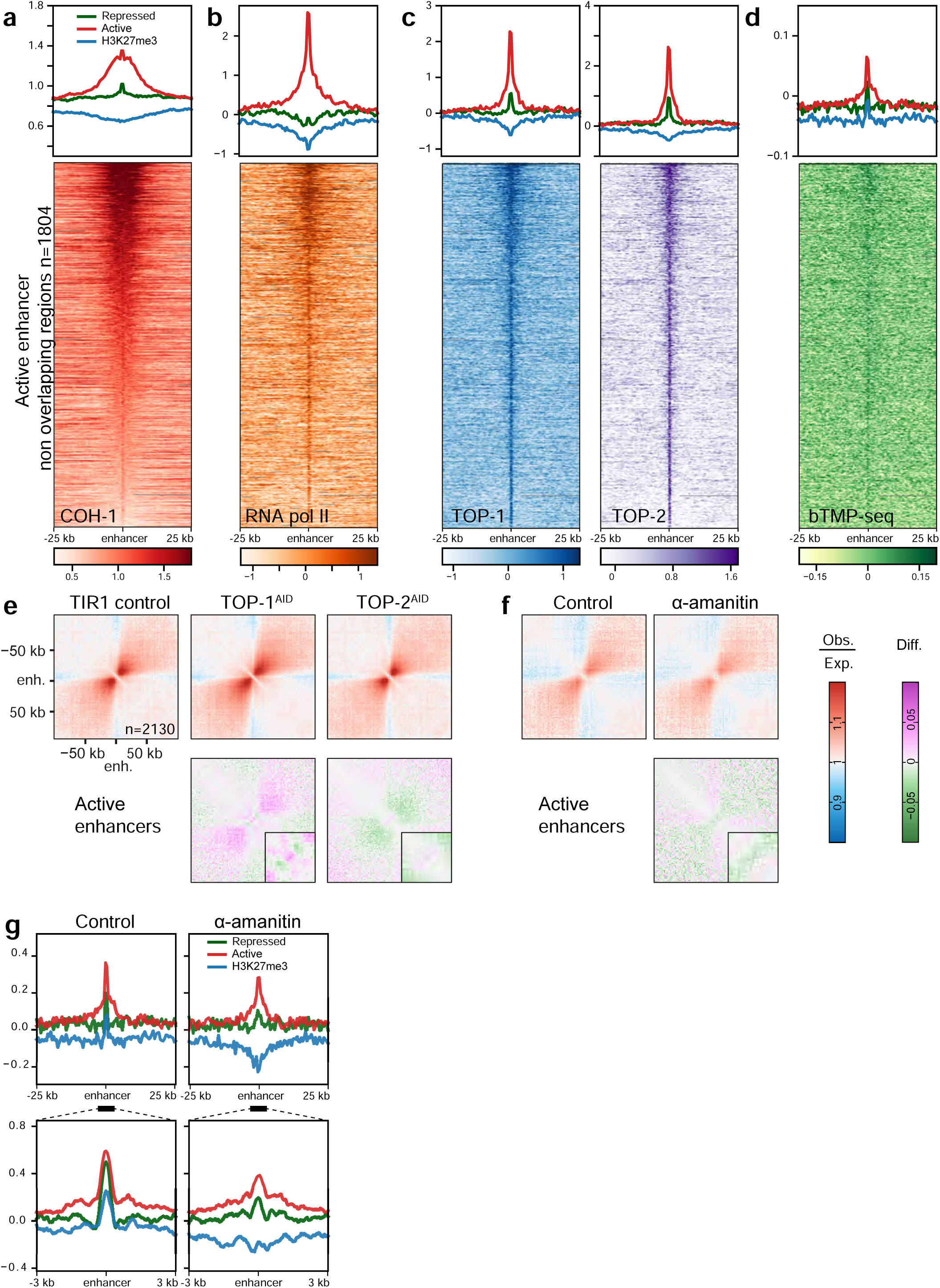
Active enhancers are bound by topoisomerase and enriched for the negative supercoil binder bTMP. (top) Average ChIP-seq profiles at active or repressed enhancers as well as H3K27me3-enriched regions for **a.** COH-1 (young adults), same as in 2b for reference. **b.** RNA pol II (L3), **c.** (left) TOP-1 (L3) and (right) TOP-2 (L3). (bottom) **d.** (top) Average bTMP-seq profiles in L3 at active or repressed enhancers and H3K27me3-enriched regions, (bottom) Heatmap of bTMP binding in active enhancer regions, sorted by COH-1 ChIP-seq enrichment. **e.** Depletion of topoisomerases slightly alters fountain strength, especially at the tip. Average contact frequency maps and difference to control, centered on active enhancers. (left) TIR1 control, (right) TOP-1 or TOP-2 auxin-mediated degradation. (bottom) The difference between the depletion of the topoisomerases and the control, (inset) zoomed in at the fountain tip. **f.** Inhibition of transcription with 𝛂-amanitin slightly alters fountain strength, especially at the tip. Average contact frequency maps and difference to control centered on active enhancers. (left) control, (right) 𝛂-amanitin inhibited transcription. (bottom) The difference between transcription inhibition and the control, (inset) zoomed in at the fountain tip. **g.** Average bTMP-seq profiles upon transcription inhibition in L3 (top) centered at active, or repressed enhancers, as well as H3K27me3-covered regions with 25 kb flanking regions (500 bp bin size), (bottom) 6 kb window centered on enhancers (20 bp bin size).

During transcription, the opening of the double helix and movement of the RNA polymerase generates negative supercoils behind the polymerase, and in the case of divergent transcription at enhancers, negative supercoils accumulate between the two polymerases progressing in opposite directions. In front of the polymerases, positive supercoils are generated, in other words, on either side of the bidirectionally transcribed enhancer sequences. Topoisomerases, the enzymes which relax supercoils by cleaving and unwinding DNA, are therefore expected to be present at active enhancers. Conversely, loop extrusion by cohesin has been shown to modify the topological state of DNA *in vitro*^25^. Indeed, a clear, specific and correlated enrichment for both TOP-1 and TOP-2 was observed at active enhancers (Fig. 3c, R=0.83). Additionally, for both topoisomerases, the enrichment correlated with cohesin^COH-1^ abundance (R=0.6 and 0.59, respectively). As for RNA polymerase II, the TOP-1 and TOP-2 ChIP-seq signal was limited to the enhancer sequences at the tip of the fountains. These findings suggest that topoisomerases are regulating the supercoiling at enhancer sequences created by repeated small-stretch transcription. To directly test whether enhancer sequences are indeed negatively supercoiled, we performed biotinylated psoralen crosslinking followed by sequencing (bTMP-seq; ^26^). bTMP intercalates preferentially with negatively supercoiled DNA and can be crosslinked to DNA using long-wavelength UV irradiation. As for topoisomerases, bTMP was enriched in the tip of the fountains at active enhancer sequences, demonstrating that these sequences are negatively supercoiled *in vivo* (Fig. 3d). Importantly, similar enrichments were observed at the detected fountain tips when plotting the abundance of COH-1, RNA pol II, TOP-1/-2 or bTMP across the 935 fountains (Fig. S6a-d).

To further investigate the role of topoisomerases in the maintenance of fountains, we created average contact frequency maps centered on active enhancers from topoisomerase depletion data (Fig. 3e; ref. ^24^). In these experiments, GFP- and degron-tagged topoisomerases were depleted by a one hour treatment with auxin, leading to the disappearance of the nuclear GFP signal and uniform depletion of the ChIP-seq signal across the genome^24^. As for our data, average contact maps in control animals showed clear fountains (Fig. 3e, TIR1 control). Upon depletion of either TOP-1 or TOP-2, fountains were only mildly affected and changes impacted mostly the tip of the fountains which show a high enrichment for both enzymes (Fig. 3e, TOP-1^AID^, TOP-2^AID^, inset). TOP-1 depletion led to a slight increase in contacts along the trunk of the fountain, while the contacts on either side of the active enhancers were slightly reduced. Conversely, TOP-2 depletion resulted in a slight decrease in contacts along the fountain trunk, with a limited increase in contacts on either side of the active enhancers (Fig. 3e, TOP-1^AID^, TOP-2^AID^, difference maps to TIR1 control). We conclude that short-term depletion of either TOP-1 or TOP-2 has only a marginal effect on fountains, mainly at the tip.

To further characterize the role of transcription in fountain maintenance, we blocked transcription for five hours using 𝛂-amanitin. We investigated first whether transcription inhibition was accompanied by altered supercoiling by performing bTMP-seq and plotted the average enrichment at different enhancer types upon 𝛂-amanitin treatment (Fig. 3g, 25 kb (top) and 6 kb (bottom) windows centered on enhancers). As expected, since transcription inhibition would decrease supercoiling, 𝛂-amanitin treatment lowered bTMP enrichment on all enhancer types, more strikingly at the H3K27me3-covered regions, although the reason for this remains unclear. At active enhancers, bTMP enrichment was decreased at the enhancers but slightly higher on either side of them, suggesting that the absence of transcription would lead to the relocation of negative supercoils away from the enhancer sequences, but not completely abolish these. Additionally, we performed Hi-C upon transcription inhibition and created average contact maps centered on active enhancers. Similarly to topoisomerase depletion, 𝛂-amanitin treatment only slightly decreased fountain strength (Fig. 3f, difference map to control). Collectively, our findings led us to the conclusion that fountains are enriched for RNA polymerase II, topoisomerases and bTMP, indicating probable DNA supercoiling. However, the presence of either RNA polymerases or topoisomerases is not essential for the maintenance of the fountains, indicating that these structures remain stable for at least one hour without topoisomerases and four hours without transcription once they are formed, most likely stabilized by the presence of cohesins.

### Cohesin^COH-1^ cleavage correlates with transcriptional activation of genes close to active enhancers and fountain tips

We previously analyzed the transcriptional consequences of COH-1 cleavage in entire animals^9^. Analyzing this data in isolation with less stringent filtering criteria, 895 genes were significantly up-regulated, and 993 genes were significantly down-regulated (p<0.05). Most changes in transcript abundance were small, with only 98 up- and 52 down-regulated genes with a fold change larger than 1.41 (|log2FC|>0.5). As ATAC-seq, modified histone ChIP-seq and RNA-seq are made on entire animals with a large number of different cell types, the target gene of each enhancer has been determined only for a handful of enhancers^6,7^. We therefore distributed genes into six categories by assigning each enhancer to the closest TSS (or to none if the gene was not the closest TSS to any enhancer, Fig. 4a). “Active”, “Repressed”, “H3K27me3-covered” categories were TSS of genes closest to only one type of enhancer; “Mixed active and repressed” category were TSS of genes closest to a mixture of enhancers from the active and either of the two categories - repressed or H3K27me3-covered; and “mixed repressed” were genes closest to a mixture of repressed enhancers and H3K27me3-covered regions. We then analyzed the impact of COH-1 cleavage on these different gene categories (Fig. 4a). When no enhancer was present in the vicinity of the gene (the largest number of genes), average expression after cleavage of COH-1 was slightly lower than in the control experiment. For genes proximal to mixed or active enhancers, the change in expression levels of these genes was biased towards higher expression (Fig. 4a, red and violet boxes). In contrast, genes proximal to H3K27me3-covered regions were evenly distributed between up- and down-regulated (Fig. 4a, green box). Therefore, genes close to active enhancers were upregulated upon COH-1 cleavage, suggesting fountain formation limits active enhancer activity. The fact that genes close to a mixture of active and repressed enhancers and even a mixture of the two repressed categories were also biased towards upregulation, suggests a role for cohesin^COH-1^ in regulating complex transcriptional landscapes, though the number of genes in these groups is small.

**Figure 4.**
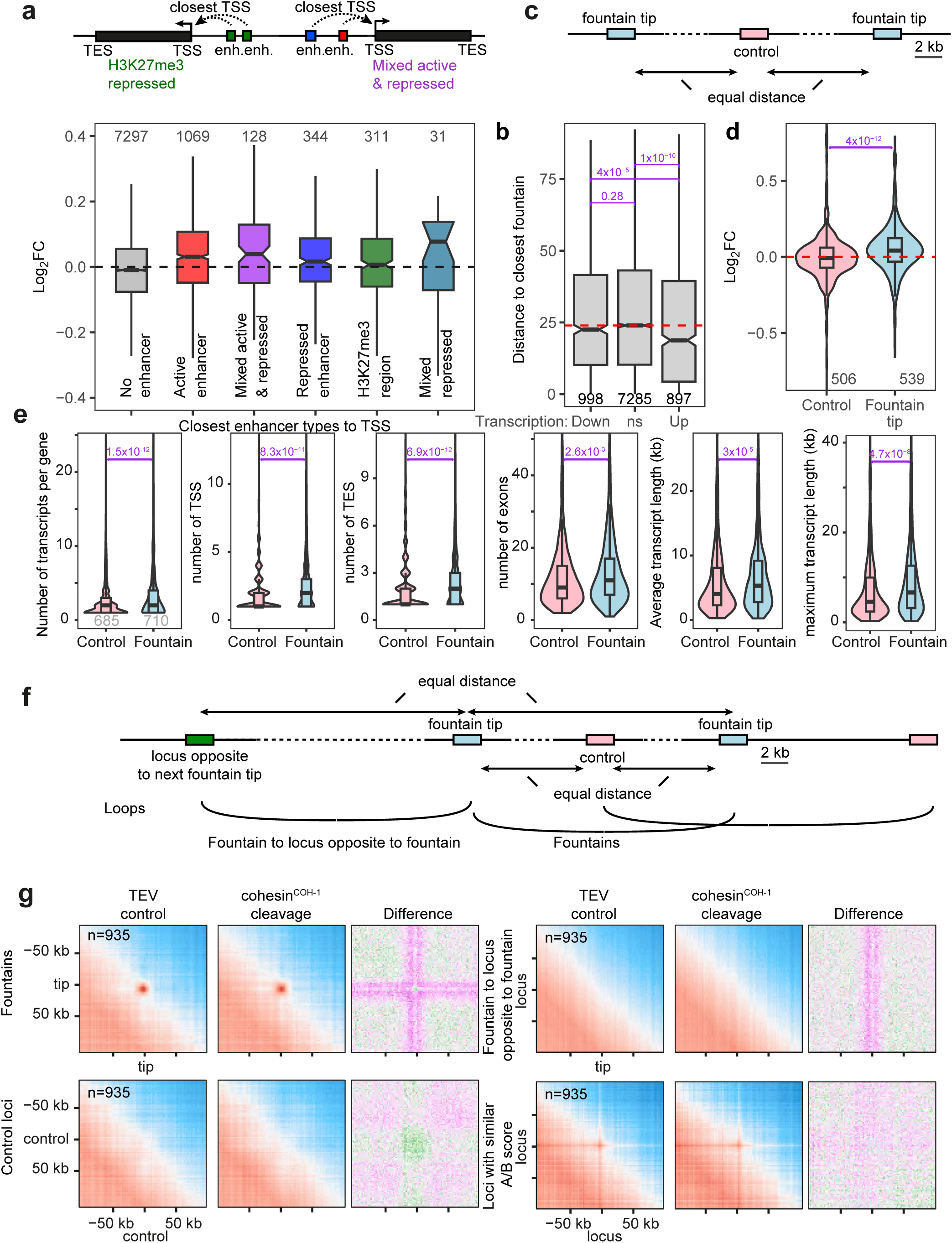
Active-enhancer proximal genes are upregulated upon COH-1 cleavage, fountains cluster in the nuclear space. **a.** Genes were grouped according to the type of L3 stage enhancers from Daugherty *et al.* (2017) that their TSS was closest to. The log2 fold change of all expressed genes upon cleavage of COH-1 are plotted according to enhancer type. **b.** Boxplot of the distance of genes from the nearest fountain tip bin. Genes were grouped by their change in expression upon COH-1 cleavage: up and down regulated or not changing significantly (NS). The dashed red line indicates the median distance to the nearest fountain tip bin of NS genes. In b, d & e, the number of genes in each group is indicated at the bottom of the plot, and adjusted p-values from Wilcoxon rank sum tests are shown in purple. **c.** Schematic representation of regions used in panels d-e. 2 kb fountain tip bins were compared with control bins of the same size located at the midpoint between neighboring fountains. **d.** Log2 fold change of all expressed genes upon COH-1^cs^ cleavage that overlap control bins (506 genes) and fountain-tip bins (539 genes). **e.** Comparison of features of genes that overlap with control and fountain-tip bins. **f.** Graphical depiction of the loops analyzed in g. **g.** Top row, left: average contact maps show loops between pairs of fountain tips under TEV control conditions. No such loops were observed for control loci (second row, left), or between fountain and loci located at the similar distance but on the opposite side to the next fountain (first row, right), or between loci with similar A/B score than fountains (second row, right). Average contact maps are shown both for the TEV control and upon cohesin^COH-1^ cleavage, conditions, as well as the differential map (cleavage/control) - same color scales as in Fig. 2d.

We next analyzed transcriptional changes upon COH-1 cleavage based on the location of the detected fountains (Fig. 2f). For this, we first measured the distance between genes and the closest fountain and split genes based on their transcriptional modulation upon COH-1 cleavage. We found that the average distance to fountain tips of non-changing or down-regulated genes was not significantly different, while upregulated genes were significantly closer to fountain tips (Fig. 4b). Second, we focused on fountain tips and compared those to a set of control regions located at the midpoint between neighboring fountain tips (Fig. 4c). Genes that overlap fountain tips were significantly biased towards upregulation whereas genes overlapping control bins were equally distributed between up and down regulation (Fig. 4d). As fountains are enriched for cohesin^COH-1^, gene upregulation upon COH-1 cleavage might be a consequence of cohesin^COH-1^ enrichment, independently of the fountains themselves. To disentangle cohesin^COH-1^ enrichment from fountain proximity, we compared gene expression changes in fountain tip bins (+/- 2 kb) *versus* genomic bins with similar cohesin^COH-1^ enrichment than fountain tips yet located more than 30 kb away from these (Fig. S7a). Upon cohesin^COH-1^ cleavage, fountain tip genes were significantly upregulated whereas non-fountain bins similarly enriched for cohesin^COH-1^ were not (Fig. S7b-d). Repeating a hundred times the same experiment by picking different fountain tips/non fountain tip bins led to the same result (Fig. S7e). Together, this demonstrates that fountain ablation by COH-1 cleavage correlates with upregulation of fountain tip-proximal genes.

Interestingly, fountain tips and their neighboring bins contain genes more highly expressed than the average of all genes in the 40 kb surrounding the fountain tip (Fig. S8a-c). Asymmetric fountains show a slightly higher gene expression in the tip-adjacent bin on the side where the fountains are shortest (Fig. S5a) and fountain disappearance upon COH-1 cleavage did not affect this asymmetry in relative transcription levels (Fig. S5b), suggesting that highly expressed genes could block cohesin^COH-1^ loop extrusion and be the cause, not the consequence of fountain asymmetry. Changes in gene expression upon cohesin^COH-1^ cleavage had a limited correlation with the number of enhancers at the fountain tip: considering 6 kb regions overlapping fountain tips and their two neighboring bins, there is an increase in log2 fold change upon COH-1 cleavage as the number of enhancers in those bins increases from 0 to 1 to 2, however after that, the linear relationship breaks down (Fig. S8de). Additionally, genes overlapping fountain bins were more complex than genes overlapping control bins: they had a significantly greater number of transcripts, TSSs and Transcription End Sites (TES) per gene, a larger number of exons, and both their average and maximum transcript length was greater (Fig. 4f), indicating that fountains could be necessary for the regulation of complex transcriptional landscapes.

### Fountain tips cluster in 3D hubs inside the nuclear space

Active enhancers exhibit a well-documented tendency to spatially cluster, forming regulatory hubs enriched in transcription factors, Mediator, and coactivators that cooperatively regulate gene expression^27,28^. To assess whether fountain tips harboring active enhancers similarly engage in higher-order clustering in *C. elegans,* we quantified average contact probabilities between *cis*-located fountain tips and corresponding control regions as outlined above (Fig. 4f). Our analysis revealed a marked and specific enrichment of contacts between fountain tip pairs, in contrast to control regions or to equidistant loci situated on the opposite flank of adjacent fountains (Fig. 4g). Notably, loci with matched A/B compartment scores and equidistant spacing relative to fountain tips exhibited only modest levels of interaction, consistent with weak chromatin state–driven clustering (Fig. 4g). These findings suggest that fountain tips cluster within the nuclear space, forming subnuclear fountain domains. To interrogate the role of cohesin in modulating fountain contacts, we examined changes in average contact frequencies following cleavage (Fig. 4g, cohesin^COH-1^ cleavage). Contact frequencies between fountain tips slightly decreased, whereas contacts between fountain trunks and neighbouring loci increased, leading to a cross-like pattern on the differential contact frequencies map. This redistribution of contact patterns extended into regions flanking the fountain tip by approximately ±14 kb, indicating a localized architectural perturbation. The observed pattern strongly supports a model in which cohesin-mediated loop extrusion forms spatially insulated fountain domains, with tips acting as hubs for enhancer clustering. Upon cohesin cleavage, these looped architectures disassemble, leading to slight contact depletion at tips and ectopic interaction enrichment along the linear genome, consistent with loss of insulation and increased trunk promiscuity.

We further investigated the impact of cohesin^COH-1^ cleavage on enhancer-promoter contacts. Promoters were first ranked either by the number of contacts they made with active enhancers under control conditions or by their linear distance to the nearest active enhancer. We then calculated the ratio of contact frequencies between cohesin^COH-1^ cleavage and control conditions, alongside changes in transcriptional activity of the corresponding gene. Consistently and independently of the chosen set of active enhancers^6,7^, the contact ratio (cleavage/control) decreased for promoters that had high contact frequencies in the control situation, while it increased for promoters with low contact numbers in the control situation (Fig. S9ab). When promoters were instead ranked by their distance to enhancers, this effect was less pronounced: contact counts ratios remained close to 1 at very short distances and were only modestly reduced at longer distances (Fig. S9cd). Transcriptional changes following cohesin^COH-1^ cleavage also depended on the baseline probability of enhancer-promoter contacts under control conditions. Promoters with high contact frequencies showed increased transcription upon cohesin cleavage, whereas those with fewer contacts exhibited little of no significant change (Fig. S9ef). Together, these results suggest that cohesin^COH-1^ loading at active enhancers serves a repressive function, limiting transcription at their target promoters by modulating enhancer-promoter contact frequencies.

### Genes upregulated upon COH-1 cleavage are mostly neuronal

Enhancers are believed to drive cell-type specific expression, and indeed putative nematode enhancers whose activity was individually tested in animals show cell-type specific expression^6,7^. We reasoned that if COH-1 cleavage led to mild upregulation of genes proximal to active enhancers, this might be due to cell-type specific expression of the latter genes, as RNA-seq is performed on entire animals. Small overall changes would be observed even if transcription levels in individual cells experience major modulation if these genes are expressed only in a few cells per animal. To interrogate whether genes with changed expression levels would belong to a specific cell type, we used two different approaches to analyze the list of significantly up and down regulated genes^29^. First, we performed Tissue Enrichment Analysis (TEA) and GO term enrichment, which allows us to determine in which cell type a gene set is most likely expressed. Genes upregulated upon COH-1 cleavage showed a striking enrichment for neuronal cell types, as well as GO terms and phenotypes associated with neurons (Fig. S10a-b). In contrast, downregulated genes were associated with a range of non-neuronal tissues, in particular germline cells and core cellular and metabolic processes (Fig. S10a-b). Second, we asked in which cells genes significantly up- or down-regulated were expressed using previously published single-cell RNA-seq datasets^30^. To this aim, we plotted the percentage of genes in the lists of up- or down-regulated genes expressed in each cell type and overlaid this percentage as a color code to the scRNA-seq UMAP. As for TEA, we found that genes upregulated upon COH-1 cleavage were mostly expressed in neuronal cell types, while genes downregulated were mostly expressed in the germline (Fig. S10c-f). Altogether, both analyses strongly suggest that genes upregulated upon COH-1 cleavage are expressed in neurons.

### Cohesin^COH-1^ cleavage leads to isoform switch of the nematode *Nrf* homolog *skn-1*

We next sought strains in which transcription factors were endogenously tagged and located either close or inside a fountain region, in particular transcription factors expressed in neuronal cell types. *skn-1/Nrf* is such a gene, as its TSS lies between two very close fountains. The tip of one fountain lies within the gene body of *skn-1* whereas the other fountain lies about 5 kb upstream of the *skn-1* gene, in the *bec-1* enhancer region, and partially merges with the first one and could not be detected by our algorithm as a separate fountain due to their proximity (Fig. 2ef, 5a). In RNA-seq data, *skn-1* shows a modest but significant 1.15 fold upregulation upon COH-1 cleavage compared to TEV control. *skn-1* expression has been widely studied, due to its function in lifespan regulation (^for^ ^review^ ^31^). SKN-1 has three isoforms named A to C, arising as a result of the use of different TSS and alternative splicing (Fig. 5a). At the third larval stage, the *skn-1c* isoform was undetectable by RNA-seq, hence we focused on *skn-1a* and *skn-1b*. The SKN-1A transcription factor is present in intestinal cells and normally located in the cytoplasm due to its membrane targeting signal at the N-terminus of the protein. Upon stress, SKN-1A relocates to the nucleus for transcriptional gene regulation^32,33^. In contrast, SKN-1B is exclusively nuclear and solely expressed in two head neurons called ASI (Fig. 5b, TEV control; ^34^).

**Figure 5.**
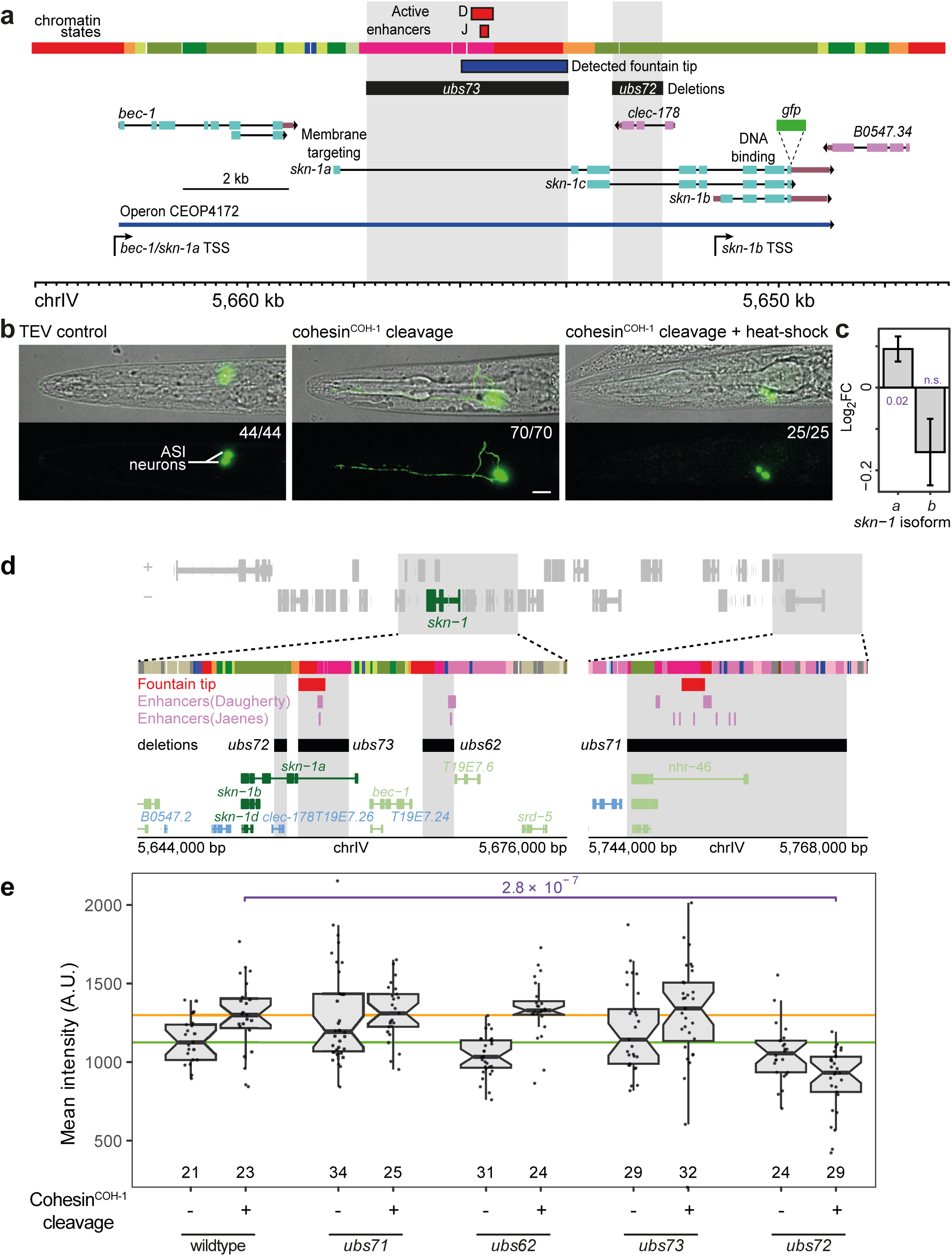
*skn-1*/*Nrf* switches isoform upon cohesin^COH-1^ cleavage. **a.** Structure of the *skn-1* genomic region, highlighting the three canonical *skn-1* isoforms *a* to *c*, the location of the GFP transgene insertion used in b, the characterized active enhancer (organism-wide), the detected fountain location (organism-wide) and the location of two of the deletions used in e. Active enhancers from Daugherty et al.^7^ (D) or Jaenes et al.^6^ (J). **b.** Expression of *skn-1* in the head of control animals, upon COH-1 cleavage or upon COH-1 cleavage and heat stress 30’ prior imaging. Scale bar: 10 µm. Numbers on the upper left corner of the lower panel indicate the number of animals in which the depicted phenotype has been observed (first number) and the number of animals imaged (second number). **c.** Transcript level differential expression of *skn-1* isoforms upon COH-1 cleavage. Adjusted p values from Wald test are shown next to the bars (n.s.: not significant). **d.** Structure of the larger *skn-1* locus, depicting chromatin states (colors as in Fig. 2g), detected fountain tips, active enhancers, as well as the putative active enhancer deletions. **e.** Boxplots of mean fluorescence intensity in ASI nuclei/cell-bodies in control conditions (no cohesin^COH-1^ cleavage, -) and upon cohesin^COH-1^ cleavage (+) for the different deletion strains depicted in d. The horizontal green and orange lines show the median of the average fluorescence intensity of the ASI nuclei for the wildtype locus without or with cohesin^COH-1^ cleavage, respectively. The number of nuclei/cell-bodies in each group is shown at the bottom of the boxplots. FDR adjusted p-values from a two-sided Wilcoxon rank sum test comparing control (no TEV expression) and cohesin^COH-1^ cleavage conditions for each enhancer deletion strain are shown in purple in between the two panels. Only significant p values are shown.

At the genomic level, *skn-1a* and *skn-1b* share several exons on the 3’ part of the transcripts, but transcription of the two isoforms is controlled by different promoters. The *skn-1b* isoform is transcribed from its own promoter while the *skn-1a* isoform is part of an operon whose promoter is located upstream of the *bec-1* gene (Fig. 5a). Several active enhancers around the *skn-1* gene have been independently identified in entire animals^6,7^ (Fig. 5ad, detailed in S11). One of these enhancers is located between the first and the second exon of *skn-1a*, although it remains unclear in which cells this enhancer is active (Fig. 5a, red squares labelled D^7^ and J^6^). Using a C-terminally GFP-tagged construct which labels all SKN-1 isoforms, we observed that in control animals at the L3 stage, most fluorescence was indeed visible in the nuclei of the two ASI neurons, since the low, diffuse cytoplasmic expression in the intestine is not visible above the autofluorescence of the gut granules, as expected from previous studies (n=44, Fig. 5b, TEV control; ref. ^35^). This exclusively nuclear fluorescence in ASI identifies SKN-1B as the expressed isoform. Upon COH-1 cleavage, a clear increase of the GFP signal was scored, in line with *skn-1* upregulation observed by RNA-seq (Fig. 5c, fluorescence quantified in 5e, wildtype). Additionally, we detected in all imaged animals (n=70) some degree of accumulation of the GFP-tagged protein in the neuronal cytoplasm, filling the entire cells in most animals, including the sensory dendrites located in front of the cell body as well as the laterally extending axons (Fig. 5b, cohesin^COH-1^ cleavage, fluorescence pattern quantified in S12b). This cytoplasmic localization required COH-1 cleavage as heat-shocked animals with TEV cleavage sites in COH-1 but no TEV expression transgene only showed nuclear fluorescence (n=21, data not shown). The presence of cytoplasmic SKN-1 in ASI neurons has never been observed previously (pers. comm. K. Blackwell, HMS). To further assess the nature of the *skn-1* isoform, we took advantage of the stress-mediated nuclear relocation of SKN-1A. We heat-shocked the animals a first time at the L1 stage to induce COH-1 cleavage, and a second time at the L3 stage, 30 minutes before imaging. Under those conditions, all imaged animals (n=25) showed no cytoplasmic ASI fluorescence anymore, as expected due to the relocation of SKN-1A to the nucleus upon stress activation (Fig. 5b, cohesin^COH-1^ cleavage + heat-shock). Differential expression analysis of *skn-1* isoforms at the transcript level in entire animals showed that *skn-1a* was indeed upregulated upon COH-1 cleavage (Fig. 5c, p=0.02), though the downregulation of the more lowly expressed *skn-1b* isoform was not significant. Additionally, after the cleavage of COH-1, we observe an increase in the expression of the *bec-1* gene to the same levels as *skn-1a,* as expected since both genes are part of the same operon (data not shown). We conclude that cleavage of cohesin^COH-1^ has dual effects: it upregulates *skn-1* expression and induces significant changes in the isoforms expressed. Specifically, there is a transition in promoter usage from *skn-1b* to the operon promoter of *bec-1* and *skn-1a*, resulting in the production of the non-neuronal protein SKN-1A. These findings suggest that the cohesin^COH-1^-mediated formation of fountains direct promoter use, most likely by restricting active enhancers activity.

### *skn-1b* proximal sequences regulate *skn-1a* upon COH-1 cleavage

This model predicts that deletion of one or more active enhancer(s) should reduce transcriptional activity and/or hinder the switch from *skn-1b* to *skn-1a* upon COH-1 cleavage. We tested this by deleting putative regulatory active regions located close to or inside the *skn-1* gene, guided by gene activity or organism-wide mapping data^6–8,23,36^ (Fig. 5d, detailed in S11). One limitation here is that enhancers and fountains were identified in entire animals, hence their specific activity in ASI neurons could not be assessed using Hi-C or ATAC-seq data.

We first deleted two distal regions containing putative active enhancers : the entire *nhr-46* gene, located 100 kb upstream of the promoter of *bec-1/skn-1a* operon (Fig. 5d,S11ab; *ubs71*), and a large enhancer-rich region situated 5 kb upstream of the *bec-1* gene (Fig. 5d,S11ab; *ubs62*). In the *ubs71* deletion strain, *skn-1b* expression was modestly elevated in control animals, but GFP intensity remained comparable to the wild-type locus while the COH-1 cleavage-induced isoform switch was slightly reinforced (n>20 animals for each deletion and condition, here and below, quantification in Fig. 5e, S12b). This indicates that this active enhancer is not required for *skn-1a* activation. The ASI-specific SKN-1::GFP mean intensity in the *ubs62* putative enhancer deletion strain was slightly lower in control conditions (∼92% of the wildtype locus strain), but upon COH-1 cleavage, expression was restored to wild type levels (Fig. 5e, *ubs62*). Additionally, strong cytoplasmic GFP localization was fully penetrant in this strain upon COH-1 cleavage (Fig. S12b). These results suggest that neither region is essential for *skn-1b* expression or *skn-1* activation. However, the enhanced signal observed in both deletion strains (Fig. S12b) raises the possibility that these enhancer-containing regions may contribute to the insulation of the *bec-1/skn-1a* promoter from neighbouring active enhancers.

Subsequently, we deleted the first intron of *skn-1a* located between the first exon of *skn-1a* and the *skn-1b* TSS, reducing its size from 4347 bp to 551 bp (Fig. 5d,S11ab; *ubs73*). The switch from *skn-1b* to *skn-1a* upon COH-1 cleavage remained unchanged (Fig. S12b). However, expression of *skn-1b* as well as increased expression of *skn-1* upon COH-1 cleavage was more variable than in control strains (Fig. 5e). The first intron of *skn-1a* thereby appears to stabilize *skn-1a* and *skn-1b* expression, although it is not essential for their transcription. We next deleted a 959 bp region in the third intron of *skn-1a*, 955 bp 5’ of *skn-1b* TSS and 10 kb downstream of *bec-1/skn-1a* operon promoter, overlapping the *clec-178* gene (Fig. 5de,S11ab; *ubs72*). *clec-178* is exclusively expressed in coelomocytes, phagocytic cells located in the body cavity, yet DNase-I accessible region conformation capture (ARC-C)^8^ shows that the *clec-178* region harbors significant contacts with the *skn-1b* promoter and the first intron of *skn-1a*, suggesting the presence of an active enhancer in this region (Fig. S11ab, ARC-C). Upon deletion of the *clec-178* region, *skn-1b* expression was marginally reduced in control animals (94% of wildtype locus, Fig. 5e, *ubs72*). However, and in contrast to the wildtype locus or all other deletions, COH-1 cleavage led to a significant downregulation of *skn-1* and reduced switching to *skn-1a* isoform (Fig. 5e,S12b). Sequences in the deleted region therefore contain sequences marginally necessary for *skn-1b* expression in ASI in control animals, but these sequences are required for *skn-1* upregulation and the switch to the *bec-1/ skn-1a* operon transcription upon COH-1 cleavage. Our interpretation is that in ASI neurons, sequences in the *clec-178* gene behave as loading sites for cohesin, creating a fountain which insulates the inactive *bec-1/skn-1a* operon promoter from the transcriptionally active *skn-1b*. Upon COH-1 cleavage and fountain ablation, these regions as well as other active enhancer regions nearby contact the operon promoter, leading to its activation. Deletion of *clec-178* sequences removes the closest highly active enhancer to *skn-1a* in ASI, leading to decreased use of the *bec-1/skn-1a* TSS upon COH-1 cleavage.

### COH-1 cleavage minimally delays animal growth and development

To further assess the function of cohesin^COH-1^, we measured the impact of COH-1 cleavage on animal growth during larval development with a time resolution of 10 minutes^37^ (Fig. S12a). COH-1 cleavage led to a very small difference in animal volume compared to TEV control (Fig. S12bc). However, this difference was minor compared to previously characterized mutants (*eat-2*, *dbl1*, *raga-1;* ref. ^37^). We conclude that COH-1 cleavage and fountains disappearance has a minimal effect on animal growth.

### COH-1 cleavage has a broad impact on animal behavior

Since neuronal gene expression is disturbed by COH-1 cleavage (see above), we wanted to analyze its impact on nervous system function using computer-assisted high-content behavioral quantification of worm crawling behavior^38^. We compared 47 core postural and locomotion parameters in wild type (N2), TEV control and upon cohesin^COH-1^ cleavage. Animals either dwelled on food or were in search of food after 6 h of deprivation (Fig. 6). We found that COH-1 cleavage affected many behavioral parameters in both conditions, whereas the behavior of TEV control animals was indistinguishable from that of wild type (Fig. S13). Some behavioral differences were quite obvious (Fig. 6a and Supplementary Movie 1 and 2). These broad behavioral differences caused animals with cleaved COH-1 to cluster separately in a principal component analysis (PCA) (PC1/PC2 space, Fig. 5b). Among the most salient features, COH-1 cleavage caused a striking increase in the body curvature of both fed and starved animals (Fig 6ae) and impaired the ability of worms to implement food search behavior upon food deprivation (Fig. 6c-g). In wild type animals, the entry into the food search locomotion state involves multiple coordinated behavioral changes^39,40^. These include a marked elevation of animal speed and of the frequency of reorientation events called omega turns (Fig 6cd). As a consequence, worms produce relatively twisted trajectories (Fig. 6f) but disperse fast, dramatically increasing worm displacement as compared to fed animals (Fig. 6g). The up-regulation of both speed and omega turn frequency upon food-deprivation was strongly reduced upon COH-1 cleavage (Fig. 6cd), which impaired dispersal (Fig. 6fg). In contrast, TEV control animals behaved essentially like wild type ones (Fig 6a-g). Collectively, these results demonstrate that the post-developmental cleavage of COH-1 produces a broad impact on animal locomotion and are in line with a model in which COH-1-dependent fountain formation is essential for normal neuronal gene expression and the ensuing proper function of the nervous system.

**Figure 6.**
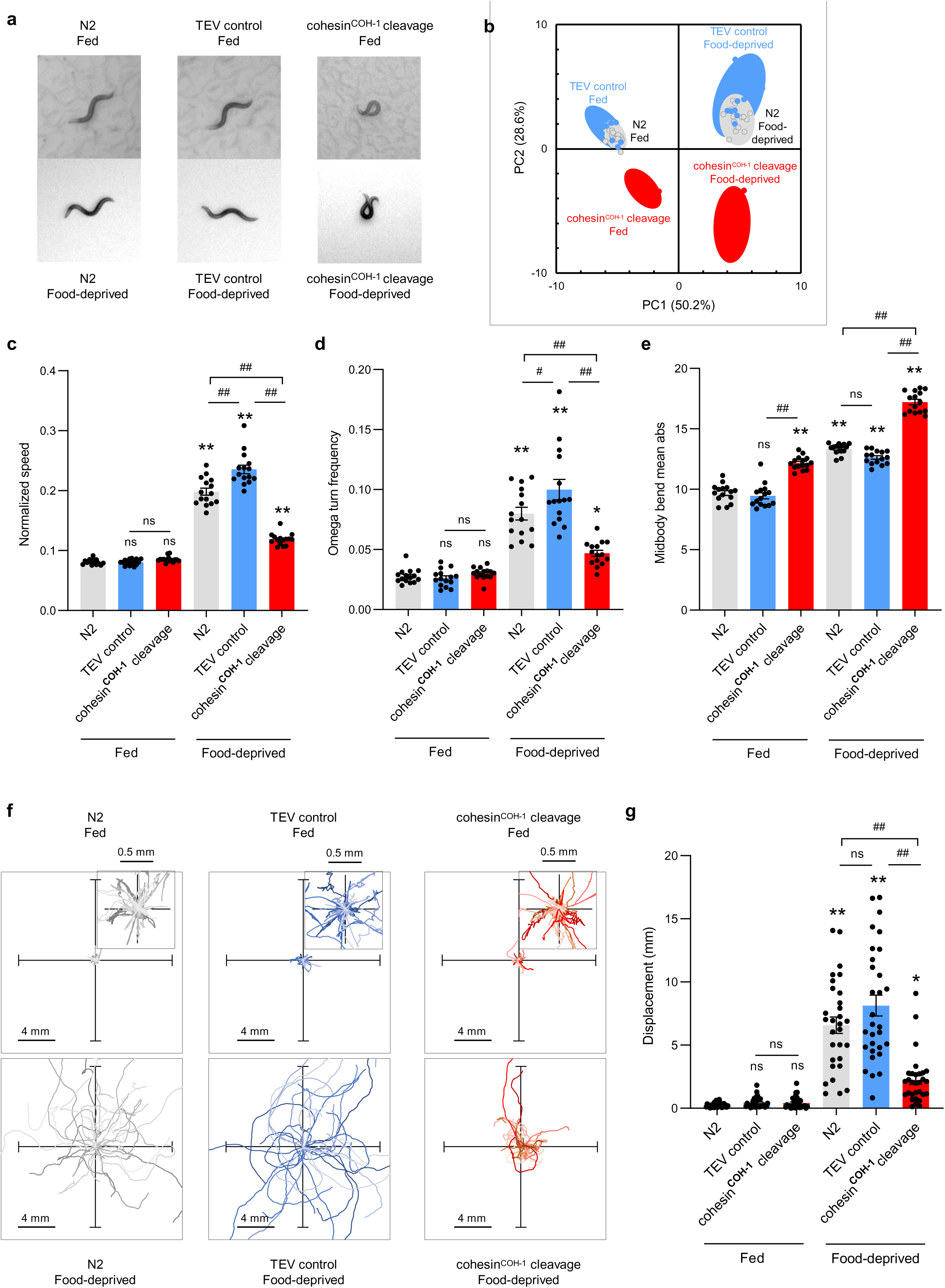
COH-1 cleavage affects the nervous system function. **a.** Representative pictures of young adult *C. elegans* either dwelling on food (fed) or foraging off-food 6 hours after food deprivation (food-deprived), illustrating postural differences between the indicated genotypes. **b**. Multidimensional behavioral states of animals presented as projections over the two main principal components (PC1 and PC2) from a single principal component analysis (PCA) over 47 postural and motion parameters, and over all the conditions. The proportion of variance explained by each PC is indicated in the axis labels. Upper left and upper right quadrants correspond to the normal dwelling and food search behavioral states adopted by wild type or TEV control worms. The lower left and right quadrants correspond to markedly altered behavioral states caused by COH-1 cleavage. Averages positions of individual replicates as data marks, 95% CI as colored ellipses. Each replicate is a separate population with ≥40 animals. **c-e**. Selected behavioral parameters reported as average ± s.e.m. of n*=*15 independent replicates. Each replicate value (data points) corresponds to the average value of a separate population with ≥40 animals. Additional behavioral alterations are presented in Fig. S14. **f**. One-minute worm trajectories (35 for each condition) plotted from a single starting (0,0) coordinate. Enlarged representations for fed worms in dwelling state (insets). **g**. Dispersal quantification. Average ± s.e.m. and individual data points for animal displacement (corresponding to how far animals moved from their starting point). *n*=30 animals. **c**, **d**, **e**, **g**. *, *p*<.05 and **, *p*<.01 versus N2 fed condition, # *p*<.05 and ##, *p*<.01 versus the indicated control by Bonferroni posthoc tests.

## Discussion

### At least three SMC complexes create looping domains on nematode chromosomes

In mammals, cohesin is the main loop extruder acting on interphasic chromosomes and its removal leads to the disappearance of most 3D structures including loops and TADs^41–43^. This study, combined with our previous data, demonstrates that at least three different SMC complexes are at play on interphase nematode chromosomes^9^. On the one hand, large-scale chromosome looping is achieved by condensin I creating loops larger than 100 kilobases, while a variant of condensin I specifically targeted to the X chromosome creates specific loop domains resembling TADs on this chromosome^44^. On the other hand, cohesin^COH-1^ achieves short-patch loop extrusion in the tens of kilobases range centered on active enhancers, giving rise to fountains. This is markedly different from Vertebrates, in which different SMC complexes extrude loops during different cell cycle stages, with cohesins and condensins active during interphase and mitosis, respectively^45^. In nematodes, the presence of different interphasic structures maintained by cohesin^COH-1^ or condensin I variants is unique, suggesting the simultaneous activity of these complexes, which has not been previously observed in Metazoan species.

Peculiar to nematodes of the *Caenorhabditis* genus, two divergent copies of the cohesin kleisins are present in somatic cells (as well as three additional ones in the germline). Early reports suggest that the two somatic kleisins play different roles, as cohesin^COH-1^ is present in somatic cells independently of the cell cycle phase while cohesin^SCC-1^ is exclusively expressed in dividing cells^20,46^. Together with previous work in which we tested the mitotic function of the different SMC complexes, our data demonstrate a functional specialization of the two somatic cohesin complexes: cohesin^SCC-1^ holds sister chromatids together during mitosis, in line with the observed *scc-1* mutant phenotypes due to chromosome segregation defects in dividing cells ^20^. In contrast, cohesin^COH-1^ has a minor role in sister chromatid cohesion, although COH-1 is the most abundant somatic kleisin^9^. The function of cohesin^COH-1^ is therefore small-scale chromatin loop extrusion during interphase, leading to the formation of enhancer fountains. This separation of function between the two cohesin complexes allows to untangle the mitotic and interphasic functions of cohesin.

### Nematode enhancers correlate with fountain 3D structures

Enhancers play a critical role in transcriptional gene regulation by conferring cell type and developmental stage-specific gene expression. However, their activity must be tightly regulated to prevent unintended activation of non-target promoters^1^. In vertebrates, this regulation is largely mediated by the formation of TADs, established by the combined action of cohesin-mediated loop extrusion and CTCF-bound boundary elements. These 3D chromatin domains restrict enhancer-promoter contacts across domain boundaries^3^, thereby minimizing ectopic gene activation. In nematodes, however, high-resolution chromatin conformation capture studies failed to detect TAD structures on autosomes^24,44,47–49^, raising the question of how enhancer-promoter specificity is maintained in the absence of TADs. Here we present evidence for the presence of loose, tens-of-kilobase-sized 3D structures centered on active enhancers, which resemble previously described flares, hinges, plumes, or jets in other organisms^10–13^. During the preparation of this manuscript, an independent study identified the same cohesin-dependent structures at active enhancers in *C. elegans,* using an orthogonal degron-based acute degradation system targeting the cohesin subunit SMC-3 and the cohesin unloader WAPL-1^50^.

These structures, which we term fountains, appear to be evolutionarily conserved. Their focal points are marked by H3K27ac - a histone modification associated with enhancer activity - not only in nematodes but also in zebrafish sperm and during zygotic genome activation (ZGA)^15^, as well as in WAPL/CTCF-depleted mouse embryonic stem cells (mESCs) and primary thymocytes^10^. As in nematodes, ZGA fountains in fishes colocalize with active enhancers^11,15^.

In thymocytes^10^, the projection angle of jets is modulated by the presence of proximity of CTCF boundaries, resulting in asymmetric loop extrusion. Similarly, nematode fountains can be asymmetrical (Fig. 2h). However, as *C. elegans* lacks a CTCF homolog, this suggests that alternative genomic or chromatin-based barriers may constrain directionality. Notably, genes located on the more-extruded side of asymmetric fountains exhibit lower expression compared to those on the opposite side (Fig. S2), indicating a possible impact transcription may play a role as genes located on the most-extruded side of the asymmetric fountains show a lower expression than genes located on the other side (Fig. S2). This difference between enhancer sides is more likely the cause of fountain asymmetry than a consequence of asymmetric fountain formation as removal of fountains by COH-1 cleavage did not change transcription level asymmetry. Conversely, transcription inhibition did not significantly modify asymmetric fountains. Instead, chromatin states associated with enhancers are more enriched on the more-extruded side, suggesting that these states might be more permissive to loop extrusion. A similar interplay between transcription and loop extrusion has been observed in Bacteria and WAPL/CTCF-depleted mammalian cells, where loop extrusion is impeded by polymerases, while SMC complexes might be pushed by transcribing complexes^19,51–53^.

Our analysis of published RNA polymerase II and topoisomerases ChIP-seq data showed that active enhancers are enriched for these factors (Fig. 3bc; ref. ^24^). RNA polymerase presence at fountain tips aligns with known bidirectional transcription at active enhancers, while topoisomerase enrichment suggests the accumulation of torsional stress and supercoiled DNA^6^. Indeed, bTMP mapping revealed specific enrichment of negatively supercoiled DNA at active enhancers as well as fountain tips. This implies that, in addition to forming chromatin loops, cohesin may generate supercoils of plectonemes during fountain formation, as observed *in vitro*^25^. Fountains may also act to locally restrict the dissipation of torsional stress generated by enhancer transcription or by the loop extrusion activity of cohesin loaded onto enhancers^22^. Degron-mediated degradation of topoisomerases I or II only slightly modified fountain strength (Fig. 3e). Topoisomerase I depletion resulted in increased contact frequencies at the fountain tip (the active enhancer locus) and generally stronger fountains, suggesting that topoisomerase I reduces local compaction. Conversely, topoisomerase II depletion led to reduced contact frequencies along the fountain trunk, consistent with a more relaxed or extended fountain structure. Transcription inhibition using 𝛂-amanitin generally slightly weakened fountains, but did not ablate them. Fountain maintenance therefore appears to be largely independent of topoisomerase activity or transcription, suggesting that once set up, fountains no longer need transcription to remain stable.

### Fountains repress active enhancer-proximal gene activity

Our results provide compelling evidence for a direct link between fountain structures and enhancer-mediated transcriptional regulation. Cleavage of cohesin^COH-1^, leading to the disappearance of fountains, is accompanied by transcriptional upregulation of genes located near fountain tips and active enhancers as well as by changes in transcript isoform usage (Fig. 4,5). This transcriptional change was not observed upon in a previous study employing acute SMC-3 degradation^14^, likely due to the shorter timeframe between cohesin depletion and sample collection in that study (1-hour degradation versus 19-hours induction of cohesin^COH-1^ cleavage in our system). Given the inherent stochastic nature of enhancer-promoter contacts and transcriptional activation (for review, see ^54^), it is plausible that changes resulting from fountain loss require more time to manifest at the transcriptional level. Our findings strongly support the notion that fountains act as repressors of active enhancer function. This upregulation of genes at fountain tips may result from multiple, spatially distinct mechanisms. Locally, the loss of cohesin may alleviate physical interference with the transcriptional machinery, reducing collisions between elongating RNA polymerases and cohesin complexes^19,55^. This model is supported by previous work showing transcriptional repression caused by repeated cohesin passage in mammalian cells following artificial targeted cohesin loading^4,56^, and may be especially relevant at fountain tips, where RNA polymerase II density is high (Fig. 3). Such a mechanism could represent a form of enhancer self-regulation: by recruiting cohesin to fountain tips, active enhancers may impose a feedback constraint on their own activity through increased polymerase collisions.

Beyond local interference, more global changes in chromatin architecture may also contribute to transcriptional upregulation upon cohesin removal. In mammalian systems, cohesin depletion has been shown to increase spatial mixing between accessible chromatin domains^57^. Similarly, in *C. elegans*, fountains collapse following cohesin^COH-1^ cleavage, with fountain trunks forming increased contacts with surrounding genomic regions (Fig. 4g). In this context, active enhancer-containing fountain tips may become more mobile, gaining the ability to form new regulatory interactions. This could enhance transcription by promoter *trans*-acting enhancer-promoter contacts and increasing regulatory crosstalk.

### Cohesin-mediated gene regulation is evolutionarily conserved

Our data reveal that genes overlapping fountain tips exhibit greater structural complexity, characterized by multiple TSS, TES and/or alternative exons - hallmarks of greater regulatory complexity typically associated with enhancers. The integrity of cohesin^COH-1^ and the presence of fountains thus emerge as critical factors in regulating such complex genes, as illustrated by the isoform switch observed for *skn-1* (Fig. 5). Remarkably, our investigation revealed a noteworthy upregulation of genes associated with neuronal function following cohesin^COH-1^ cleavage (Fig. S10). This conserved role of cohesin in interphase neuronal gene regulation transcends species boundaries. In *Drosophila*, cleavage of the cohesin kleisin in post-mitotic neurons results in impaired axonal and dendritic pruning, as well as abnormal larval locomotion^58^. Similarly, in humans, mutations in cohesin or cohesin-associated genes have been implicated in cohesinopathies, a group of neurodevelopmental disorders^59^. The most well-known of these conditions is Cornelia de Lange syndrome (CdLS), associated with mutations in cohesin subunit NIPBL or other SMC complex subunits^60,61^.

In nematodes, cleavage of COH-1 and subsequent loss of cohesin^COH-1^ produced a spectrum of behavioral phenotypes (Fig. 6,S14). Notably, in the absence of food, the animals exhibited altered foraging behavior, displaying sluggish movement instead of active foraging (Fig. 5). This finding is reminiscent of genetic screens in nematodes that identified mutations in the *mau-2* gene (MAternally affected Uncoordination (*mau*)). *mau-2* mutants display defects in axon guidance leading to behavioral phenotypes^62^ and the gene gave its name to the mice and human Scc4/MAU2 gene necessary for cohesin function. Our study strongly suggests that the observed *mau-2* phenotypes in nematodes are a consequence of cohesin^COH-1^ malfunction in neurons. Similarly, the global dysregulation of genes observed in human CdLS patients further supports the connection to cohesinopathies^63^. Comparison of genes modulated in post-mortem brain neurons of CdLS patients with transcripts modulated upon cohesin^COH-1^ cleavage in nematodes using gene set enrichment analysis, revealed a significant overlap for upregulated genes between the two species (Fig. S15;ref. ^63^). The nematode orthologs of genes upregulated in CdLS were significantly enriched among genes upregulated upon COH-1 cleavage (Fig. S15c), whereas the orthologs of downregulated transcripts in CdLS patients did not display the same pattern (Fig. S15b). Collectively, these similarities strongly indicate that cohesin plays a role in regulating gene expression levels in neurons, most likely through the formation of 3D insulating structures. Further comprehensive studies are required to systematically characterize dysregulated neuronal genes and assess neuronal function upon cohesin^COH-1^ cleavage in *C. elegans*.

In summary, our study provides compelling evidence establishing cohesin^COH-1^ as the key interphase cohesin in *C. elegans*, with a crucial involvement in neuronal function through the formation of insulating 3D structures at active enhancers. Leveraging the extensive repertoire of genetic tools available in nematodes, further exploration of cohesin^COH-1^ holds great promise as a genetically tractable model for investigating human cohesinopathies.

## Acknowledgements

We would like to thank Cihan Elcin, Laurence Bulliard et Lisa Schild for technical help, Dr. Pamela Nicholson and her team at the NGS platform of the University of Bern, the Meister laboratory for discussion. Some data were acquired on machines supported by the Microscopy Imaging Center (MIC) of the University of Bern. Some strains were provided by the CGC, which is funded by NIH Office of Research Infrastructure Programs (P40 OD010440). This work was funded by the Swiss National Science Foundation 31003A_176226/310030_212472 (to PM), PCEFP3_181204 (to BT), 10030_197607, BSSGI0_155764 and PP00P3_150681(to DAG), the University of Bern and Fribourg, the Agence Nationale pour la Recherche (ANR-18-CE45-0022-01 to DJ), the Wellcome Trust (to NG) and the Novartis Foundation for Medical/Biological Research (to PM).

## Methods

### General worm growth and collection

A large unsynchronized worm population of the genotype of interest was grown on four 140 mm peptone-rich plates for about three days to get gravid adults. The worms were bleached, and the eggs hatched overnight in the M9 buffer without food. The synchronized L1s were plated on four 140 mm NGM plates (80,000 – 100,000 worms per plate) and left to grow at 22 ℃ for 24 hours. For specific conditions requiring TEV expression, synchronized L1 animals were left on NGM food for 3 hours, before heat-shocking them at 34°C for 30 minutes. After 19 hours, the L3s were washed from the plates using M9 and did a couple of washes to remove bacteria before proceeding with downstream experiments. For the genomic DNA experiments, the L3s were resuspended in 20 ml of ice-cold M9 and 20 ml of ice-cold 60 % sucrose, shaken vigorously, and gently layered 4 ml of ice-cold M9 on top. 30 ml of the supernatant (on top of the sucrose), now containing floating worms, is aspirated and distributed into 50 ml falcon tubes that were subsequently filled to 50 ml with ice-cold M9. Spun at 1000 rpm for 1 minute at room temperature, discarded the liquid, and washed the worms with ice-cold M9 and once with room-temperature M9. The worms were left at room temperature for 25 minutes for the remaining bacteria in the worms’ gut to get digested, washed once with M9, and removed liquid before proceeding to DNA isolation. For Hi-C, RNA-seq and behavioral data presented here (Fig. 1, 2, 3f, 4, 5), we used heat-shocked animals expressing the TEV protease in the absence of TEV cleavage site as controls (labeled TEV control).

### RNA-seq

RNA sequencing libraries were made by Novogene before sequencing using Illumina NovaSeq (paired-end (PE), 150 bp read length). Reads were aligned to the WS285 transcriptome with Salmon (v.1.9.0) in quant mode with the flags --validateMappings--seqBias --gcBias. Counts per gene were compared with DESeq2 (v1.36.0). GO term enrichment was performed using command line versions of the Wormbase TEA tool ^29^ and WormCat ^65^. The COH-1^cs^ RNAseq data was taken from ^9^ but reanalysed on its own comparing just the COH-1^cs^ (PMW828) strain to the TEV-only control strain (PMW366), aligned to the WS285 transcriptome and pre-filtered before DESeq2 analysis to only include samples with at least 10 reads in half the samples. The different pipeline yielded slightly different results tables, but correlation of the shrunken log2 fold change was 0.97. Of a total 9180 genes left after filtering out genes that oscillate during development ^66,67^, 895 were significantly up regulated and 993 down regulated (adjP<0.05), but as noted previously^9^, most of the log2 fold changes were extremely small, with only 98 genes having a LFC > 0.5 and 52 genes with a LFC < -0.5. Transcript level quantification was carried out by mapping reads to transcripts using Salmon as described above, but adding 100 bootstrap samples. The counts were converted to hd5 format with Wasabi (version 1.0.1) and differential expression was carried out with Sleuth (version 0.30.1), filtering out low count transcripts with the default basic_filter, and also removing transcripts from oscillating and non-protein-coding genes by their ids. Scripts used to produce some of the plots for this paper can be found at https://github.com/CellFateNucOrg/Luthi_etal.

### Nuclei Isolation

400,000 L3s worms were washed with cold nuclear isolation buffer (NIB) (250 mM sucrose, 10 mM Tris-HCl (pH 7.9), 10 mM MgCl_2_, 1 mM EGTA, 0.25 % NP-40, 1 mM DTT, and protease inhibitors). Spun for 1 minute in a microcentrifuge, discarded the supernatant, and then snap-freeze the reaction tube in liquid nitrogen. The frozen worm pellet was squeezed into a pre-chilled mortar, placed in a ceramic bowl, put the pestle in it, and hit five times with a hammer. Removed the pestle and ground it into powder form with an electric drill. The powder was scraped from the mortar into a new tube. An equal volume of NIB was added to the powder, and the fully thawed mixture was transferred into a Kontes™ 2ml glass dounce on ice and centrifuged at 4℃. Dounced the ground-up worms ten times with the “loose” pestle, then ten times with the “tight” pestle, after which it was transferred to a 1.5 ml Eppendorf tube and centrifuged for five minutes at 100 rcf in a microcentrifuge. The supernatant was transferred to a new Eppendorf tube, making sure not to take up any worm debris. Added an equal volume of NIB again and repeated the douncing and spinning four more times till all nuclei were collected. The nuclei are then counted using a hemocytometer and fluorescent microscope.

### bTMP-Seq

bTMP was synthesized as described ^26^ and supplied by Nick Gilbert’s research group. Washed nuclei with wash buffer (WB) (10 mM Tris (pH 7.4), 10 mM NaCl, 3 mM MgCl_2,_ and 0.1 mM EDTA) to remove leftover NIB. Centrifuged for five minutes at 3000 rpm at 4 ℃ in a microcentrifuge and discarded the supernatant. Making sure to work with 10 million nuclei per 200 µl volume wash buffer, added 4 µl of 30 mg/ml stock solution of bTMP (i.e., 600 µg/ml final concentration) to the sample. The reaction mixture was incubated in the dark at room temperature for 30 minutes in an Eppendorf tube on a rotator. Transferred the sample into a 96-well plate to increase the surface area and crosslinked for 10 minutes (8000 energy). Transferred nuclei to a 1.5 ml Eppendorf tube and spun down at 3000 rpm at 4 ℃ in a microcentrifuge for five minutes. The supernatant was discarded, and the volume was made up to 600 µl with WB and incubated with 200 µg/ml proteinase K (12 µl of 20 mg/ml stock) at 55 ℃ for 16 hours (overnight). DNA was purified by phenol:chloroform extraction and isopropanol precipitation as follows: Added an equal volume of phenol:chloroform to the sample and transferred it to a 2 ml PLG tube and mixed thoroughly by inversion for 1 minute. Spun in a microcentrifuge at room temperature to separate the phases at maximum speed and transferred the top aqueous phase to a new 1.5 ml Eppendorf tube. The phenol:chloroform extraction was repeated once. Added 0.6 volumes of 2-Propanol, mixed by inverting the tube once, and spun at maximum speed for 30 minutes at room temperature in a microcentrifuge. All the 2-Propanol was discarded by decanting, washed the pellet twice with freshly prepared 70 % ethanol, and spun for 1 minute at room temperature at maximum speed between the washes. After the last wash, removed all traces of ethanol by inverting the tube and allowed to dry for approximately 10 minutes. DNA was resuspended in 41 µl 1x TE buffer (10 mM Tris (pH7.4), 1 mM EDTA) and quantified by QuantiFluor® dsDNA System (Promega). The DNA (1-2 µg DNA / 100 µl volume) was fragmented into approximately 400 bp fragments using the Bioruptor® sonication system (diagenode) at 30 seconds ON and 90 seconds OFF at low power setting for 12 minutes. For the immunoprecipitation (IP), 20 ng of the DNA sample was collected as input and stored at -20 ℃. The rest of the sample was made up to 1 ml with 1x TE buffer. In a 1.5 ml Eppendorf, 1 ml 1x TE buffer and 50 µl avidin conjugated to magnetic beads (Dynabeads™ MyOne™ Streptavidin C1) were added. The mix was rotated at room temperature for 5 minutes and separated from the beads from the wash on a magnetic stand, after which the clear liquid was discarded; this wash step was repeated twice with 1x TE buffer. Added the 1 ml DNA sample to the washed beads and rotated overnight at 4 ℃. Placed the reaction tube on a magnetic stand and removed and discarded the clear liquid. Washed the DNA-bound beads with 1 ml of TSE I (20 mM Tris-HCl pH 8.1, 2 mM EDTA, 150 mM NaCl, 1 % Triton X-100, and 0.1 % SDS) and mixed by rotating at room temperature for 5 minutes, separated the beads from the wash on a magnetic stand after which the clear liquid was discarded. Subsequent washes were done with TSE II (20 mM Tris-HCl pH 8.1, 2 mM EDTA, 500 mM NaCl, 1 % Triton X-100, and 0.1 % SDS), TSE III (20 mM Tris, pH 8.1, 0.25 M LiCl, 1 mM EDTA, 1 % NP-40 and 1 % deoxycholate (DOC)) and 1x TE buffer as done for TSE I. The IP DNA was eluted by adding a 50 µl elution mix (10 mM EDTA and 95 % formamide) and heated at 95 ℃ for 10 minutes, vortexing the sample every 2 to 3 minutes. Placed the reaction tube on a magnetic stand and transferred the clear elute to a new 1.5 Eppendorf tube. Added 150 µl Milli-Q water to make up to 200 µl and purified the DNA according to the SPRIselect User Guide for SPRI-based size selection. 20 µl of Milli-Q water preheated to 70 ℃ was used for the final DNA elution, and transferred the elute to a new tube and quantified the DNA by Qubit. The Illumina library was then made according to the NEBNext® Ultra™ II DNA library Prep kit for illumina® instruction manual and followed the standard protocol. The libraries were then sequenced at the Next Generation Sequencing (NGS) platform of the University of Bern on Illumina NovaSeq 6000 (single end (SE) 50 bp read length). The sequence data can be found in the supplementary table.

### bTMP-seq data analysis

The data generated were preprocessed as follows: Adapter was trimmed using Trim Galore! Version 0.6.6, after which the reads were aligned to the *C. elegans* reference genome version ce11 using Burrows-Wheeler Aligner (BWA) version 0.7.17. The aligned reads were then sorted based on their genomic coordinates using Samtools version 1.8. Sequencing duplicate reads were removed using Picard version 2.21.8, after which blacklisted genes regions and multi-mapping reads were removed using Samtools. For each control and treatment replicates per experiment, the reads were subsampled to the lowest number of reads (>= 5 million reads), after which the read length was elongated to fragment length by extending reads to 200 bp from the start. The coverage was normalized to the number of million reads (RPM) by adding mean coverage. The IP data was then normalized to input data using the ratio operation in deepTools version 3.5.1. Next, the input normalized data were normalized to input normalized genomic DNA data using the ratio operation in deepTools. To make a direct comparison across all experiments, the data were Z-Score normalized using a custom R script (R version 3.6.1). Replicates were averaged using WiggleTools version 1.2.2. The profile plots were then made using deepTools.

### Public datasets analysis (ChIP-seq, ATAC-seq, enhancer types)

All normalized publicly available ChIP-seq and ChIP-chip data sets were lifted to the *C. elegans* reference genome version ce11, after which the replicates were averaged, and profiles were plotted using deepTools. For the segmentation of enhancer types from Jänes et al., 2018 into active or repressed enhancers and H3K27me3-enriched regions, we used the ChromHMM chromatin states from Daugherty et al. 2017.

### Hi-C analysis

Data from ^24,49^ was processed using HiC-Pro at 2kb resolution and converted into balanced cool and mcool formats using cooler^68^. Hi-C data pileups and corresponding figures were created using the ARA function of GENOVA in R ^69^. To account for the difference in resolution between Hi-C data (2 kb) and enhancer/TSS/HOT sites mappings (1 bp), the genome was tiled into 2 kb bins. Enhancer locations with their annotation (active/repressed/H3K27me3-covered) were taken from ^6,7^, lifted to ce11 and mapped to 2 kb bins. Consecutive or overlapping bins of the same type were merged before performing pileups. The overlap between non-identical type enhancers was minimal, with only 8, 98 and 34 active/H3K27me3-covered, active/repressed and repressed/H3K27me3-covered bins, respectively, out of 2130, 1586 and 1483 active, H3K27me3-covered and repressed bins, respectively. The same approach was taken for the ^6^ enhancer list, except that the enhancer type was determined by mapping the enhancer locations to the ChromHMM chromatin states from ^23^. To calculate an expected value for the average contact frequency maps, the pileups were calculated with a shift of 20’000 kb (10 bins) compared to the bin of interest containing the enhancer, and removal of the first and last percentile (0.01-0.99) of the observed/expected ratio values. Pileups were displayed using the “visualise” function of GENOVA. For genome-wide Hi-C data, contact maps were loaded onto HiGlass for data exploration and ratio map creation ^70^.

### Inferring fountain positions, lengths and asymmetries

We base our fountain detection algorithm on a standard method used in computer vision: a mask representing an idealized version of the object or motif to detect inside an image is locally convoluted to the image. To each pixel of the image a score of object localization is assigned and maxima or peak detection algorithm is then applied to find the pixels where the motifs are localized. In the context of fountain detection, the image is a Hi-C related matrix and the mask is representing a fountain.

More concretely, to focus on cohesin-dependent fountains (*i.e.* to exclude other motifs that may generate false positive detection and to detect fountains that may be hindered inside other strong cohesin-independent motifs), we used the matrix *M* of log10 ratio between the TEV control and cohesin^COH-1^-cleaved Hi-C maps as an input image (Fig. 2F). Based on a simple polymer model of loop extrusion, we defined an idealized fountain mask *F(α_-_,α_+_)* where *α_-_* and *α_+_* are related to the left and right extrusion speeds respectively and control the extent and asymmetry of the fountain (Fig. S3). First, we considered a symmetric mask *α_-_* = *α_+_* = *α_0_*and chose *α_0_* = 0.35 to have a similar fountain size than those observed around enhancers (Fig. 1). This would correspond to an average extrusion speed of ∼4.6/T kbp/min, with T the average residence time in minutes of extruding cohesins on chromatin. We then compute for each position *i* along the genome the element-wise product matrix *fm(i)* between *F(α_0_,α_0_)* and *M* centered around *i*. We define the fountain score *S(i)* as the mean value of the *fm(i)* elements (Fig. S1a-c). We used the *find_peak* function of python package *scipy* to detect the peaks in the *S(i)* profile and compute their prominence *P(i)*. These peaks represent putative fountain positions. To select statistically significant peaks, we generate a null model for each chromosome by computing several random matrices by randomly shuffling the subdiagonals of the original matrix *M*. By repeating the fountain score and peak detection operations on these matrices, we obtain a list of ‘random’ prominences as an empirical null model. Peaks obtained from *M* with a prominence *P(i)* larger corresponding to a p-value of the null model lower than 0.05 were then selected as significant. To detect for possible asymmetry in fountain shapes that may alter the inference of the exact fountain position, we finally scanned around the approximated fountain position for the presence of asymmetric fountains by computing fountain scores for various asymmetric binarized masks *F(α_-_, 2α_0_ - α_-_)* with *α_-_* ∊ [0.05:0.05:2α_0_ - 0.05] ranging from left- to right-handed shapes (Fig. S3). From this, we estimate an asymmetry score (∊ [-1,1]) representing the degree of asymmetry of the fountain (<0: left-handed fountain corresponding to faster extrusion upstream of the fountain tip (5’ side of the tip), >0: right-handed fountain corresponding to faster extrusion downstream of the fountain tip (3’ side of the tip), Fig. 2H, S1). For example, the first and fifth quintile of asymmetry score (Fig.2H) have a median score of ∼± 0.4 respectively, corresponding to a difference in extrusion speed of ∼2/T kbp/min between the upstream and downstream regions around the fountain origin. To estimate the length of each fountain, we consider the average value *<M(d)>* of the log-ratio matrix in a band of width *w* and size *d* perpendicular to the main diagonal and centered around the fountain origin. The fountain length *l* is then defined as (*2d_opt_+1)x2 kb* with *d_opt_* the size *d* for which the difference between <M(d)> and the average value of M at distances larger than *d* is maximal (Fig. S1). Note that similar mask-based strategies are also employed by other Hi-C-motif-detection methods such as ChromoSight^71^ or fontanka^15^. Results from our method and those obtained with fontanka are very consistent (Fig. S1d-f). All the details about mask definition, computation of fountain and asymmetry scores as well as estimation of the null model are given in a python *jupyter* notebook given in Supplementary Information and available at https://github.com/physical-biology-of-chromatin.

### Transcription inhibition experiments

Starting with 400,000 L3s in 10 ml M9 in a 50 ml falcon tube, added 200 µg/ml α-amanitin (Sigma-Aldrich) and OP50 that had been inactivated by heating at 65°C for 30 minutes. As a control, incubated the same amount of worms and inactivated OP-50 in the same volume with an equal volume of Milli-Q water. The reaction mixtures were incubated, rotating at room temperature in the dark for five hours. Spun the worms at 1000 rpm for 1 minute; the liquid was discarded, then washed the worms once with M9 and spun down at 1000 rpm for 1 minute. Resuspended the worms in 100 µl M9 with 200 µg/ml α-amanitin (for control, added Milli-Q water instead).

### Fluorescent imaging of nematodes

Bleach-synchronized L1s were grown on OP-50 for 3 hours at 22°C and then heat-shocked at 34°C for 30 minutes to induce TEV expression and COH-1 cleavage. L3 animals were collected 19 hours post-induction and anesthetized using 200 µl of 0.25 M levamisole. Alternatively, worms were grown at 20C, heat shocked 3 hours after seeding L1s, and L3s collected 24h post-induction. No difference in phenotype was observed between these two regimes. Worms were transferred on a 2 % agarose pad and imaged with a Nikon Ti2 microscope equipped with a Yokogawa spinning disk at 25°C. Z-stacks spaced by 0.3 µm covering 15 µm were acquired for GFP (488 nm), and the brightfield channel and maximum projections were merged in Fiji ^72^.

To evaluate the ectopic expression of *skn-1a* in ASI cells, a custom script in python was used to perform a maximum intensity projection of the Z-stacks and determine the 0.5 and 99.5th quantiles of the GFP signal in each of the skn-1::GFP images without any enhancer deletion. The average of these values was then used as lower and upper bounds of the signal for all the Z-projected images in the data set. Files were saved with randomly assigned anonymous names and scored blindly by two separate individuals (see scoring scheme in Fig. S11a). The average of the two scores for each worm was used. To quantitatively assess the SKN-1::GFP signal in ASI nuclei and cell-bodies proximal to the nucleus, the fluorescence signal was segmented using otsu thresholding. Segmented objects were size selected (min=500, max=6000) to remove fluorescence from gut granules or other objects. One image with more than 4 objects (expect to see 1-2 ASI from 1-2 worms) was removed due to excessive background signal. The quality of segmentation was confirmed by manual inspection. The binary segmentation was used to measure mean fluorescence of the ASI nuclei/cells.

### Microchambers preparation and growth analysis

Eggs from bleach synchronized parental culture were transferred into agarose-based arrayed 600 x 600 x 20 μm microchambers ^37^, mounted onto a 3.5 cm gas permeable polymer dish (ibidi). Dishes were placed in a custom-made stage holder of a 22°C temperature-controlled Nikon Ti2 epifluorescence microscope equipped with a Hamamatsu ORCA Flash 4 sCMOS camera. Images were acquired with a 10x 0.45 NA objective every 10 minutes with 10 ms exposure time to minimize motion blur ^73^, refocusing using Nikon’s NIS software autofocus. Three hours after the start of time lapse imaging, the dish holder was removed from the microscope and dishes were subjected to a 30 minutes heat shock at 34°C to activate TEV expression. The dish holder was then mounted back onto the microscope and time lapse imaging restarted 75 minutes after dish removal. For the analysis, animals were selected which had hatched at least 150 minutes before heat-shock induction. Volume trajectories were determined using automated image analysis ^73^ and manually curated for mistakes in molt annotation.

### Behavioral analysis

Behavioral analysis was conducted at 22°C in young adult animals, as previously described ^40^. Briefly, animals were synchronized and, except for N2 controls, submitted to a heat-shock to induce TEV expression as described above. Fed animals were maintained on 6-cm NGM plates with a lawn of OP50 bacteria. Food-deprived animals were deposited on unseeded NGM plates after 3 washes with distilled water. Several hours prior to experiments, plates were transferred into a custom-designed temperature- and vibration-controlled recording system. High resolution (2448×2048 pixels) 3-min movies of worm behavior (∼50 animals/movie) were acquired with a DMK33UX250 camera (The Imaging Source), at 10 frames per second using IC Capture software (The Imaging Source). Movies were analyzed with the Tierpsy tracker (v1.4; ^38^ and we focused on a subset of 47 interpretable parameters, the choice of which was previously discussed ^40^. PCA and hierarchical clustering were performed with Clustvis webtool by default SVDimpute algorithm ^74^. Each condition and genotype were recorded on at least 3 separate days with multiple replicates per day.

### Comparative Gene Set Enrichment Analysis

Human-nematode orthologous genes were downloaded from Ortholist2 ^75^. Only orthologs that were detected by at least two programs were used in the analysis. A table of genes differentially expressed in Cornelia de Lange Syndrome neurons compared to control neurons ^63^ was kindly provided by Matthias Merkenschlager. Genes were ranked by their Log2 fold change after COH-1 cleavage and their ranks used to test for enrichment of worm orthologs of significantly up and down regulated genes (padj<0.05) in CdLS patients using the fgsea package in R.

## Supplementary data

### Worm strains

| Strain name | Genotype |
| --- | --- |
| PMW366 | *ubsSi20[hsp-16.2p::TEV::unc-54 3'UTR; Cbr-unc-119(+)] II; unc-119(ed3) III* |
| PMW828 | *ubsSi20[hsp-16.2p::TEV] II; unc-119(?) III; coh-1(ubs26[1xFLAG TEVcs]) X* |
| PMW836 | *scc-1(ubs19[1xFLAG TEVcs]) ubsSi20[hsp-16.2p::TEV] II; unc-119(?) III; wls54[scm::gfp] V* |
| PMW844 | *scc-1(ubs19[1xFLAG TEVcs]) ubsSi20[hsp-16.2p::TEV] II; unc-119(?) III; coh-1(ubs26[1xFLAG TEVcs]) X* |
| PMW1116 | *coh-1(ubs26[TEVcs]) X; ubsSi20[hsp-16.2p::TEV::unc-54 3'UTR; Cbunc-119] II; ujls113 II; skn-1(hq82([skn-1::gfp]) IV* |
| PMW1117 | *coh-1(ubs26[TEVcs]) X; ujls113 II; skn-1(hq82([skn-1::gfp]) IV* |
| PMW1063 | *ubsSi20[hsp-16.2p::TEV::unc-54 3'UTR; Cbunc-119]II; xeSi301[Peft-3::luc::gfp::unc-54 3'UTR, unc-119(+)]III; coh-1(ubs26[TEVcs])X* |
| PMW1064 | *ubsSi20[hsp-16.2p::TEV::unc-54 3'UTR; Cbunc-119]II; xeSi301[Peft-3::luc::gfp::unc-54 3'UTR, unc-119(+)]III* |
| PMW1182 | *ubsSi20[hsp-16.2p::TEV::unc-54 3'UTR; Cbunc-119] II; skn-1(hq82(skn-1::gfp)) ubs62(bec-1 enhancer long deletion) IV; coh-1(ubs26[TEVcs]) X.* |
| PMW1217 | *ubsSi20[hsp-16.2p::TEV::unc-54 3'UTR; Cbunc-119] II; skn-1(hq82([skn-1::gfp]) nhr-46(ubs71[deletion]) IV; coh-1(ubs26[TEVcs]) X* |
| PMW1218 | *ubsSi20[hsp-16.2p::TEV::unc-54 3'UTR; Cbunc-119] II;*<br>*skn-1(hq82([skn-1::gfp]) clec-178(ubs72[deletion]) IV;*<br>*coh-1(ubs26[TEVcs]) X* |
| PMW1219 | *ubsSi20[hsp-16.2p::TEV::unc-54 3'UTR; Cbunc-119] II;*<br>*skn-1(hq82([skn-1::gfp] ubs73[very large intron deletion]) IV ;*<br>*coh-1(ubs26[TEVcs]) X* |

### *skn-1* enhancer deletion location

| Deletion | Chromosome number | Start | End |
| --- | --- | --- | --- |
| ubs62 deletion | chrIV | 5665244 | 5667587 |
| ubs71 deletion | chrIV | 5747318 | 5766196 |
| ubs72 deletion | chrIV | 5654185 | 5655138 |
| ubs73 deletion | chrIV | 5655970 | 5659745 |

### Guide RNAs/primers used for deletion

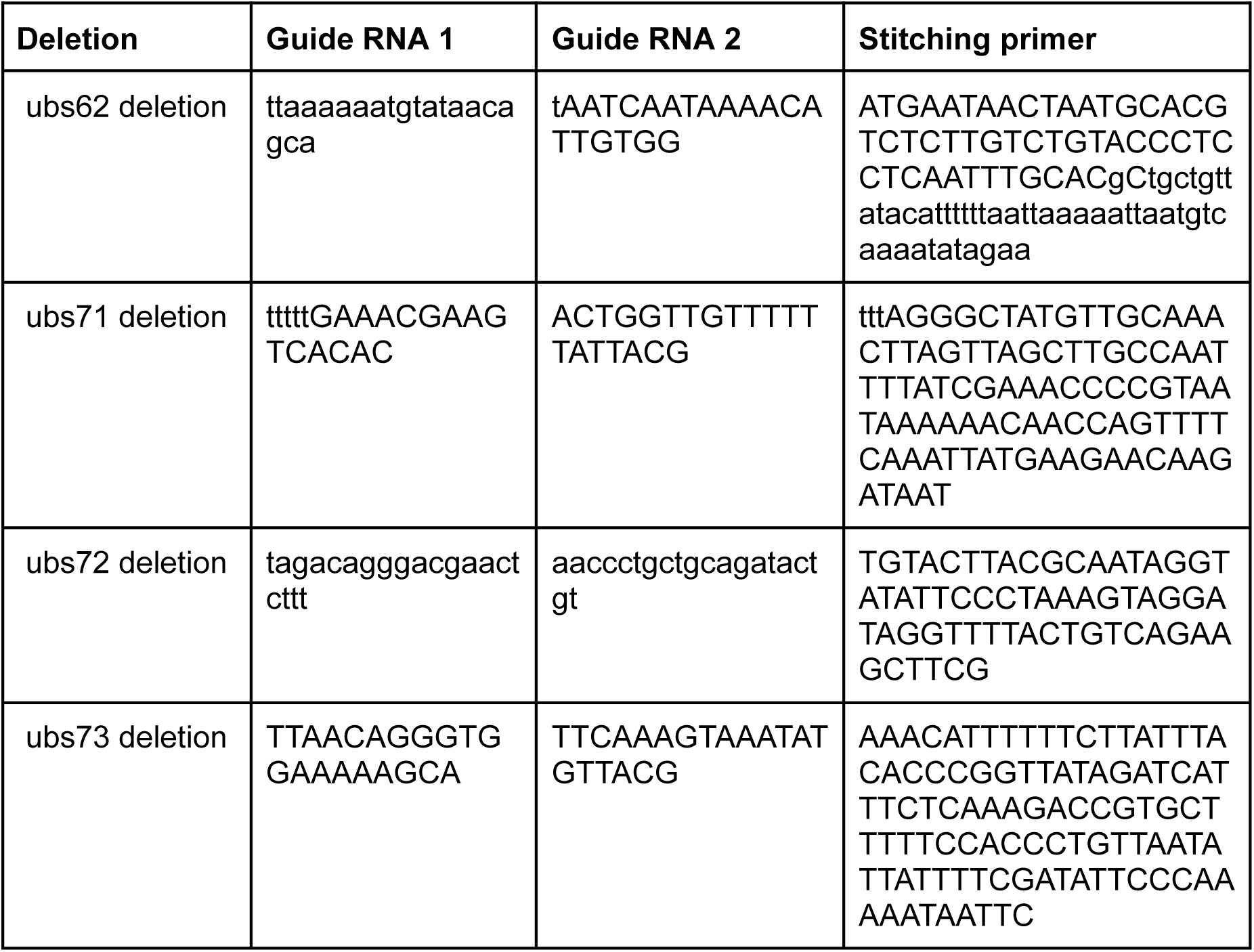

### Public datasets

| Data set | Larval stage | GEO Accession code | Reference |
| --- | --- | --- | --- |
| RNAP II ChIP-seq | L3 | GSE188851 | 24 |
| TOP-1 ChIP-seq | L3 | GSE188851 | 24 |
| TOP-2 ChIP-seq | L3 | GSE188851 | 24 |
| COH-1 ChIP-seq | Young adult | modENCODE project | 76 |
| cohesin<sup>COH-1</sup> cleavage Hi-C | L3 | GSE199723 | 9 |
| Cohesin<sup>SCC-1</sup> cleavage Hi-C | L3 | GSE199723 | 9 |
| TIR1 control Hi-C | L3 | GSE188851 | 24 |
| TOP-1 Hi-C | L3 | GSE188851 | 24 |
| TOP-2 Hi-C | L3 | GSE188851 | 24 |
| cohesin<sup>COH-1</sup> cleavage RNA-seq | L3 | GSE199723 | 9 |
| Cohesin<sup>SCC-1</sup> cleavage RNA-seq | L3 | GSE199723 | 9 |
| Enhancers | L3 | GSE114494 | 6 |
| Enhancers | L3 | GSE89608 | 7 |
| HOT regions | Late embryo | GSE49870 | 16 |

## Supplementary files

**Supplementary file 1.**

Python script for fountain detection on ratio maps. (ipynb format).

**Supplementary file 2.**

Bed format file with detected fountains genome-wide (result of supplementary file 1).

**Supplementary file 3.**

Gallery of the 935 identified fountains, one fountain per row. The first column shows the Hi-C contact map around the identified fountain, upper right: TEV control conditions; lower left: cohesin^COH-1^ cleavage. The second column shows the log-ratio map between cohesin^COH-1^ cleavage and TEV control. The third column shows the Hi-C contact map as in the first column, upper right: cohesin^SCC-1^ cleavage; lower left: cohesin^COH-1^ and cohesin^SCC-1^ simultaneous cleavage. The last column is the log-ratio map between simultaneous cohesin^COH-1^ and cohesin^SCC-1^ cleavage and cohesin^SCC-1^ cleavage. For each contact map, genes are indicated on the upper and left side (horizontal lines) as well as the identified active enhancers^7^ (vertical ticks). The COH-1 ChIP-seq enrichment from modENCODE is depicted on the lowest track on the top of the Hi-C-map, as well as the right-most track on the left side.

---

**Supplementary Figure 1.**
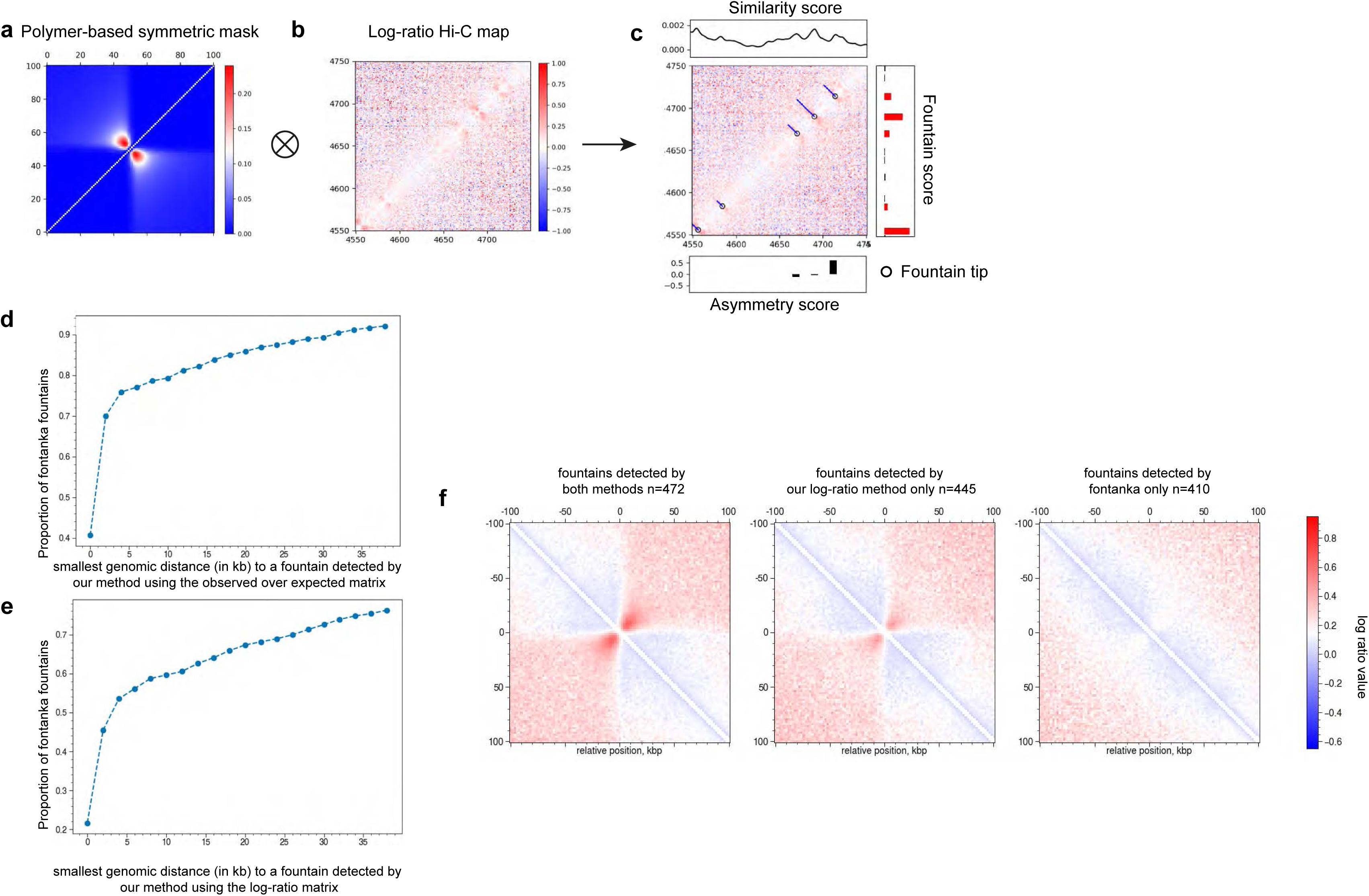
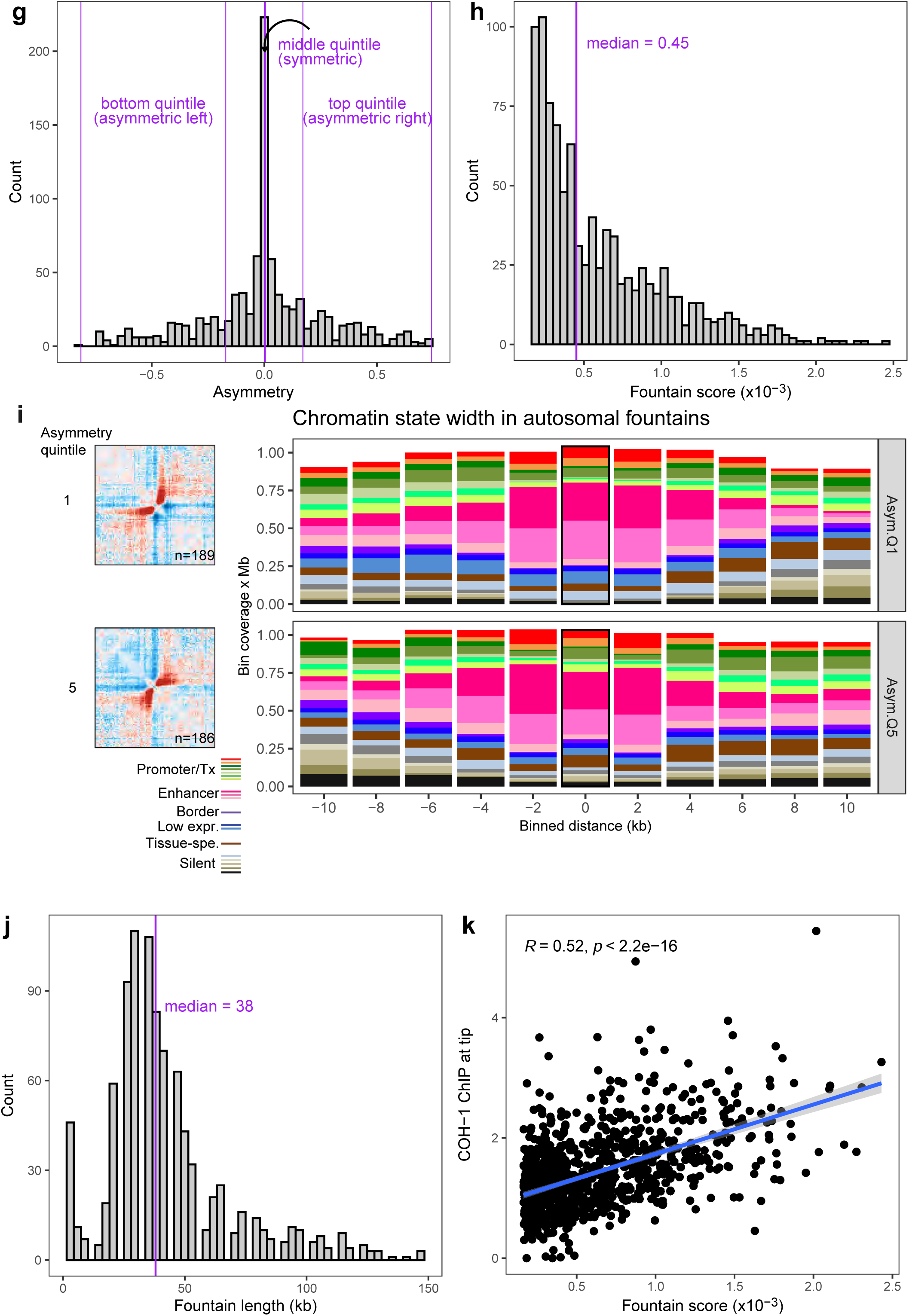
Inferring fountain tip positions and asymmetries. **a.** A polymer-model-based mask of a symmetric fountain is convoluted along the diagonal of the **b.** log-ratio matrix (TEV control divided by cohesin-depleted) to compute a local fountain score. **c.** Maxima of the profile of fountain score (black line in top panel) along the genome with significant prominence (red bars in right panel) are assigned as fountain tips (black circles). Asymmetry in fountain shapes is measured using asymmetric binarized masks around the position of the fountain tip determined previously (<0: left-handed fountain, >0: right-handed fountain, top panel). Fountain lengths (blue segments) are estimated by considering the average value of the log-ratio matrix perpendicular to the main diagonal around fountain origin. **d.** Comparison between our fountain detection method and fontanka^15^. Fountains were detected using fontanka on the TEV control dataset using default parameters and the same symmetric polymer-model-based mask that we developed. Note that fontanka works on the observed-over-expected matrix of a single input Hi-C dataset and thus cannot be run over the log-ratio matrix (TEV control divided by cohesin^COH-1^ cleavage) that we are using to detect fountains. To allow fair comparison with our method, we therefore applied our strategy on the observed-over-expected matrix of the TEV control dataset and compared the inferred positions of fountains. **d.** Percentage of fountains detected by fontanka that are closer than the distance on the x axis from a fountain detected by our method. About 75% of fontanka-detected fountains are also detected (genomic distance less than 5 kbp) by applying our approach to the observed-over-expected TEV control Hi-C map. **e.** Comparison of fontanka-inferred fountains on observed-over-expected TEV control Hi-C map with the positions of the fountains detected by applying our method to the log-ratio matrix (TEV control over cohesin^COH-1^-cleavage). About 55% of fontanka-detected fountains colocalized with log-ratio matrix-fountains. **f.** Average contact frequency maps of the log ratio matrix around fountains detected by both methods (left), only by our approach (center) and only by fontanka (right). Interestingly, fountains detected only by fontanka do not exhibit a clear signature over the log-ratio matrix (right) suggesting that they may represent cohesin-independent motifs, while fountains detected by our method only have a clear signature on the log-ratio matrix (center). **g.** Histogram of fountain asymmetry score. The score was divided into 5 equally sized groups with the lowest and highest quintiles corresponding to asymmetric fountains oriented left and right respectively, and the central quintile (values all around 0) representing symmetric fountains. **h.** Histogram of the fountain prominence score with the median score marked in purple. **i.** Fountain asymmetry is associated with asymmetric enhancer states. Fountain asymmetry scores were divided into five equally sized bins as in g. Left panel: average contact frequency maps in regions around asymmetric fountains from the first and fifth quintiles. Right panel: width in bp of each chromatin states in fountain regions from the highest and lowest asymmetry quintiles (up to 10 kb upstream and downstream of fountain tips. **j.** Histogram of fountains length in kb with the median length marked in purple. **k.** Pearson correlation of the fountain prominence score with the COH-1 ChIP-seq signal measured at the 2 kb fountain tip bin.

**Supplementary Figure 2.**
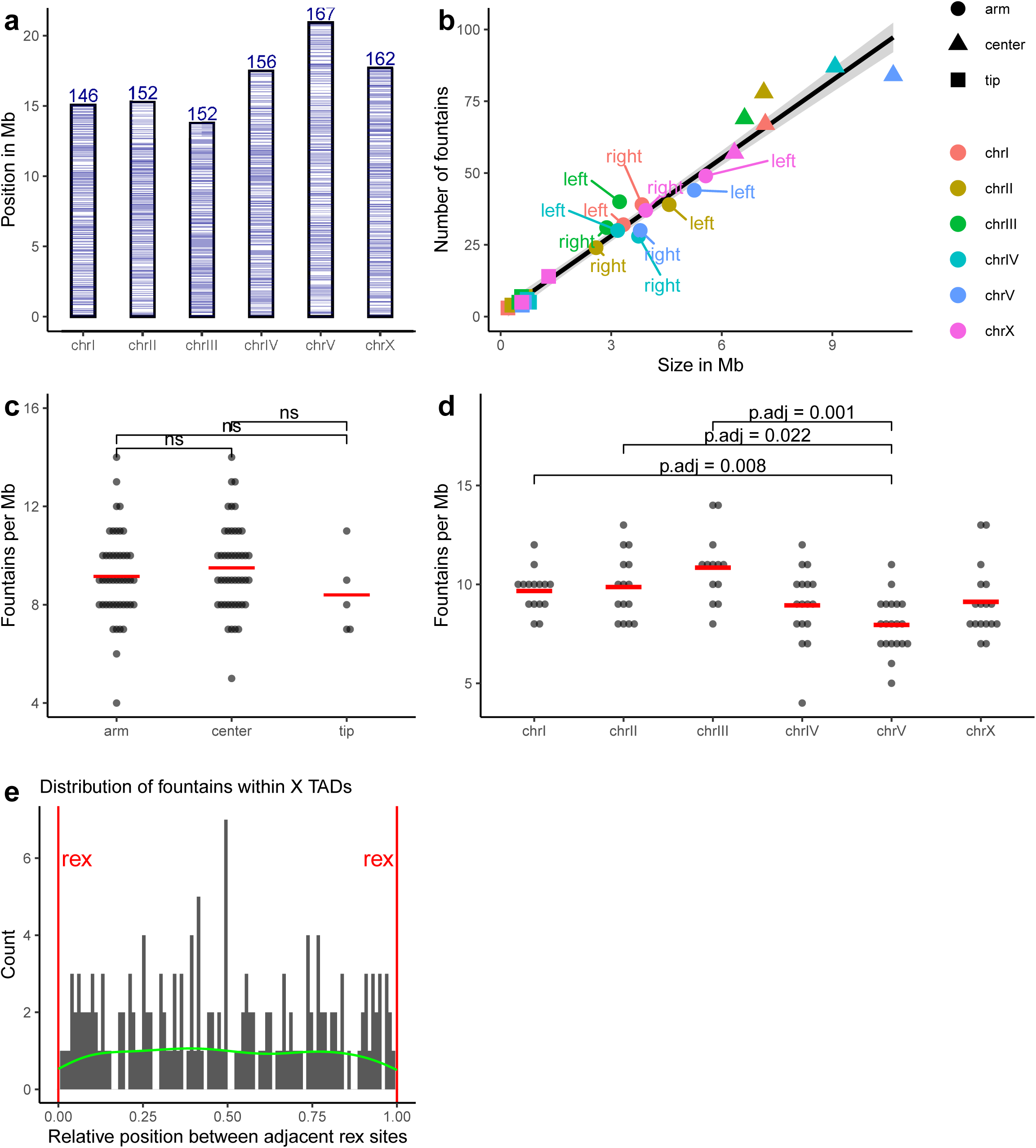
Analysis of the genome-wide distribution of identified fountains. **a.** Location of all the fountains plotted as horizontal blue lines (with transparency) on the chromosomes to provide a visual representation of their distribution. The total number of fountains per chromosome is indicated above the bars **b.** The number of fountains identified per chromosomal region (see ref. ^64^) as a function of the region length. A fitted linear regression line is shown in black. **c. d.** Fountains per 1Mb bin plotted by region type or by chromosome. The mean number of fountains per Mb per group is shown with horizontal red lines. A two sided Wilcoxon rank sum test was used to compare the groups. For clarity, only statistically significant comparisons are indicated with their FDR adjusted p-values in d, while in c the differences were all non-significant (ns). **e.** Histogram of relative position of fountains in-between adjacent *rex* sites (red) shown in gray and their density as a green line.

**Supplementary Figure 3.**
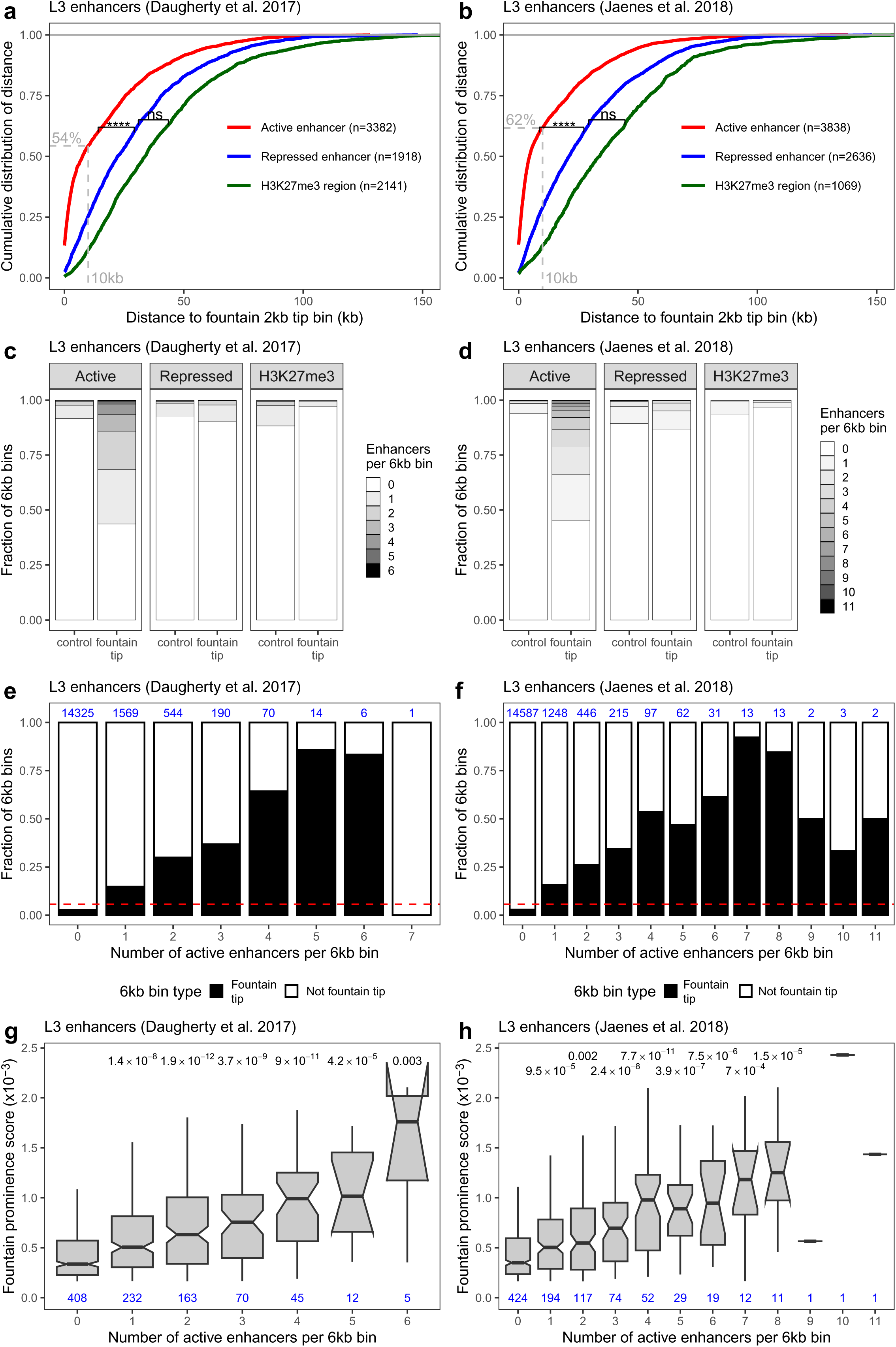
Enhancer location and clustering at fountain tips. **a,b.** Cumulative distribution of the distance of enhancers from the 2 kb fountain tip bin for the different types of enhancers. The percentage of enhancers found within 10 kb of the fountain tip is shown in gray. A one-sided Kolmogorov-Smirnov test was used to compare the curves (**** p value < 0.0001, ns: not significant). **c,d.** Fountain tip bins were resized to 6 kb to include the neighboring bins and the number of enhancers of each type that overlapped these bins was counted. Six kilobase regions located midway between neighboring fountains were used as controls (Fig. 4c). **e,f.** Fountain tips are enriched for clustered active enhancers. The entire genome was divided into 6 kb bins and the number of active enhancers in each bin was counted. The number of 6 kb fountain tip bins containing a given number of active enhancers is shown as a fraction of all genomic 6 kb regions with the indicated number of enhancers (x axis). The total number of genomic 6 kb bins with a given number of enhancers is shown in blue on top. The red line depicts the ratio between the total number of fountain 6 kb bins and the total number of genomic 6kb bins. **g,h** Boxplots of the fountain prominence score of fountain-tip 6kb bins according to the number of enhancers present in that bin. FDR adjusted p-values from a two sided Wilcoxon rank sum test comparing each group to the fountains bins without any enhancers, are shown above the boxplots. **a,c,e,g** plots are for ref. ^7^ L3 enhancers and **b,d,f,h** plots are for ref. ^6^ L3 enhancers where the enhancer type was determined by finding the longest overlap with ChromHMM states from ref. ^7^.

**Supplementary Figure 4.**
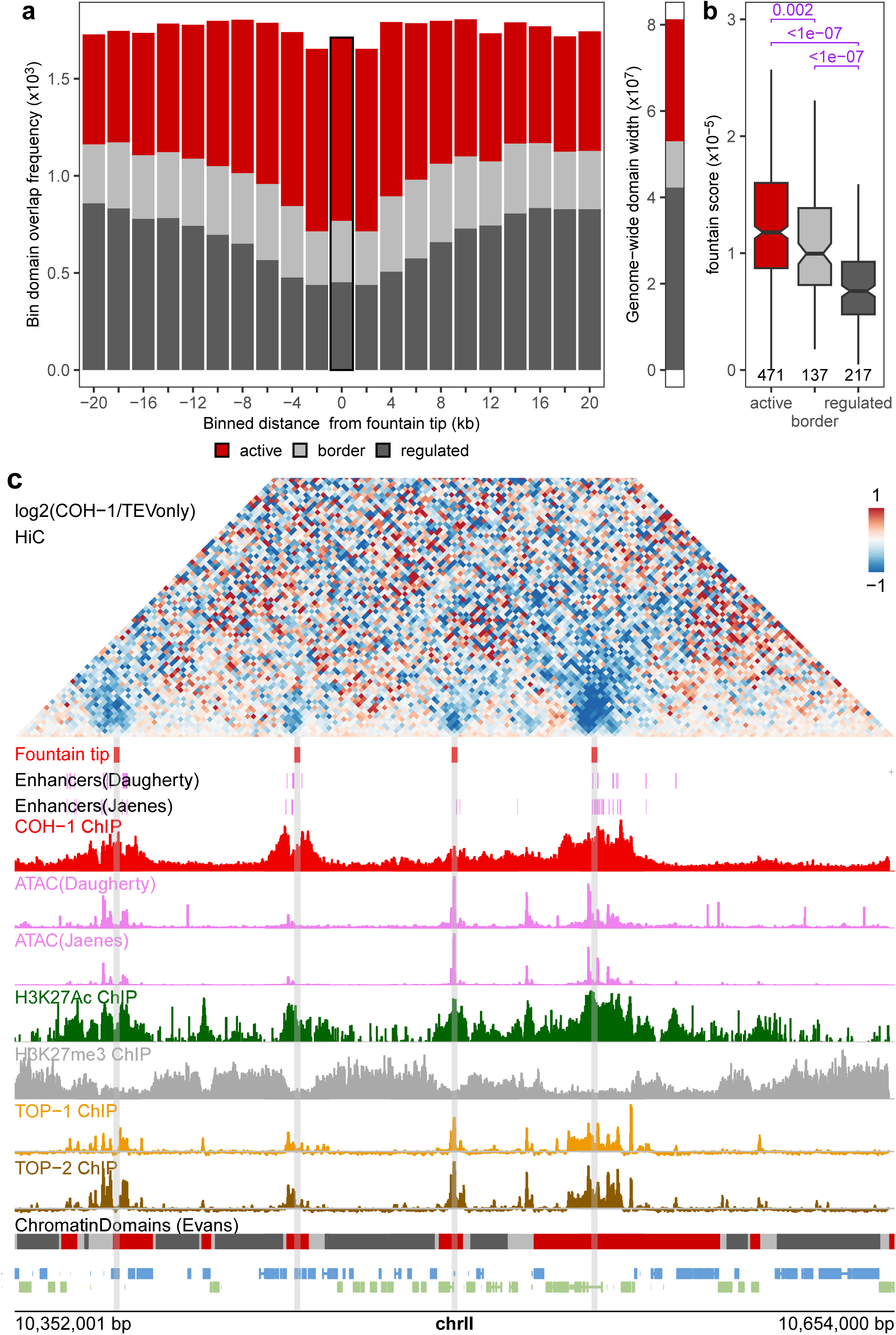
Fountain overlap with chromatin domain types. **a.** Frequency of overlap of different autosomal chromatin domain types^23^ with fountain tip bins and 2 kb bins up to 20 kb upstream and downstream of the fountain tips (left panel), compared to the cumulative genome wide width of these domain types (right panel). **b.** Comparison of the fountain score for fountains that overlap different types of domains. The number of fountains in each group is indicated below the boxplots. Adjusted p-values from Wilcoxon rank sum test are shown in purple. **c.** Example of a 0.3 Mb region on chromosome II containing several fountains that fall within active chromatin domains. Only L3 active enhancers from ^7^ and ^6^ are shown in the enhancer tracks. Chromatin IP data: COH-1 in young adults (GSE50324), H3K27 acetylation (GSM624432) and H3K27me3 in L3 larvae (GSM1206310), TOP-1 (GSM5686806 & GSM5686807) and TOP-2 in L2-L3 larvae (GSM5686812 & GSM5686813), bottom track: gene locations.

**Supplementary Figure 5.**
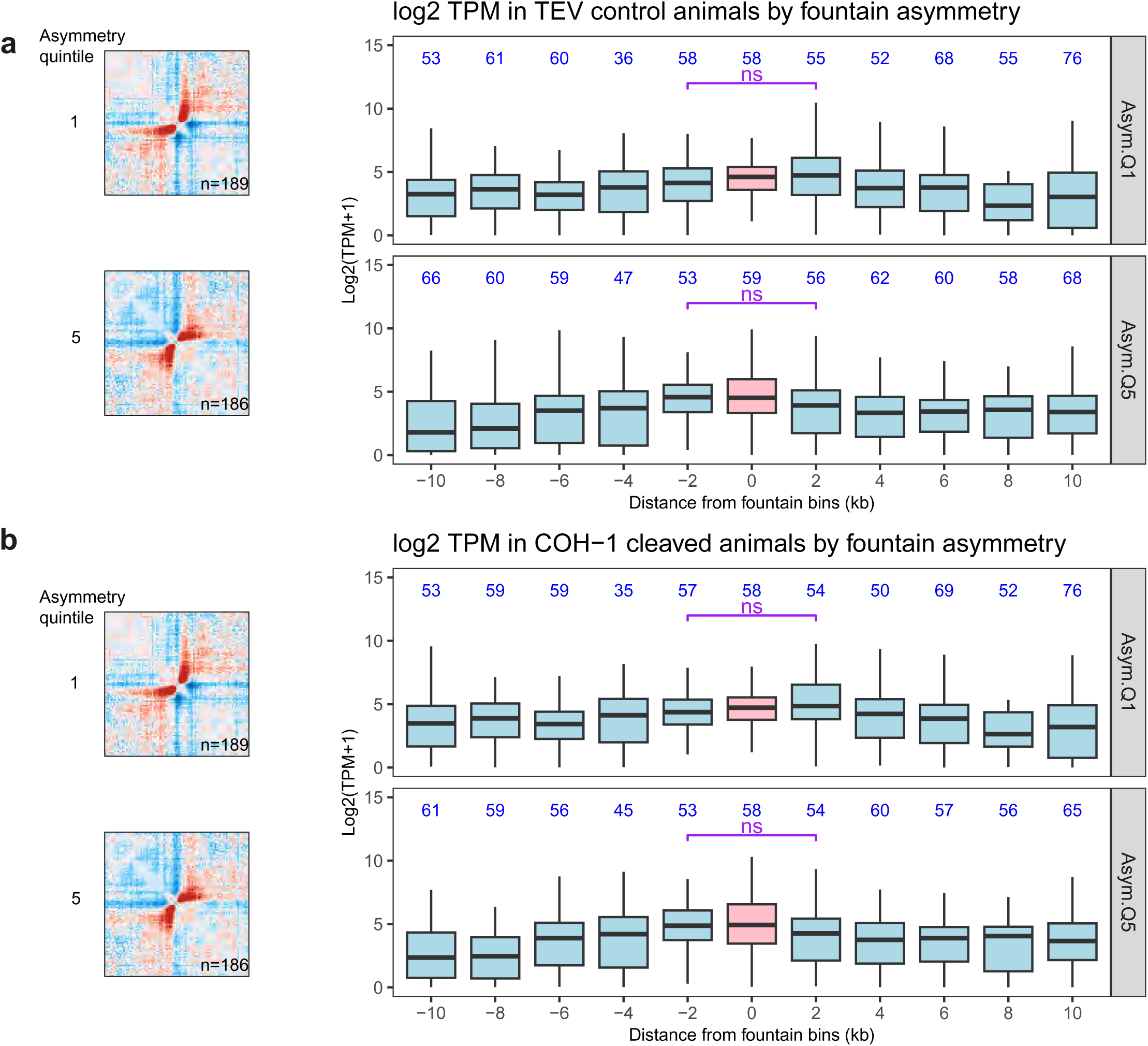
Fountain asymmetry is associated with asymmetry in expression levels. Fountain asymmetry scores were divided into five equally sized bins (quintiles, see Fig S1g). 2 kb bins overlapping the fountain-tips (pink) from the highest and lowest quintiles, and up to 10 kb upstream and downstream (light blue), were used to quantify transcript abundance for genes whose TSS overlaps them. **a.** Left panel: average contact frequency maps in regions around asymmetric fountains from the first and fifth quintiles. Right panel: Boxplot of gene expression (TPM) in TEV-only control samples of genes whose TSSs overlap the respective 2 kb bins around fountain tips. **b.** Left panel as in a. Right panel: Boxplot of gene expression (TPM) in COH-1 cleavage samples of genes whose TSSs overlap the respective 2 kb bins around fountain tips. In both a and b the purple line indicates that a two sided Wilcoxon rank sum test was carried out to compare the expression of bins immediately upstream and downstream of the fountain tip (ns, not significant). The number of genes in each bin are indicated in blue.

**Supplementary Figure 6.**
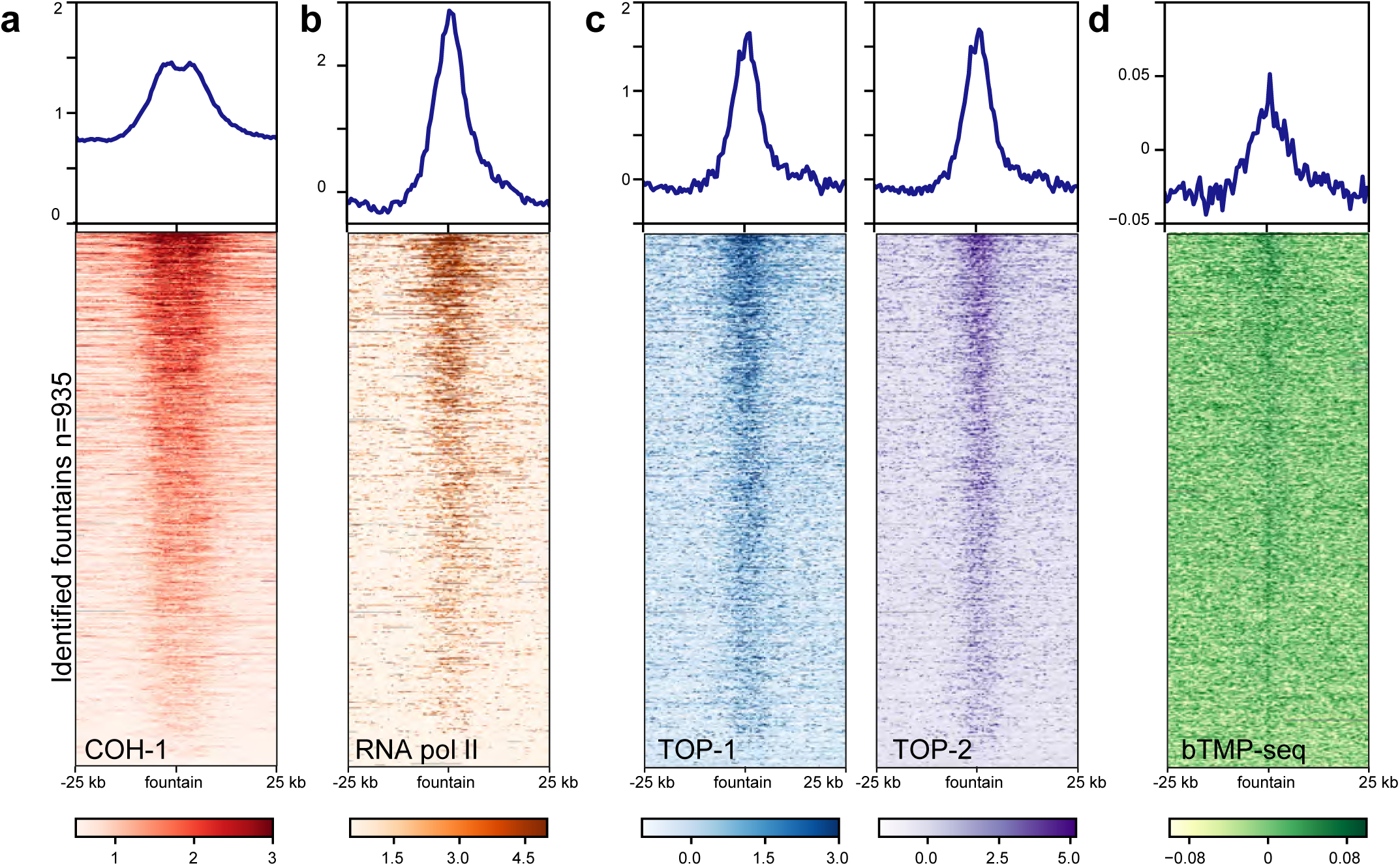
Fountain tips are enriched for COH-1/topoisomerases and bTMP, similar to active enhancers. (top) Average ChIP-seq profiles at fountains as in Fig. 3a-d for **a.** COH-1 (young adults). **b.** RNA pol II (L3), **c.** (left) TOP-1 (L3) and (right) TOP-2 (L3) **d.** bTMP (L3). (bottom) Heatmap of **a.** COH-1, **b.** RNA pol II, **c.** (left) TOP-1 and (right) TOP-2 **d.** bTMP enrichment centered on fountain tips, sorted by COH-1 ChIP-seq enrichment.

**Supplementary Figure 7.**
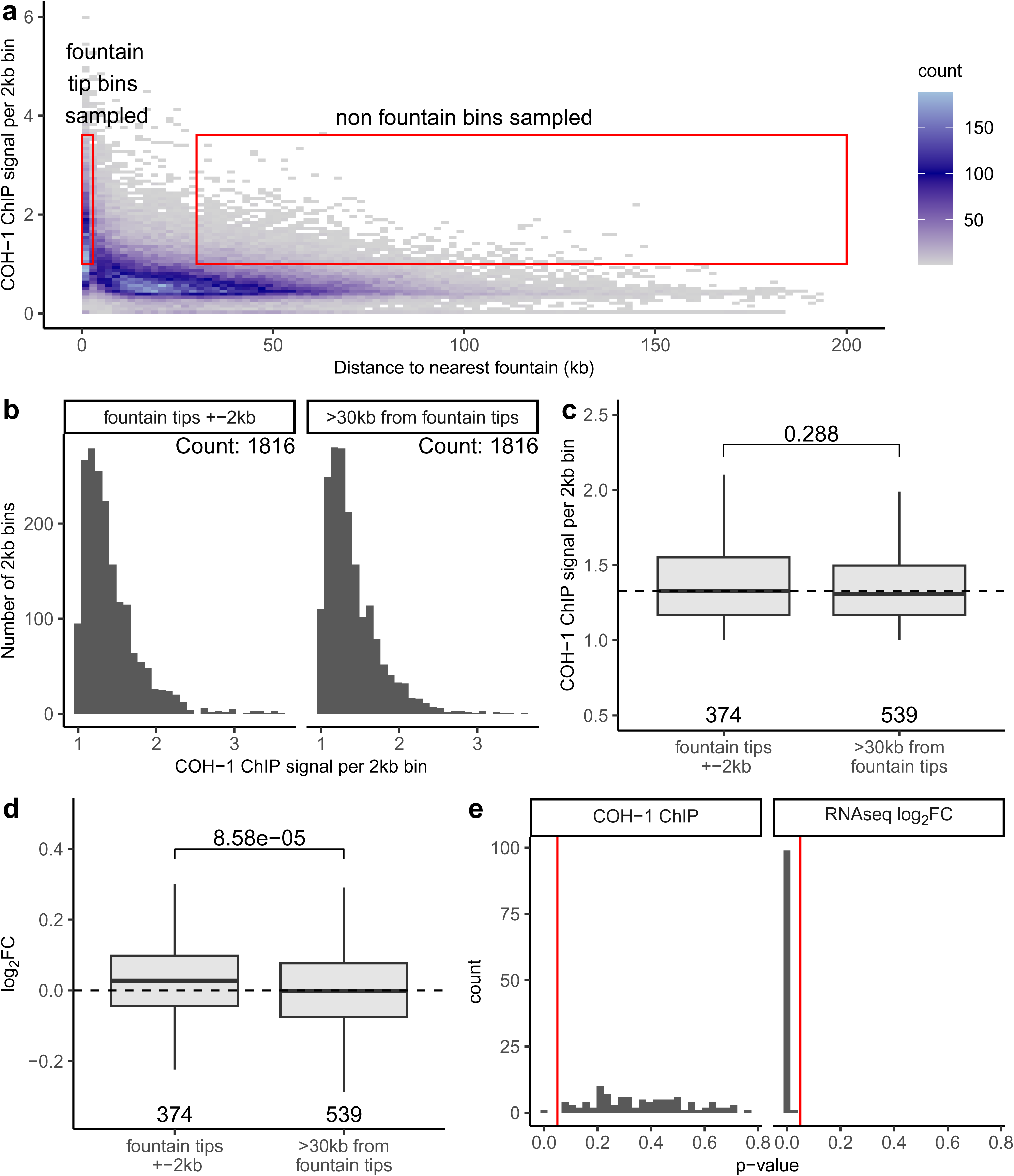
Gene upregulation is specific to fountains and not COH-1 enrichment. **a.** Binned density map of COH-1 ChIP signal by distance from the nearest fountain for all 2kb bins in the genome. To identify bins with matched COH-1 levels, we sampled from fountain tip bins and their immediate neighbours and compared them to bins that were at least 30 kb from the nearest fountain as indicated by the regions enclosed in red boxes. **b.** Histograms of the ChIP signal from two examples COH-1-matched, equally-sized sets of bins from fountain tips and distal regions. The matched sets were created by sampling without replacement from quantiles of COH-1 ChIP signal of the bins indicated in panel a. **c.** Boxplot of the COH-1 ChIP signal of all bins from the matched sets in b, that overlapped a TSS of an expressed gene in the COH-1 cleavage RNAseq dataset. For c and d, the p-value from a one-sided Wilcoxon rank sum test is shown above the boxplots and the number of bins in each group are shown below. **d.** Boxplot of the log2 fold change in gene expression upon COH-1 cleavage of genes whose TSS overlaps bins from the COH-1 matched sets of 2kb bins shown in c. **e.** Histograms of the p-values from one-sided Wilcoxon rank sum tests comparing the COH-1 ChIP signal and RNAseq log2 fold change between fountain tips and regions at least 30 kb from fountains, obtained by repeating 100 times the sampling procedure from the regions shown in panel a, to create matched COH-1 subsets as shown for one example in panels b-d. The vertical red line shows the statistical significance threshold of 0.05.

**Supplementary Figure 8.**
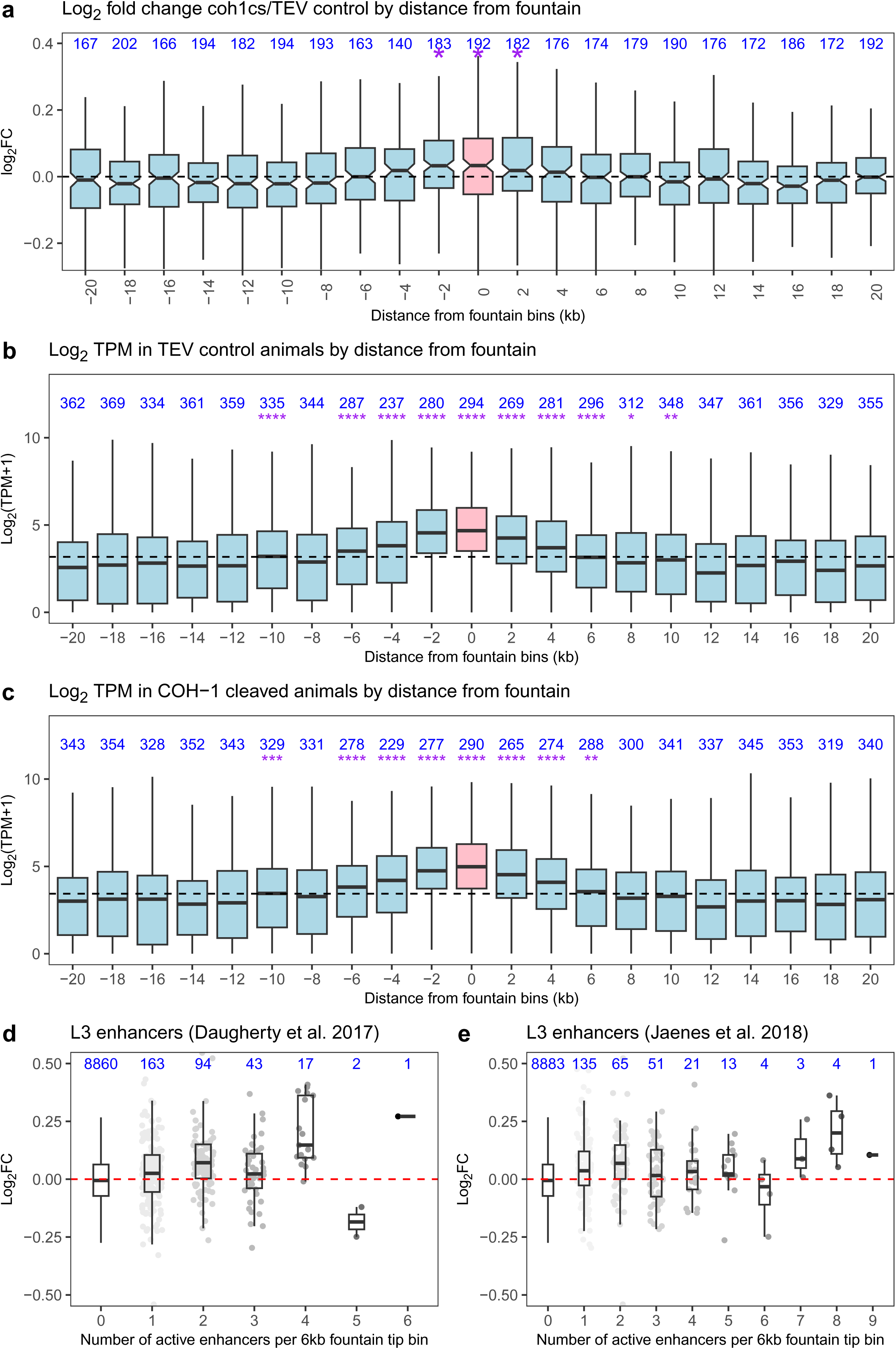
Gene expression around fountain tips. **a-c** 2 kb bins overlapping fountain tips (pink) and up to 20 kb upstream and downstream (light blue), were used to quantify log2 fold change (**a**) and transcript abundance in control (**b**) and upon cohesin^COH-1^ cleavage (**c**) for genes whose TSS overlaps them. **d,e.** Effect of active enhancer clustering on gene expression: Log2 fold change of genes whose TSS overlaps 6kb bins at the fountain tips with different numbers of active enhancers. The number of genes in each group is shown in blue. Individual data points are shaded in gray.

**Supplementary Figure 9.**
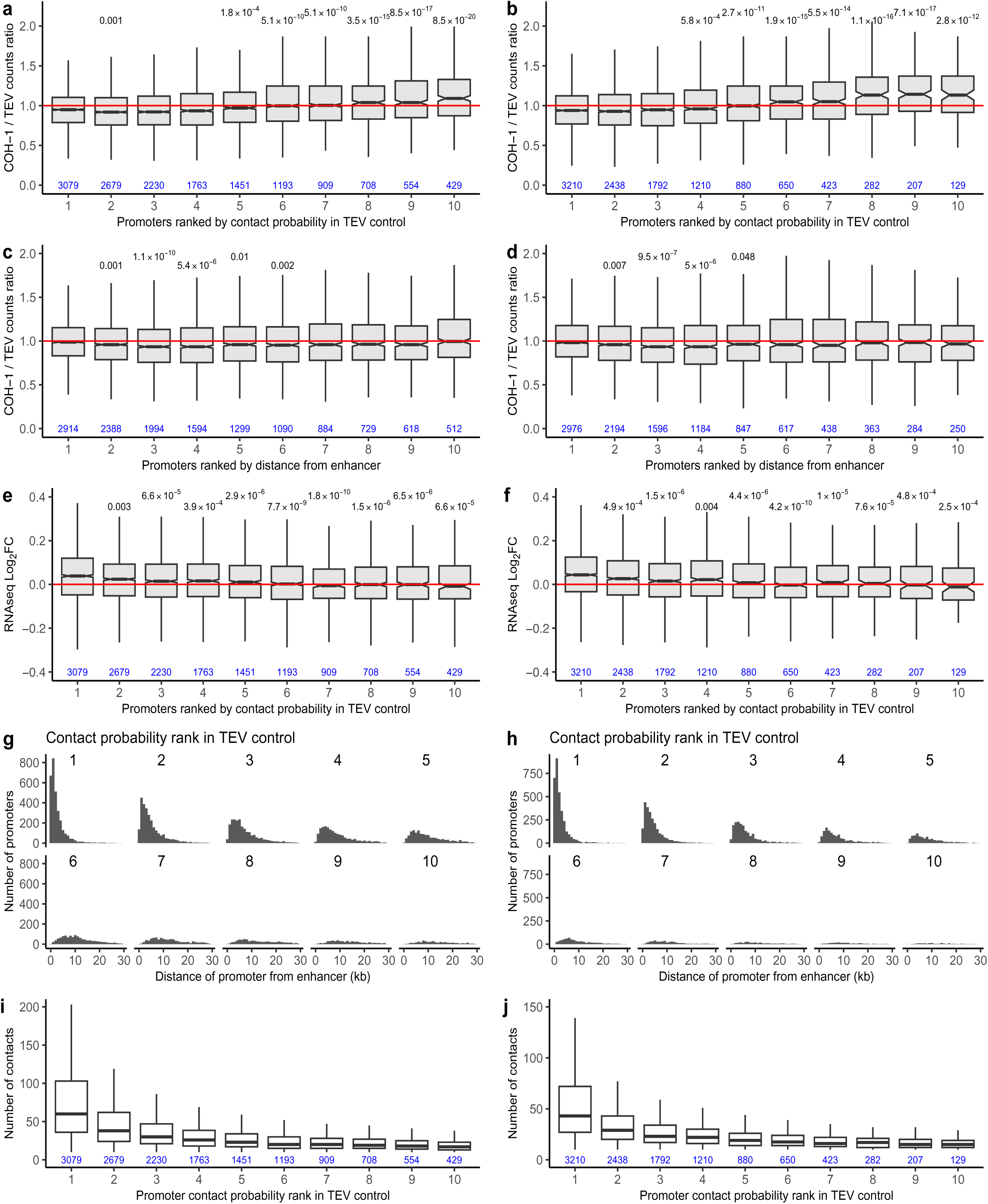
Changes in active enhancer promoter contacts upon COH-1 cleavage and transcriptional consequences on target genes. **a, b.** The ratio of Hi-C fragment contacts from COH-1 cleavage and TEV-only control data for each L3 active enhancer with its closest 10 transcript promoters up to a maximum distance of 30kb. Promoters were ranked by the contact probability with the enhancer in the TEV control Hi-C data. **c, d.** Same as in a, but promoters were ranked by their distance from the active enhancer. **e, f.** RNA-seq log2 fold change between COH-1 cleavage and TEV control for promoters ranked by their contact probability with the enhancer in the TEV control Hi-C data. **g, h.** Histogram of the distances of the closest 10 promoters to the enhancer grouped by their contact probability rank in TEV control Hi-C data. **i, j.** Boxplot of the number of enhancer-promoter contacts grouped by the contact probability of rank of the promoter with the enhancer in TEV control Hi-C data. For a-f, i & j the number of enhancer-promoter pairs in each ranked group is show in blue at the bottom of the boxplot. For a-f the significant FDR adjusted p-values for a two-sided Wilcoxon rank sum test comparing each group to the first are shown at the top. a,c,e,g,i panels are for L3 active enhancers from Daugherty et al. (2017) and b,d,f,h,j panels are for L3 active enhancers from Jaenes et al. (2018).

**Supplementary Figure 10.**
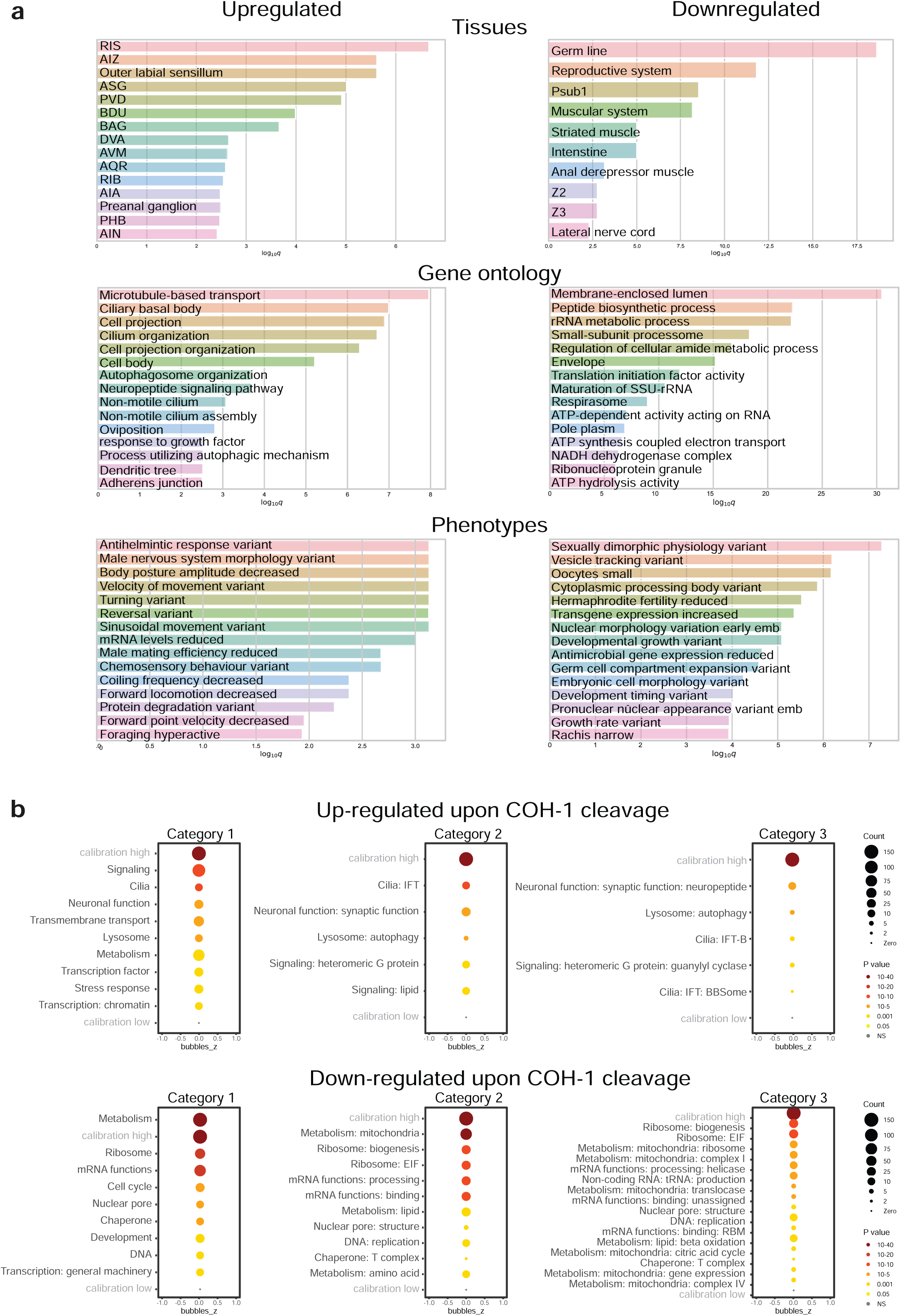
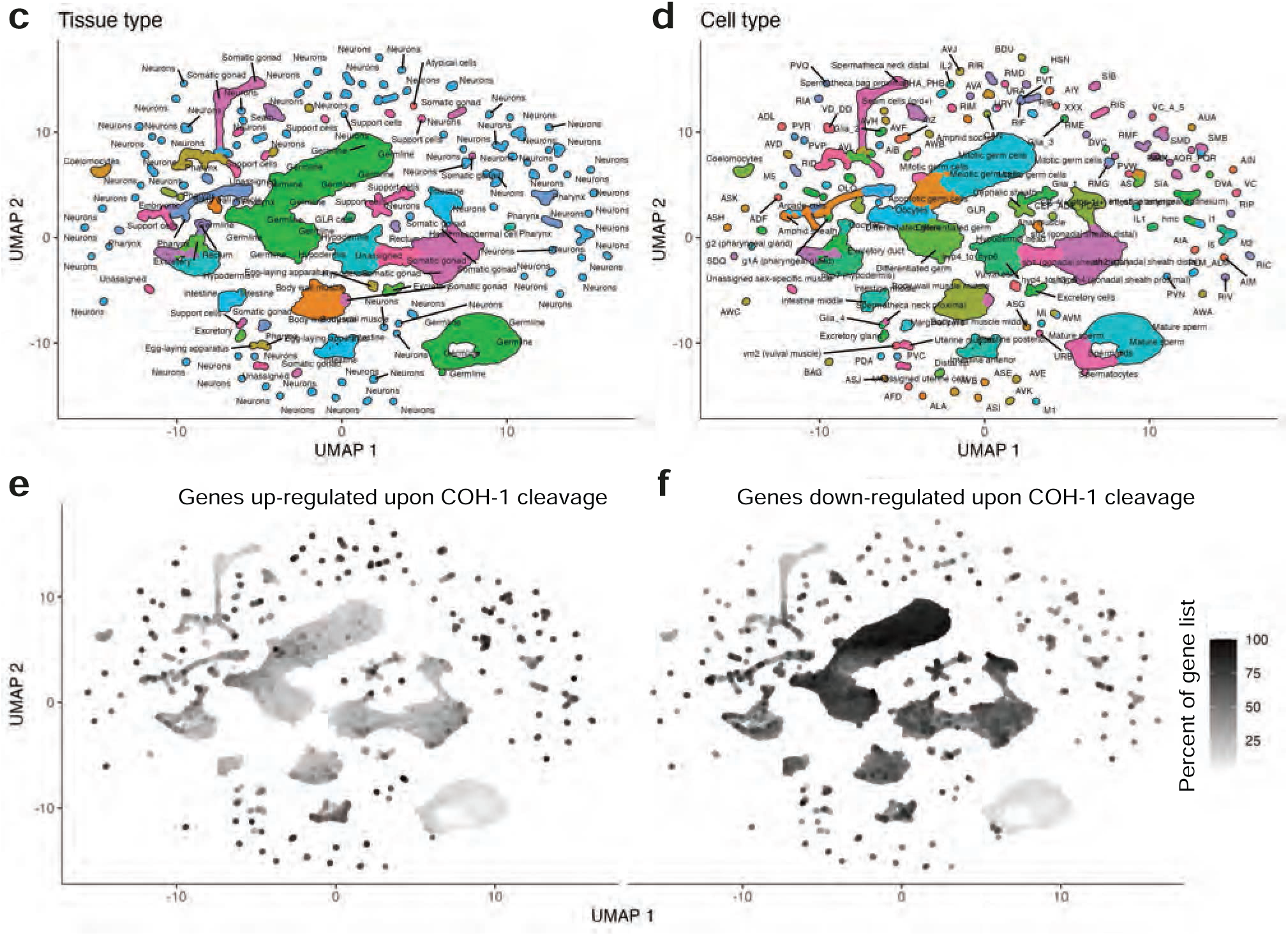
GO term, tissue, cell and phenotype enrichment for genes changing upon COH-1 cleavage. **a.** Tissue Enrichment Analysis for genes significantly up and down regulated upon COH-1^cs^ cleavage (Padj<0.05), using tissues, Gene Ontology and Phenotypes annotations^29^. **b.** WormCat enrichment analysis for genes significantly up and down regulated upon COH-1^cs^ cleavage (adjP<0.05; ref. ^65^). **c.** UMAP visualization of the 180 identified cell types as in ref. ^30^, annotated according to tissue type. **d.** Same map as in c with individual cell types annotated. **e.** Same map as in c with individual cells colored according to the percentage of genes up-regulated upon COH-1 cleavage expressed in those cells (scale in f). **f.** Same map as in c, with individual cells colored according to the percentage of genes down-regulated upon COH-1 cleavage expressed in those cells.

**Supplementary Figure 11.**
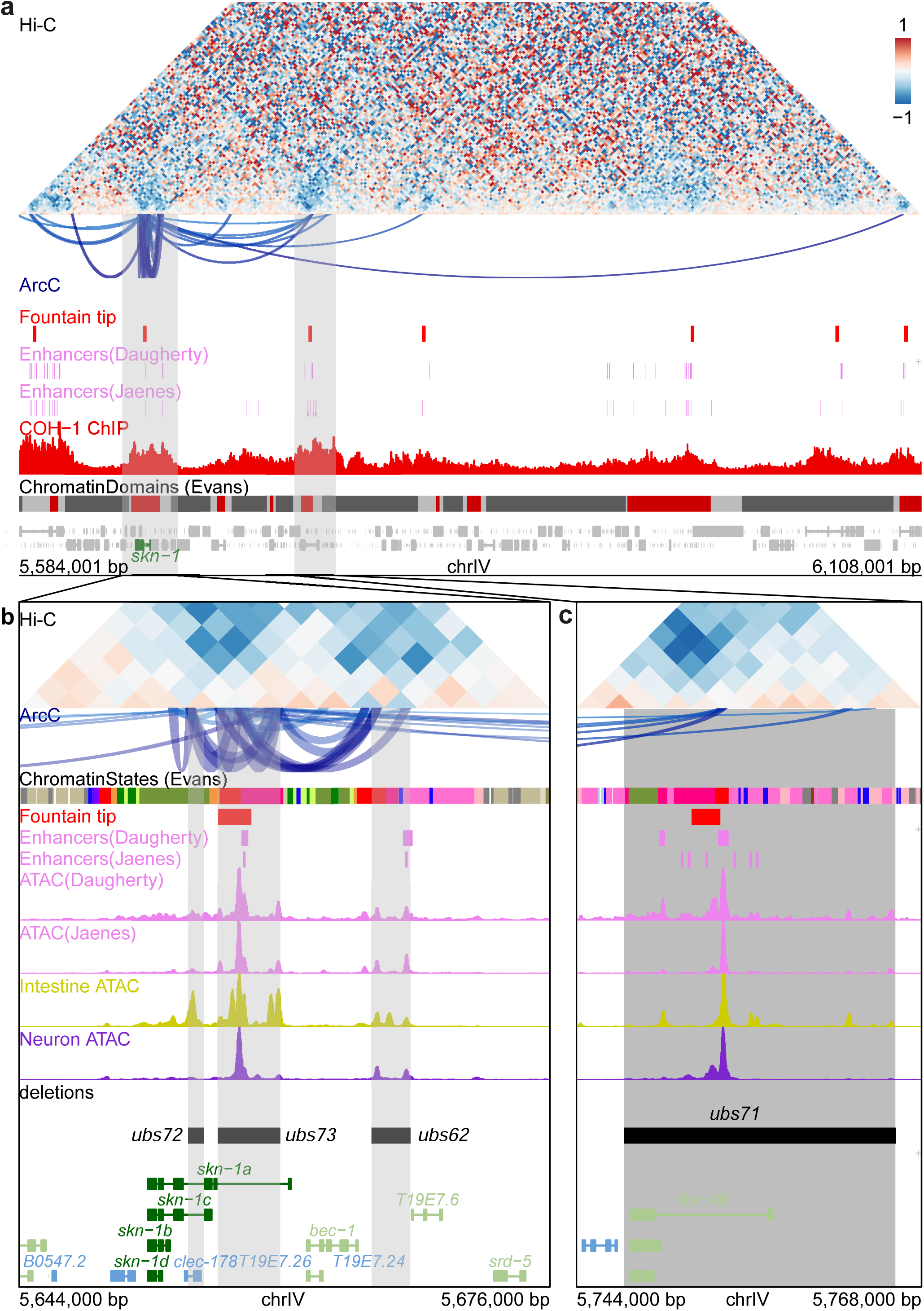
Deletion of putative *skn-1* enhancers. **a.** View of a 524 kb region around the *skn-1* gene (highlighted in green in the gene track). The region was chosen by finding all significant ARC-C cis interactions^8^ with at least one anchor in the *skn-1* gene and its flanking intergenic regions. Grey highlights in the top panel indicate regions shown in detail below: **b.** a 32 kb region around the *skn-1* and *bec-1* genes and **c.** a distal 24 kb region around the *nhr-46* gene. The regions highlighted in the bottom panels correspond to the regions targeted by the deletions. Hi-C: log2 of the ratio of cohesin COH-1 cleavage and TEV control Hi-C maps at 2 kb resolution. Fountain tip: 2 kb bin found at the tip of fountains. Enhancers(Daugherty): L3 active enhancers from ^7^. Enhancers(Jaenes): L3 active enhancers from ^6^. COH-1 ChIP: from young adults (GSE50324). ChromatinDomains (Evans): active (red), regulated (dark grey) and border (light grey) domains as per ^23^. Chromatin states (Evans): see previous reference, red-pink colors indicate enhancer/promoter chromatin states. Intestinal and neuronal ATAC: tissue-specific ATAC in L2 larvae from ^36^. Deletions: CRISPR deletions carried out in this study. The refgene transcript track labeled by gene name is shown on the bottom with transcripts coloured by strand (forward - blue, reverse - green) and *skn-1* transcripts highlighted in dark green.

**Supplementary Figure 12.**
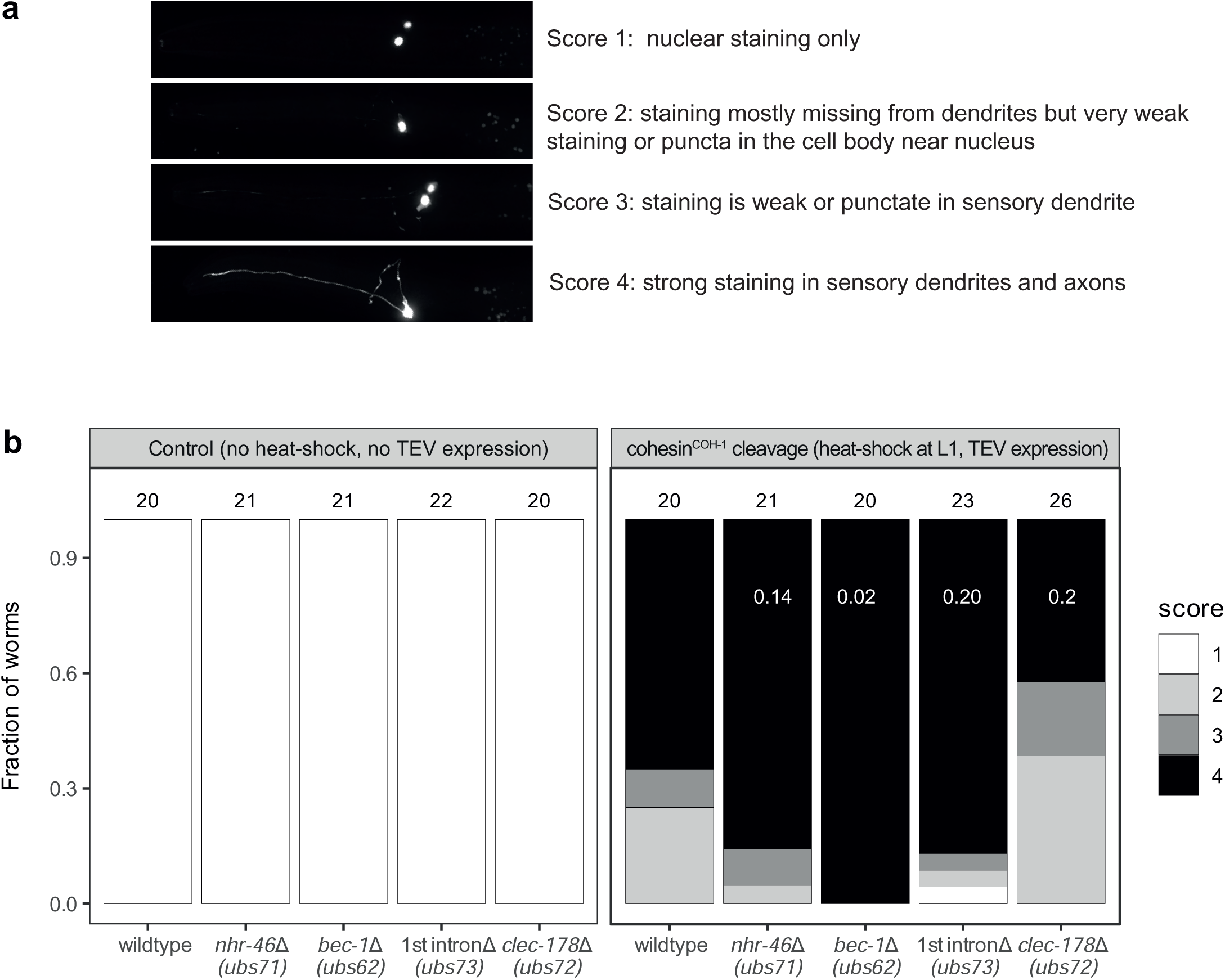
The effect of enhancer deletions on *skn-1::GFP* expression in ASI. **a.** Scoring scheme used to evaluate ectopic expression of *skn-1::GFP* in ASI cells. Images were scored blindly by two separate individuals and then averaged. **b.** Images of the heads of control and COH-1 cleavage worms were scored blindly as described in a. The number of animals scored in each group is shown above the bars. FDR adjusted p-values from an extended Cochran–Armitage test comparing all COH-1 cleavage enhancer deletion strains to the COH-1 cleavage wildtype (without enhancer deletions) strain are shown inside the bars in white.

**Supplementary Figure 13.**
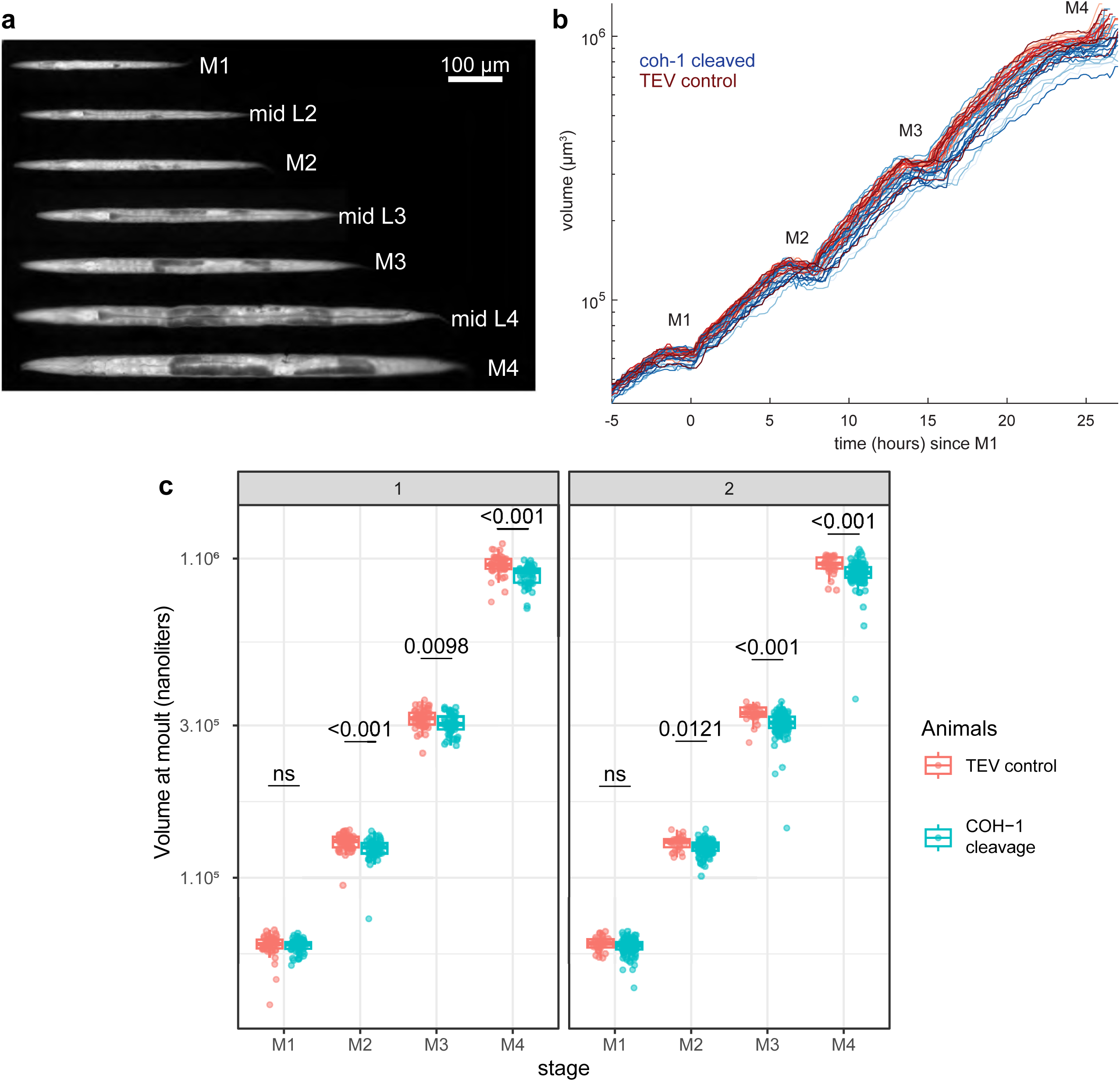
Growth analysis of animals upon TEV control expression or COH-1 cleavage. **a.** An individual animal imaged in a micro chamber at indicated developmental milestones. Contrast was adjusted for each time point individually and the animal straightened computationally. Scale bar: 100 µm. **b.** Example growth curves for 10 control animals and 10 animals upon COH-1 cleavage, synchronized on the first molt. Molts are marked by M. **c.** Body volume at molts for two independent experiments in control animals and upon COH-1 cleavage, n=57;60 (first experiment), 29;122 (second experiment). Box represents 1^st^ quartile, median and 3^rd^ quartile of the data (bottom to top), with individual animals marked as dots. Values above samples are Wilcoxon rank sum test p-values.

**Supplementary Figure 14.**
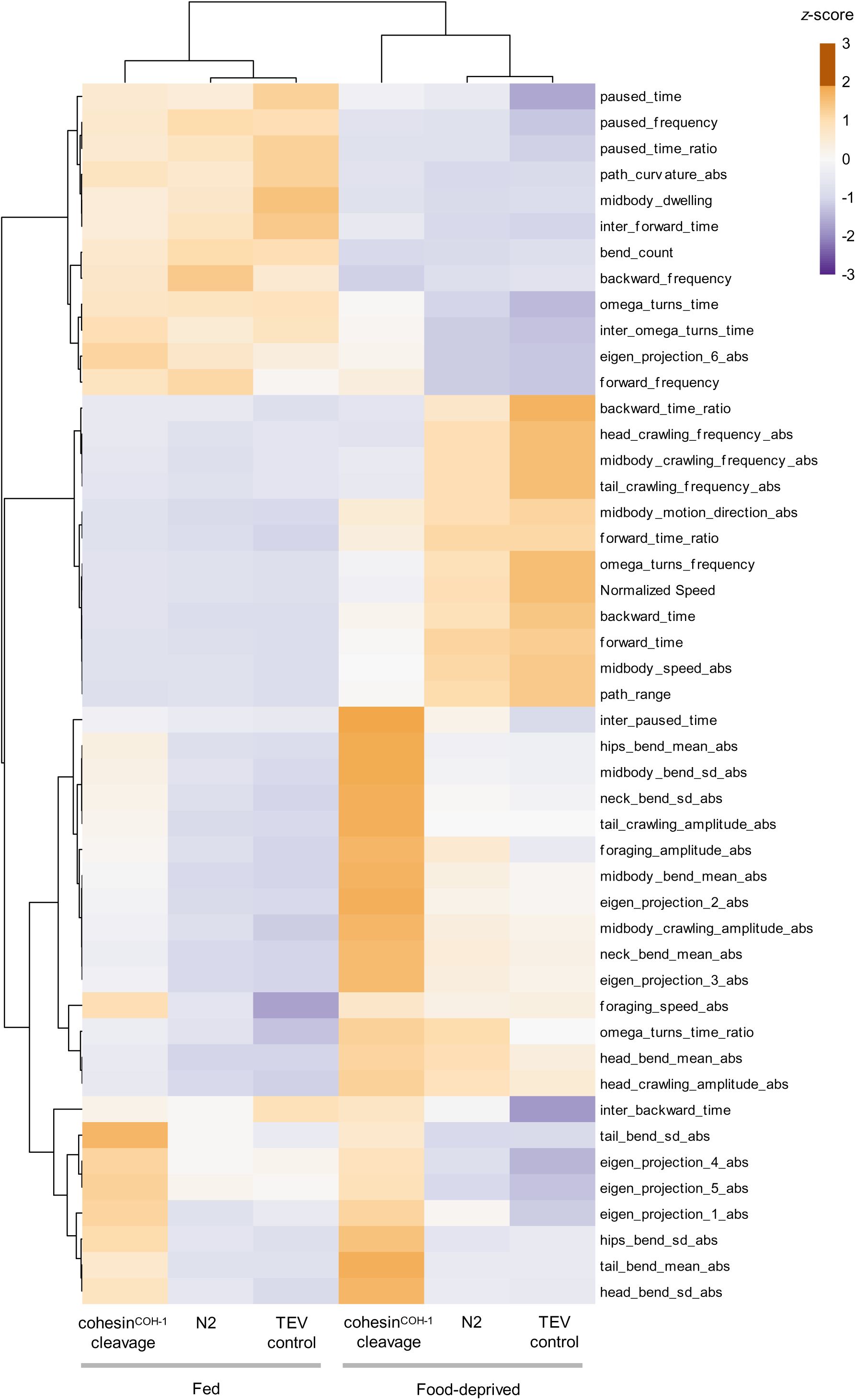
COH-1 cleavage produces broad effects on animal posture and locomotion. Heat-map of behavioral parameters (as z-scores) across the indicated conditions (same treatment as in Fig. 6) and hierarchical clustering based on Euclidean distances (trees). Each data point represents the average value for 3-min recordings on n*=*15 independent replicates (each scoring ≥40 worms).

**Supplementary Figure 15.**
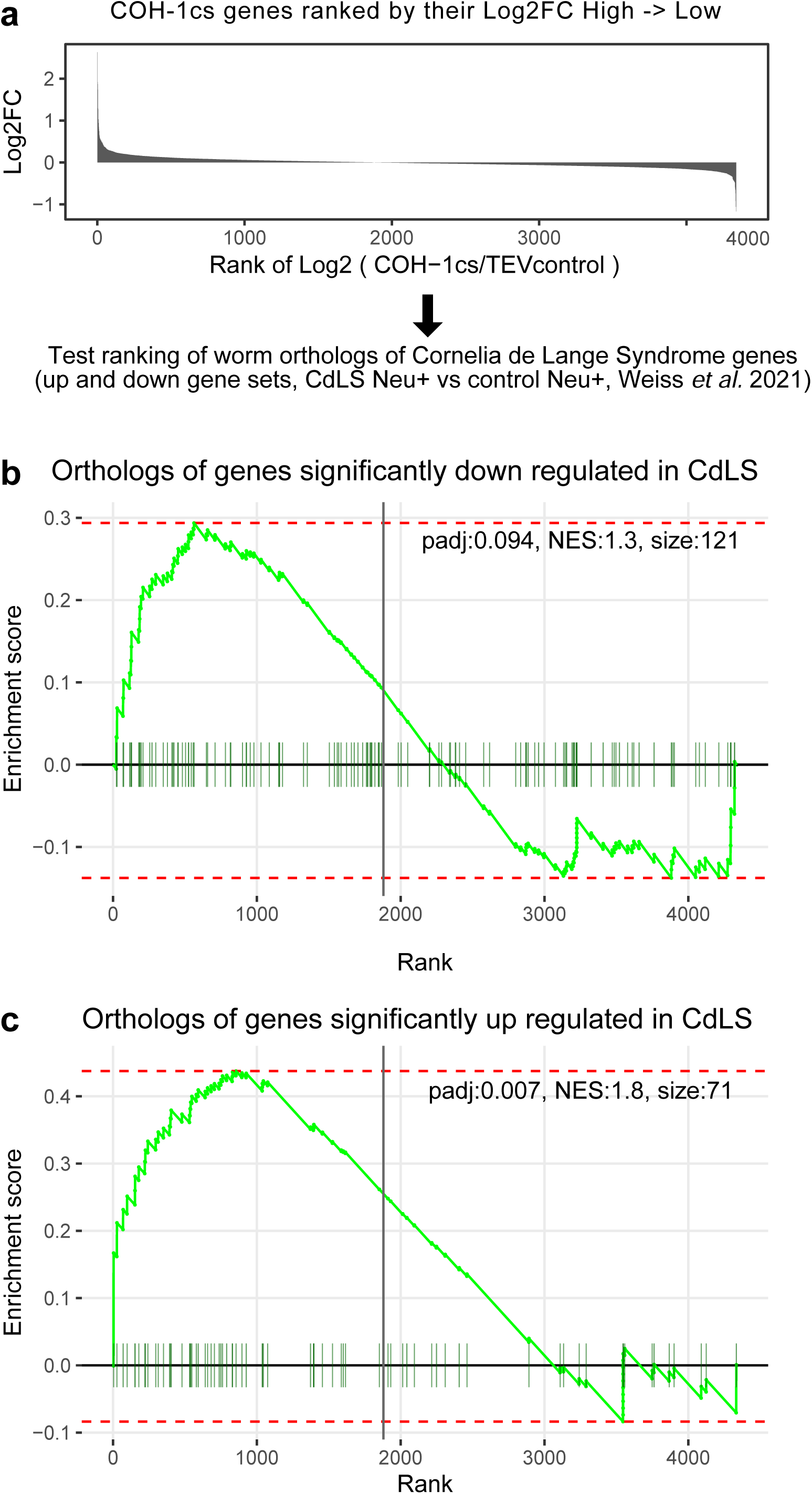
Gene set enrichment analysis (GSEA) of Cornelia de Lange syndrome genes among their nematode orthologs ranked by their log2FC upon COH-1 cleavage. **a.** Schematic of GSEA procedure: genes were ranked in descending order by their log2 fold change upon COH-1 cleavage. The ranks of *C. elegans* orthologs of genes significantly changing in Cornelia de Lange Syndrome (CdLS) patient neurons vs control neurons^63^ were tested for enrichment. **b.** Enrichment of nematode orthologs of genes significantly down regulated in CdLS patient neurons. **c.** Enrichment of nematode orthologs of genes significantly up regulated in CdLS patient neurons. The adjusted p value (padj), normalized enrichment score (NES) and the size of the gene set is shown in each panel on the top right corner. Green ticks indicate the position of the orthologs of CdLS genes among the nematode genes ranked by their log2FC upon COH-1 cleavage. The light green line indicates the enrichment score: a running sum of the degree to which the CdLS genes are enriched at the beginning of the COH-1 cleavage ranked list (positive values) versus the end of the list (negative values).

