## Supplementary files 1-2 for "Cohesin forms fountains at active enhancers in *C. elegans*": Supplementary_file_1_fountain_detection.ipynb

```
# %% [markdown]
# Algorithm for fountain detection in Hi-C map

# %% [code]
#loading useful libraries
%matplotlib widget

import numpy as np
import matplotlib.pyplot as plt
import cooler
import cooltools
import copy
from scipy.special import gammaln
from scipy.signal import find_peaks
from scipy import stats

# %% [markdown]
We based our fountain detection algorithm on a standard method in computer vision: a mask representing an idealized version of the objects or motifs to detect and localize is defined and convoluted to an image such that each pixel of the image obtained a score of object localization. A maxima or peak detection algorithm on these scores then allow to find the pixels where the motifs are localized. For example, this principle is used by other Hi-C-related methods such as ChromoSight (Matthey-Doret et al, Nature Comms 11: 5795, 2020) or Fontanka (Galitsyna et al., bioRxiv 2023).   

# %% [markdown]
### Mask definition

For fountain detection, we defined the mask based on the loop extrusion model and polymer physics in order to focus on motives emerging from loop extrusion by SMCs loaded at specific positions, the fountain origins. 

#### Loop extrusion model
Chromatin is binned at a resolution $r$ (ie one bin=$r$ bp). Let's assume that extruding SMCs are loaded at a site $i_0$ at a rate $k_b$, they extrude chromatin at rates $v_-$ and $v_+$ in "-" and "+" direction respectively and may unbind at a rate $k_u$. We neglect boundaries or excluded volume between cohesins. The number of SMCs $N_{i,j}$ that link bin $i$ ($\le i_0$) to bin $j$ ($\ge i_0$) follows the evolution equation:
\begin{eqnarray}
\frac{d N_{i_0,i_0}}{dt} &=& k_b-(v_-+v_++k_u)N_{i_0,i_0}\\
\frac{d N_{i,j}}{dt} &=& v_- N_{i+1,j}+v_+ N_{i,j-1}-(v_-+v_++k_u)N_{i,j}
\end{eqnarray}
with $N_{i,j}=0$ if $i>i_0$ or $j<i_0$.
At steady-state, this leads to the recursion relation
\begin{eqnarray}
N_{i_0,i_0} &=& \frac{k_b}{v_-+v_++k_u}\\
N_{i,j} &=& \left(\frac{v_- N_{i+1,j}+ v_+ N_{i,j-1}}{v_-+v_++k_u}\right)
\end{eqnarray}
That one can easily solve using the generating function $f(x,y)=\sum N_{i,j} x^{i_0-i} y^{j-i_0}$. This leads to 
\begin{equation}
N_{i,j}=\beta\alpha_-^{i_0-i} \alpha_+^{j-i_0}C_{j-i}^{j-i_0}
\end{equation}
with $\alpha_{\pm}=v_{\pm}/(v_-+v_++k_u)$, $\beta=k_b/(v_-+v_++k_u)$ and $C_n^k$ the combinatorial coefficient of choosing $k$ elements among $n$.

#### Polymer model
Let's assume that at a given time, there is only one loop between $i$ and $j$ ($i\le j$) in the region of interest, and that the polymer dynamics is faster than the loop lifetime. Then the average squared distance between two loci $l$ and $k$ ($l<k$) is given by
\begin{eqnarray}
D^2(l,k|i,j)/b^2 & = & (k-l) \quad\textrm{if $k\le i$ OR $l\ge j$} \\
& = & (i-l)+(k-i)(j-k+1)/(j-i) \quad\textrm{if $l< i$ AND $i<k\le j$} \\
& = & (k-j)+(j-l)(l-i+1)/(j-i) \quad\textrm{if $j>l\ge i$ AND $k> j$} \\
& = & (k-l)(j-i-k+l+1)/(j-i) \quad\textrm{if $i\le l,k\le j$} \\
& = & 1+i-l+k-j\quad\textrm{otherwise}
\end{eqnarray}
with $b$ the size of one bin and where we use the standard expression of squared distance for gaussian linear and ring chains. 
Therefore the probability for bin $l$ and $k$ to be in contact is given by $P_c(l,k|i,j)\approx A/(D^2(l,k|i,j))^{3/2}$ (small radius of contact limit) with $A$ a constant.
Thus the total contact probability, integrating over all the possible loops, is given by 
\begin{equation}
P_c(l,k|i_0)=\frac{1}{Z}\sum_{i\le i_0, j\ge i_0} \frac{\alpha_-^{i_0-i} \alpha_+^{j-i_0}C_{j-i}^{j-i_0}}{(D^2(l,k|i,j))^{3/2}}
\end{equation}
with $Z$ a normalization factor depending on $A$ and $\beta$.

#### Mask
We define the mask as the log ratio of $P_c$ computed above and the probability of contact in a null model without SMC loading
\begin{equation}
F(l,k)=\log\left[\sum_{i\le i_0, j\ge i_0} \frac{\alpha_-^{i_0-i} \alpha_+^{j-i_0}C_{j-i}^{j-i_0}(k-l)^{3/2}}{Z'(D^2(l,k|i,j))^{3/2}}\right]
\end{equation}
with $Z'=\sum_{i\le i_0, j\ge i_0}\alpha_-^{i_0-i} \alpha_+^{j-i_0}C_{j-i}^{j-i_0}=1/(1-\alpha_+-\alpha_-)$.

# %% [code]
#function that compute the mask: 
#2i0+1 is the size of the mask that will be centered around i0 position
#alp: '\alpha_+', alm: '\alpha_-', that control the shape of the mask that become more or less elongated and more or less symmetric.
def filter(i0,alp,alm):
    sfilter=2*i0+1
    F=np.zeros((sfilter,sfilter))
    for i in range(i0+1):
        for j in range(i0,sfilter):
            coef=np.exp((j-i0)*np.log(alp)+(i0-i)*np.log(alm)+gammaln(j-i+1)-gammaln(j-i0+1)-gammaln(i0-i+1))
            for l in range(sfilter):
                for k in range(l+1,sfilter):
                    if (k<=i) or (l>=j):
                        f=k-l
                    elif (l<i) and (k<=j):
                        f=(i-l)+(k-i)*(j-k+1)/(j-i)
                    elif (l>=i) and (k>j):
                        f=(k-j)+(j-l)*(l-i+1)/(j-i)
                    elif (l>=i) and (k<=j):
                        f=(k-l)*(j-i-k+l+1)/(j-i)
                    else:
                        f=1+i-l+k-j
                
                    F[l,k]=F[l,k]+coef*((k-l)/f)**(1.5)
                    F[k,l]=F[l,k]

    F=np.log10(F*(1-alp-alm))
    return F

# %% [code]
# plot some examples of masks
fig,ax=plt.subplots(1,3)
fig.set_figheight(3)
fig.set_figwidth(3*3)
ax[0].matshow(filter(50,0.35,0.35),cmap='bwr')
ax[0].set_title('symmetric mask',fontsize='6')
ax[0].tick_params(axis='x', labelsize=6)
ax[0].tick_params(axis='y', labelsize=6)
ax[1].matshow(filter(50,0.1,0.6),cmap='bwr')
ax[1].set_title('left-handed asymmetric mask',fontsize='6')
ax[1].tick_params(axis='x', labelsize=6)
ax[1].tick_params(axis='y', labelsize=6)
ax[2].matshow(filter(50,0.5,0.2),cmap='bwr')
ax[2].set_title('right-handed asymmetric mask',fontsize='6')
ax[2].tick_params(axis='x', labelsize=6)
ax[2].tick_params(axis='y', labelsize=6)
plt.show()

# %% [markdown]
### Inferring fountain positions

To focus on cohesin-dependent fountains and exclude other motifs that may generate false positive detection or to find fountains that may be hindered inside other strong motifs, we convoluted the mask along the diagonal of the log-ratio matrix of WT HiC on cohesin-depleted HiC. 

# %% [code]
#loading of the Hi-C matrix
c366 = cooler.Cooler('./cooldata/366_merge_2000.cool') #WT data
c828 = cooler.Cooler('./cooldata/828_merge_2000.cool') #cohesin-depleted data

# %% [code]
#compute the log-ratio matrix for a specific chromosome
ch='chrI' #chromosome to consider

mWT=(c366.matrix(balance=True).fetch(ch))
mcoh=(c828.matrix(balance=True).fetch(ch))
mat = np.log10(np.divide(mWT,mcoh))

# %% [code]
#plot WT Hic, cohesin-depleted HiC and log-ratio
fig1 = plt.figure()
fig1.set_figheight(3)
fig1.set_figwidth(3*3)
ax1 = plt.subplot2grid(shape=(1, 3), loc=(0, 0), colspan=1,rowspan=1)
ax2 = plt.subplot2grid(shape=(1, 3), loc=(0, 1), colspan=1,rowspan=1,sharex=ax1,sharey=ax1)
ax3 = plt.subplot2grid(shape=(1, 3), loc=(0, 2), colspan=1,rowspan=1,sharex=ax1,sharey=ax1)
ax1.matshow(np.log10(mWT),cmap='jet',origin='lower')
ax1.set_title('WT Hi-C',fontsize='6')
ax1.tick_params(axis='x', labelsize=6)
ax1.tick_params(axis='y', labelsize=6)
ax2.matshow(np.log10(mcoh),cmap='jet',origin='lower')
ax2.set_title('cohesin-depleted Hi-C',fontsize='6')
ax2.tick_params(axis='x', labelsize=6)
ax2.tick_params(axis='y', labelsize=6)
ax3.matshow(mat,cmap='bwr',origin='lower')
ax3.set_title('log-ratio WT/cohesin-depleted',fontsize='6')
ax3.tick_params(axis='x', labelsize=6)
ax3.tick_params(axis='y', labelsize=6)
plt.show()

# %% [markdown]
First, we estimate an empirical null model that will be used to select statistically-significant fountains. To do so, we generate random matrices bu randomly shuffling the subdiagonals of the matrix to investigate. We apply a mask-based detection method (see Materials and Methods of the main text) to each matrix and estimate the distribution of peak prominences for these matrices taken as an empirical null model.

# %% [code]
#compute a the distribution of local prominences for a null model based on randomizing the subdiagonals
#filter parameter
i0=50 #choosen to be larger than typical fountain size (in bin size)
als=0.35 #=\alpha_0=\alpha_+ = \alpha_-  choosen such that fountains have the expected typical size
F=filter(i0,als,als)

#randomize mat
niter=100 #number of randomization
fs=np.zeros((niter,len(mat))) #fountain score for each randomization
promr=[]
for it in range(niter):
    matr=np.copy(mat)
    for k in range(min(len(matr)-1,2*i0+1)):
        m=mat.diagonal(k).copy()
        np.random.shuffle(m)
        il=np.arange(len(matr)-k)
        matr[il,il+k]=m
        matr[il+k,il]=m
    for i in range(len(matr)):
        d=np.multiply(matr[max(0,(i-i0)):min(len(matr),(i+i0+1)),max(0,(i-i0)):min(len(matr),(i+i0+1))],F[i0-(i-max(0,(i-i0))):i0+(min(len(matr),(i+i0+1))-i),i0-(i-max(0,(i-i0))):i0+(min(len(matr),(i+i0+1))-i)]) #matrix fm
        fs[it,i]=d[np.isfinite(d)].mean()
    peakr,propr=find_peaks(fs[it,:],prominence=(None,None)) #find the peak in fountain score profile
    promr=np.append(promr,propr["prominences"]) #compute the prominences of the peaks

# %% [code]
#compute the threshold to determine significancy score based on the distributions of prominences in the null - randomized - model
fdr=95 #percentage threshold
promth=np.percentile(promr,fdr)

# %% [markdown]
We then apply the mask detection method to the original matrix and select the fountains with a prominence corresponding to an empirical p-value less than 0.05.

# %% [code]
# pass a symmetric mask along the diagonal of the true log-ratio matrix
i0=50 #choosen to be larger than typical fountain size (in bin size)
als=0.35 #=\alpha_0=\alpha_+ = \alpha_-  choosen such that fountains have the expected typical size
F=filter(i0,als,als)

fountainscore=np.zeros(len(mat)) #fountain score S regarding the symmetric mask
semfountain=np.zeros(len(mat)) #standard error on the score
for i in range(len(mat)):
    d=np.multiply(mat[max(0,(i-i0)):min(len(mat),(i+i0+1)),max(0,(i-i0)):min(len(mat),(i+i0+1))],F[i0-(i-max(0,(i-i0))):i0+(min(len(mat),(i+i0+1))-i),i0-(i-max(0,(i-i0))):i0+(min(len(mat),(i+i0+1))-i)]) #matrix fm
    fountainscore[i]=d[np.isfinite(d)].mean()
    semfountain[i]=stats.sem(d[np.isfinite(d)])

# %% [code]
#infer the position of fountain origin
#find the peak in fountain score profiles
peako,prop=find_peaks(fountainscore,prominence=(None,None))
#select only the peaks with prominence larger than the threshold
prom=prop["prominences"]
a=np.where(prom>=promth)[0]
peak=peako[a]
promp=prom[a]

# %% [code]
#visualize the log-ratio and WT maps, the positions of the fountain origins, fountain score profile and the prominence scores for each peak 
fig2 = plt.figure()
fig2.set_figheight(5)
fig2.set_figwidth(8)
x=np.arange(len(mat))
ax2 = plt.subplot2grid(shape=(5, 8), loc=(0, 0), colspan=4,rowspan=1)
ax3 = plt.subplot2grid(shape=(5, 8), loc=(0, 4), colspan=4,rowspan=1,sharex=ax2,sharey=ax2)
ax1 = plt.subplot2grid(shape=(5, 8), loc=(1, 0), colspan=4,rowspan=4,sharex=ax2)
ax4 = plt.subplot2grid(shape=(5, 8), loc=(1, 4), colspan=4,rowspan=4,sharex=ax1,sharey=ax1)
ax1.matshow(mat, cmap='bwr',vmin=-1,vmax=1,origin='lower') #log-ratio map
#ax1.scatter(peak,peak,color="none",edgecolor="black") #approximated position of the fountains
ax4.matshow(np.log10(mWT),cmap='jet',origin='lower') #WT map
ax4.scatter(peak,peak,color="none",edgecolor="black") #approximated position of the fountains
ax2.plot(x,fountainscore,'k') #fountain score profile
ax3.bar(peako,prom,width=8,color='k') #prominence of every peak in the fountain score profile
ax3.bar(peak,promp,width=8,color='r') #significant prominences, ie prominences at detected fountains

ax1.xaxis.set_tick_params(which='both', top=False, labeltop=False,
                             bottom=True, labelbottom=True)
ax4.xaxis.set_tick_params(which='both', top=False, labeltop=False,
                             bottom=True, labelbottom=True)
ax2.axes.get_xaxis().set_visible(False)
ax3.axes.get_xaxis().set_visible(False)
ax3.axes.get_yaxis().set_visible(False)
ax4.axes.get_yaxis().set_visible(False)
plt.show()

# %% [markdown]
### Fountain asymmetry
We compute for each fountain a score for possible asymmetries in fountain shapes.

# %% [code]
#for each peak found, pass several asymetric binarized masks around and conserve the position and height of the peaks
dbin=5 #scanning will be done +/- dbin around the approximated fountain origins inferred above
almin=0.05 #minimal alpha value to be scanned
ft=0.1 #threshold to binarized the mask (choosen such that the number of non-zero entries in the mask is high enough >~ 150 but the difference between symmetric, right-tilted, left-handed and right-handed masks are significant)
alplist=almin+2*(als-almin)/20*np.arange(21) #list of scanned alp values

posm=np.zeros((21,len(peak))) #for each peak and each mask, store the position of the maximal height
heim=np.zeros((21,len(peak))) #for each peak and each mask,store the maximal height
heisem=np.zeros((21,len(peak))) #for each peak and each mask,store the standard error on the maximal height
for k in range(21):
    Fb=filter(i0,alplist[k],2*als-alplist[k]) > ft                
    Fb=Fb/Fb[:].sum()
    for j in range(len(peak)):
        imin=max(0,peak[j]-dbin)
        imax=min(len(mat)-1,peak[j]+dbin)
        jets=np.zeros(imax-imin+1)
        jetsem=np.zeros(imax-imin+1)
        for i in range(imin,imax+1):
            d=np.multiply(mat[max(0,(i-i0)):min(len(mat),(i+i0+1)),max(0,(i-i0)):min(len(mat),(i+i0+1))],Fb[i0-(i-max(0,(i-i0))):i0+(min(len(mat),(i+i0+1))-i),i0-(i-max(0,(i-i0))):i0+(min(len(mat),(i+i0+1))-i)])
            jets[i-imin]=d[np.isfinite(d)].mean()
            jetsem[i-imin]=stats.sem(d[np.isfinite(d)])
        posm[k,j]=imin+np.argmax(jets)
        heim[k,j]=jets.max()
        heisem[k,j]=jetsem[np.argmax(jets)]

# %% [code]
# for each peak, compute the average position on the position whose peak height is "close" (within 2sem) to the max value
asymscore=np.zeros(len(peak)) #asymmetric score: <(alp-als)/als>=average relative value of the difference between the parameter alp-als computed over the alp value where the height is maximal, if the score is >0 ie right-handed SMC loading site (peakpos<peak), if <0: left-handed loading site (peakpos>peak) 
asymscoresem=np.zeros(len(peak)) #standard error of the asymetric score

for j in range(len(peak)):
    m=heim[:,j].max()
    d=np.argmax(heim[:,j])
    a=np.where(m-heim[:,j]<=heisem[d,j])[0]
    asymscore[j]=alplist[a].mean()/als-1
    asymscoresem[j]=stats.sem((alplist[a]/als-1))

# %% [markdown]
### Fountain length
Finally, we estimate a fountain length for each detected one.

# %% [code]
# for each loci, we compute the sum of the ratio matrix in a band of width 2w+1 at a given distance of the main diagonal and parallel to it
w=3 #half band width
d=50 #maximal distance
s=np.zeros((len(mat),d))
ct=np.zeros((len(mat),d))
for a in range(len(mat)):
    i=a
    for j1 in range(d):
        for i1 in range(max(-w,j1-i),min(w+1,len(mat)-1-j1-i)):
            m=mat[i+i1-j1,i+i1+j1]
            if np.isfinite(m):
                s[a,j1]=s[a,j1]+m
                ct[a,j1]=ct[a,j1]+1
            m=mat[i+i1-j1,i+i1+j1+1]
            if np.isfinite(m):
                s[a,j1]=s[a,j1]+m
                ct[a,j1]=ct[a,j1]+1
s=np.divide(s,ct) #vector of sum for each loci

# %% [code]
def fitcurve(param,xdata,ydata):
    [a,b,c,x0,x1,d,e]=param
    return (a+b*(xdata/x0)/(1+(xdata/x0))+c*(xdata/x1)**d/(1+(xdata/x1)**e))-ydata

# %% [code]
#for each fountain and each distance d to the main diagonal, we compute the difference between the average value of s for distance below d and the average value of s beyond d
#the rationale being that this difference should be maximal at a distance corresponding to the size of the fountain
#to facilitate the inference of the maximum we first fit the evolution of the difference as a function of d by a smooth function
#from which it is easy to estimate the position of the maximum
from scipy.optimize import least_squares


lgt=np.zeros(len(peak))

for i in range(1,len(peak)-1):
    v=np.zeros(d)
    for it in range(d):
        v[it]=np.nanmedian(s[peak[i],:(it+1)])-np.nanmedian(s[peak[i],(it+1):])
    
    mc=10000
    mp=np.zeros(7)
    solu=np.zeros(d)
    for it in range(20):
        pfit=least_squares(fitcurve,[-np.random.rand(),1,1,1,it+1,1,1],jac='3-point',method='lm',args=(np.arange(d-10),v[:-10]))
        if pfit.cost<mc:
            mc=pfit.cost
            mp=pfit.x
            solu=(pfit.fun)+v[:-10]
    pk,pp=find_peaks(solu)
    if len(pk)>0:
        lgt[i]=pk[0]
    else:
        lgt[i]=0


# %% [markdown]
### Plot fountains, lengths & asymmetry

# %% [code]
#visualize the log-ratio map, the approximated position of the fountain origins 
fig3 = plt.figure()
fig3.set_figheight(5)
fig3.set_figwidth(8)
x=np.arange(len(mat))
ax2 = plt.subplot2grid(shape=(5, 8), loc=(0, 0), colspan=4,rowspan=1)
ax3 = plt.subplot2grid(shape=(5, 8), loc=(0, 4), colspan=4,rowspan=1,sharex=ax2)
ax1 = plt.subplot2grid(shape=(5, 8), loc=(1, 0), colspan=4,rowspan=4,sharex=ax2)
ax4 = plt.subplot2grid(shape=(5, 8), loc=(1, 4), colspan=4,rowspan=4,sharex=ax1,sharey=ax1)
ax1.matshow(mat, cmap='bwr',vmin=-1,vmax=1,origin='lower') #log-ratio matrix
ax1.scatter(peak,peak,color="none",edgecolor="black")#first scanned peaks (with symmetric mask)
for i in range(len(peak)):
    ax1.plot(peak[i]-np.arange(lgt[i]),peak[i]+np.arange(lgt[i]),color='b')

ax4.matshow(np.log10(mWT),cmap='jet',origin='lower') #WT map
ax4.scatter(peak,peak,color="none",edgecolor="black")#first scanned peaks (with symmetric mask)

ax2.bar(peak,asymscore,width=8,color='k') #asymmetry score at fountains
ax3.bar(peak,promp,width=8,color='r') #symmetry prominence peak

ax1.xaxis.set_tick_params(which='both', top=False, labeltop=False,
                             bottom=True, labelbottom=True)
ax4.xaxis.set_tick_params(which='both', top=False, labeltop=False,
                             bottom=True, labelbottom=True)
ax2.axes.get_xaxis().set_visible(False)
ax3.axes.get_xaxis().set_visible(False)
ax4.axes.get_yaxis().set_visible(False)
ax3.yaxis.tick_right()

plt.show()

# %% [markdown]
### Save data

# %% [code]
#save data in a txt file
file='./fountainposition_'+ch+'_.txt'
doc=open(file,'w')
doc.write("fountain_pos.\t symm._prom._fountain_score\t symm._fountain_score\t SE_sym_f-score\t  Asymetry\t SE_asym. \t length\n")
for j in range(len(peak)):
    doc.write(str(peak[j])+'\t'+str(promp[j])+'\t'+str(heim[10,j])+'\t'+str(heisem[10,j])+'\t'+str(asymscore[j])+'\t'+str(asymscoresem[j])+'\t'+str(lgt[j])+'\n')
doc.close()

# %% [code]
```
